# Uncovering Long Non-Coding RNAs and Exploring Gene Coexpression Patterns in Sorghum Genomics

**DOI:** 10.1101/2024.10.25.620275

**Authors:** Shinde Nikhil, Habeeb Shaikh Mohideen, Raja Natesan Sella

## Abstract

Sorghum (*Sorghum bicolor L. Moench*) is a versatile crop with significant phenotypic and genetic diversity. Despite the availability of multiple resequenced sorghum genomes, noncoding genomic regions remain underexplored. Long noncoding RNA (lncRNA) is a major transcript category with relatively low expression and complex expression patterns that can be identified via RNA-seq. However, it is challenging to distinguish them from protein-coding genes because of their low abundance and tissue-specific expression. In the present study, we employed a dual sorghum reference genome scheme to identify 9467 (31.49% cultivar-specific) and 9551 (28.93% cultivar-specific) lncRNAs in sorghum cultivars BTX642 and RTX430, respectively. Our results showed that a few lncRNAs were linked to pre- and post-flowering drought tolerance in these two genotypes. The NPCTs encoded elements such as transposon-derived Pol polyproteins (RE-1, RE-2, TNT-1), pentatricopeptide repeat proteins, receptor-like protein kinases, zinc finger BED proteins, and putative disease resistance proteins, etc., suggesting their involvement in drought-specific phenotype development. Cis and trans acting targets for identified lncRNAs were predicted. The results showed that upregulated lncRNAs targeted C2H2, B3, putative zinc finger domain, Ras-related protein, L-ascorbate peroxidase-S, lanosterol synthase, and DUF domain genes during drought, which highlights their role in drought tolerance. The downstream gene coexpression analysis revealed time point-specific lncRNA interactions with TFs, resulting in substantial alterations in major TF numbers, including AP2/ERF-ERF, bHLH, bZIP, C2H2, MYB, and NAC. This study provided a more comprehensive understanding of drought tolerance in sorghum by identifying lncRNAs and examining their gene coexpression patterns.

## Introduction

Sorghum *(Sorghum bicolor L. Moench*) is an important food crop that serves as a staple food for over 500 million people worldwide (FAO, 2011). Its primary gene pool consists of 34129 genes with several structural variants (SVs) (McCormick et al. 2018; Cooper et al. 2019). Past studies reported nearly 45000 gene families in sorghum with over 53.69% dispensable gene families, suggesting that sorghum is more genetically diverse than maize (Tao et al. 2021; Nikhil et al. 2024). Despite gene content variation, sorghum reported several SVs, especially deletions that played important roles in local adaptations and plant responses to stresses such as drought (Songsomboon et al. 2021). These included deletions in the MYB/SANT-like DNA-binding domain protein, which is linked to drought tolerance in sugarcane (Salvato et al. 2019). Past studies reported that the overexpression of several genes, such as molybdenum cofactor biosynthesis protein (Lu et al. 2013), expansin (Lü et al. 2013), bZIP (Ying et al. 2012), anthocyanidin 5,3-O-glucosyltransferase (Cui et al. 2017), and the penta-tricopeptide repeat family (Jiang et al. 2015), enhances drought tolerance in plants. However, the regulation of these genes is poorly understood. The loss of function or overexpression of a few genes is insufficient to address plant responses to environmental stimuli and the genetics underlying stress tolerance. A previous analysis reported that long noncoding RNAs (lncRNAs) regulate the expression of several functional genes that confer stress tolerance to plants (Mercer et al. 2009; Xin et al. 2011). Thus, it is important for the growth and development of plants.

Briefly, lncRNAs constitute the major class of noncoding RNAs that impart plants with a wide range of characteristics (Jha et al. 2020). They are >200 nucleotides long, do not encode proteins, and affect gene expression at several levels, including epigenetic, transcriptional, post-transcriptional, translational, and post-translational mechanisms (Panda et al. 2023). Most lncRNAs are tissue and condition-specific and play important roles in physiology and development (Statello et al. 2021). Although these RNAs constitute the majority of noncoding RNAs, their low expression levels are crucial for ensuring specificity to their regulated targets (Mattick et al. 2023). In addition to exhibiting low expression levels, lncRNAs have low sequence conservation and were initially regarded as transcriptional noise (Mattick et al. 2023; Wang et al. 2023). LncRNAs are categorized into five categories based on their position in the genome: sense, antisense, bidirectional, intronic, and intergenic lncRNAs (Wang and Chekanova 2017). They can be polyadenylated or nonpolyadenylated transcripts with low coding potential (Liu et al. 2015a). Previous research has identified a novel non-polyadenylated transcript group in Arabidopsis and rice, which spans 50–300 nt (Liu et al. 2013; Wang et al. 2014). These transcripts were named as intermediate-sized ncRNAs (im-ncRNAs) (Wang et al. 2014). Most of the intergenic regions in Arabidopsis, rice, and maize transcribe lncRNAs that become polyadenylated and are thought to be transcribed by RNA polymerase II (pol II) and some by Pol IV or Pol V, including several unstable lncRNAs (Lu et al. 2012; Ariel et al. 2014; Li et al. 2014). In the widely accepted view, the lncRNAs are stable, RNA polymerase II-transcribed, polyadenylated, capped molecules (Tuck and Tollervey 2013; Ulitsky and Bartel 2013), which can be detected via RNA-seq (Doughty and Kerkhoven 2020).

Given the increasing significance of lncRNAs in stress-responsive gene regulation, this study aimed to identify lncRNAs associated with pre- and post-flowering drought tolerance in sorghum genotypes RTX430 and BTX642, using RNA-seq accessions from previous studies (Varoquaux et al. 2019). The downstream gene coexpression analysis will investigate the functional roles of coexpressed TFs and lncRNAs in the maintenance of this crop during pre- and post-flowering drought.

## Methods

### Data Sources

The raw RNA-Seq accessions of the pre- and post-flowering drought-tolerant sorghum genotypes RTX430 and BTX642 were downloaded from the NCBI Sequence Read Archive (SRA) from past studies (Varoquaux et al. 2019) **(Supplemental Data S1)**. The sorghum reference genome (v3.1), RTX430 cultivar reference (v2.1), and BTX642 cultivar reference (v1.1) were downloaded from Phytozome [https://phytozome-next.jgi.doe.gov]. For conservation analysis, the maize reference genome v5.0 was downloaded from Ensembl Plants [https://plants.ensembl.org], rRNA sequences of sorghum and non-coding gene annotations (*.bed12) of both sorghum and maize were obtained from RNAcentral. (Sweeney et al. 2020).

### Analysis workflow

This analysis was conducted using our in-house pipeline named LncRAnalyzer [https://gitlab.com/nikhilshinde0909/LncRAnalyzer]. To implement it on a dual-reference genome scheme, the first run was conducted using the sorghum reference genome, annotation, rRNA sequences, and raw sequencing reads (*.fastq.gz) files. The second run utilized a cultivar-specific reference genome, annotations, and sequencing reads that remained unmapped to the sorghum reference genome **(Figure 1(a))**. The LncRAnalyzer is comprised of numerous steps, which are discussed below:

#### 1. Data preprocessing

The analysis workflow included quality trimming of raw sequencing reads using FastP (Chen et al. 2018). Then, quality-trimmed reads were aligned to sorghum rRNA sequences using HISAT2 (Kim et al. 2019), followed by sequence alignment map (SAM) to binary alignment map (BAM) conversions and BAM sorting using SAMtools (Li et al. 2009). Finally, the unmapped reads to rRNA sequences were obtained using the *bamToFastq* method of BEDTools (Quinlan and Hall 2010) and compressed using *gzip*.

#### 2. Alignment, reference-guided assembly, and class-code selection

The unmapped reads to rRNA sequences were aligned to sorghum reference genomes using HISAT2, allowing a maximum of 2 mismatches. Reference-guided transcripts from individual *.bam files were assembled in conservative mode with a transcript length >200, default splice junctions, and minimum locus gap separation using StringTie (Pertea et al. 2015). The transcripts *.gtfs from StringTie were merged into a single *.gtf file using the *stringtie--merge* option with minimum transcript length >200, minimum mapped reads >3, and TPM values >1. Furthermore, uniexonic transcripts were discarded from the merged *.gtf file using GFFRead (Pertea and Pertea 2020). Finally, annotation comparison was performed via GFFCompare (Pertea and Pertea 2020), and transcripts belonging to class codes “i”, “o”, “u”, “x”, and “j” were obtained with GFFRead.

#### 3. Identification of lncRNAs and NPCTs

The coding potential, features, and protein domain homologies-based categorization of lncRNA/NPCT transcripts were performed using CPC2, CPAT, RNAsamba, LGC, and PfamScan. CPC2 uses sequence information alone for coding potential-based characterization of putative lncRNA/NPCT transcripts. Subsequently, CPAT was employed to assess coding potential for given set of transcripts using a hexamer table and a logit model, which were re-trained on CDS and known lncRNA sequences of sorghum. RNAsamba is a neural network-based approach that uses mRNAs and known lncRNA sequences to train datasets into HDF5 files. Later, the *rnasamba classify* method was used to estimate their coding potentials. LGC uses sequence information alone to differentiate lncRNAs from protein-coding genes based on ORF length and GC content feature relationships. In addition, PfamScan was used to discard transcripts that exhibited sequence homology with known protein families. This approach omits transcripts that can be translated into protein domains and increases the probability of lncRNA identification.

We utilized FEELnc, a random forest-based approach to report mRNA spliced and intergenic lncRNAs. First, known mRNA transcripts were filtered from lncRNAs using *FEELnc_filter.pl*. Next, coding potential estimation was performed using the reference genome (*.fa), annotations (*.gtf), and merged *.gtf with *FEELnc_codpot.pl*. Here, *FEELnc_codpot.pl* uses two approaches: shuffle which identifies mRNA spliced lncRNAs, and intergenic, which identifies intergenic lncRNAs. Finally, *FEELnc_classifier.pl* was used to assign classes i.e., genic/intergenic, divergent/convergent, sense/antisense, etc., to the reported lncRNA transcripts. The Slncky tool was also used to identify high-confidence lncRNAs by comparing transcripts with annotated features in the genome.

#### 4. lncRNAs and NPCTs outputs

The summary report enumerates the lncRNAs and NPCTs identified by each method in Tab-Separated Values (*.TSV) format. The intersection of all the core methods was taken to obtain true lncRNAs and NPCTs. True lncRNAs were written into *.gtf file, while true NPCTs were written into *.fa. An upset plot describing the intersection of all methods was prepared using the UpSetR (Conway et al. 2017) R package. Additionally, we assigned classes to identified true lncRNAs based on their orientation (sense/antisense), type (intronic/exonic/intergenic), location (upstream/downstream), and distance from protein-coding genes (bp). The biogenesis of identified true lncRNAs from various Transposable Elements (TEs) such as LTR, TIR, SINE, LINE, MITE, and Helitrons was determined by comparing the TE annotations from APTEdb [http://apte.cp.utfpr.edu.br/].

### Downstream Analysis

#### 1. LncRNA conservation analysis

For lncRNA conservation analysis, we constructed LiftOver files by aligning the sorghum reference genome v3.1 and maize reference genome v5.0 using LAST (Frith et al. 2010) followed by construction of chain files using the ucsc-axtChain tool (Karolchik et al. 2004). We used Slncky to report lncRNA orthologs between sorghum and maize. Additionally, the conserved lncRNAs between cultivars were obtained by comparing the genomic coordinates of the lncRNAs using GFFCompare. The class code “=” transcripts were conserved because they had exact exonic overlaps or intron chain matches with the reference. Furthermore, known lncRNAs for sorghum were obtained from PLncDB (Jin et al. 2021), and significant hits (e-value < 10^-5^) were considered conserved lncRNAs.

#### 2. LncRNAs Target Predictions

LncRNA target prediction was performed using a methodology previously described for maize (Liu et al. 2022). Protein-coding genes (PCGs) located 100 kb upstream/downstream of lncRNAs were analyzed to identify cis-targets for lncRNAs. The PCGs that were located 100 kb upstream/downstream of lncRNA and exhibited an expression Pearson’s correlation coefficient (PCC) > |0.6| with a p-value < 0.05 were classified as cis targets.

Triplex-forming oligos (TFOs), i.e., lncRNAs or mi-RNAs, form an RNA–DNA triple helix and regulate the expression of nearby or distant genes via chromatin modifications in eukaryotes (Li et al. 2016; Matveishina et al. 2020). To report lncRNA binding sites in the reference genome, we used triplextor with parameters (minimum guanine rate 50%, score >15, and maximum error rate 5%) (Buske et al. 2012); RNA:DNA triplex sequences located 1000 bp upstream and 200 bp downstream of the transcription start site (TSS) were considered potential target regions for lncRNAs. The PCGs that have lncRNA binding at the promoter region and showed an expression PCC > |0.9| with a p-value < 0.01 were considered as trans targets.

#### 3. Functional annotation of NPCTs

The identified NPCTs were directly searched for homologies with plant protein sequences from SwissProt using BLASTX. Next, the open reading frames (ORFs) of the NPCTs were predicted via TransDecoder with default parameters (Haas and Papanicolaou 2017), followed by homology searches with SwissProt and Pfam using BLASTP and HMMER, respectively. Finally, a functional annotation table was generated from homology searches using Trinotate, as previously described (Ghaffari et al. 2014; Sayadi et al. 2016).

### Time-ordered transcriptomics and gene coexpression studies

#### 1. Transcription factor (TF) prediction

The mRNA sequences from the reference genomes were obtained using GFFRead. TF prediction was performed using the PlantTFCat (Dai et al. 2013), PlantTFDB (Guo et al. 2008), and iTAK (Zheng et al. 2016) web servers. The resulting tables from the above three methods were combined into a single TSV file using R.

#### 2. Differential gene expression and count normalization

The gene and transcript abundance estimates for PCGs and true lncRNAs were performed using featureCounts (Liao et al. 2014) from individual *.bam files. We performed GxT (Group x Timepoints) normalization for genes and lncRNAs using DESeq2 (Love et al. 2014) with the likelihood ratio test (LRT) for individual tissue types. The top 10 most significant lncRNAs were taken and plotted as a line graph for each tissue type. The differentially expressed (P-value < 0.05 and |log2fold| > 2|) lncRNAs (DELs) were retrieved and plotted on a volcano plot via the enhanced volcano plot (Blighe K et al. 2022) R package. The upper quantile normalization and timepoint replication averages of normalized counts were obtained, and TF/lncs and all gene matrices in TSV format were prepared for gene coexpression studies using R.

#### 3. Time-ordered gene coexpression and network visualization

We used the TO-GCN (Chang et al. 2019) package to report coexpressed TFs and lncRNAs in leaf and root tissues of RTX430 and BTX642. For RTX430, we compared control and pre-flowering drought conditions, while for BTX642, we compared control and post-flowering drought conditions. The TF/lncs and all gene matrices obtained in the previous step were used to estimate PCC cutoffs. Next, a gene coexpression network was constructed using the obtained cutoff values, which resulted in eight coexpression types (C1+C2+, C1-C2+, C10C2+,…, C1-C2-). Here, C1 represents condition 1 (e.g., control), and C2 represents condition 2 (e.g., pre-flowering drought for RTX430 or post-flowering drought for BTX642). The “+” and “-” symbol indicates the type of correlation between conditions. Finally, the top 10 highly expressed TFs/lncs at the first time point were used as the seed for the breadth-first search (BFS) to assign time-ordered levels for each condition. The TO-GCN levels for condition C1+C2+ were further mapped against the actual time point expressions using z-score statistics to report changes in the expression of TF/lncs with time, as previously described (Chang et al. 2019). A heatmap showing changes in the expression of TFs/lncs with time was prepared with the *geom_tile* function of the *ggplot2* (Gómez-Rubio 2017) R package. Finally, time-ordered gene coexpression networks were visualized via Cytoscape (Otasek et al. 2019).

### Weighted Gene Coexpression Network Analysis (WGCNA)

We performed WGCNA analysis to obtain coexpression modules associated with pre- and post-flowering drought tolerance in RTX430 and BTX642, respectively. We used DESeq2 normalized counts to perform variance-stabilizing transformation (VST) and filtered genes with higher variability than the third quartile (75%) for WGCNA (Langfelder and Horvath 2008). The third quartile normalized counts were processed using WGCNA to obtain soft threshold (R2 > 0.85) values, module detections, module-trait associations, module eigengenes, and gene significance correlations. The modules showed strong associations with pre-flowering drought for RTX430 and post-flowering drought for BTX642 were selected for downstream analysis.

## Results

### Analysis workflow

#### 1. Data preprocessing

We preprocessed 389 RNA-seq single-end libraries using FastP to remove adapter contamination. Among 8565.40 million reads, 0.34% from both genotypes presented adapter contamination and were removed. Next, unmapped reads with the sorghum rRNA sequences obtained, resulting in a total of 0.66% of reads in both genotypes that aligned with rRNA sequences **(Supplemental Data S2).**

#### 2. Alignment, reference-guided assembly, and class-code selection

Unmapped reads to rRNA sequences were aligned to the sorghum reference genome (v3.1) and presented average alignment rates of 93.37% and 91.77% for BTX642 and RTX430, respectively. The remaining reads were unmapped and further aligned to the cultivar-specific reference genome. This reported average alignment rates of 47.22% and 44.81% for BTX642 and RTX430, respectively **(Supplemental Data S3)**. Reference-guided transcripts were assembled and merged into a single *gtf using StringTie. This reported several transcripts in both genotypes, of which 15.65% were single exonic and discarded. **(Table 1 (a))**. An annotation comparison was performed to retrieve the transcripts belonging to class codes “i”, “o”, “u”, “x”, and “j”. BTX642 reported a total of 50791 and 14088 transcripts that belonged to the desired class codes on the sorghum reference genome and its cultivar-specific reference, respectively. While RTX430 has 51722 and 12582 transcripts that belong to class codes “i”, “o”, “u”, “x”, and “j” on the sorghum reference genome and its cultivar-specific reference, respectively **(Table 1(b)).** We reported a total of 8277 intergenic transcripts in BTX642 and 9251 intergenic transcripts in RTX430 that belonged to class code “u” on both the sorghum and cultivar-specific reference genomes **(Supplemental Figure S1(a)).**

#### 3. Identification of lncRNAs and NPCTs

The class code-selected transcripts were evaluated for coding potentials, features, and sequence homologies with protein domains and classified into lncRNAs and NPCTs. CPAT reported the highest number of lncRNAs in both genotypes, followed by RNAsamba and other approaches. FEELnc identified a total of 14501 mRNAs spliced and 6942 intergenic lncRNAs in BTX642 with the dual reference genome scheme. Similarly, the numbers of mRNAs spliced and intergenic lncRNAs in RTX430 were 14780 and 7448, respectively **(Table 2).** Most of the lncRNAs reported by FEELnc were spliced from mRNAs only, which highlights the role of mRNA splicing in the gene regulation of sorghum. The intersection of all methods is summarized as an upset plot, which shows true lncRNAs and true NPCTs **(Supplemental Figures S1(b)-S1(i)).**

#### 4. lncRNAs and NPCTs outputs

We identified a total of 9467 true lncRNAs in BTX642, which included 6485 (68.50%) on the sorghum reference genome and 2982 (31.50%) on the cultivar-specific reference genome. While RTX430 reported 9551 true lncRNAs, of which 6787 (71.16%) were identified on the sorghum reference genome and 2764 (28.94%) were identified on the cultivar-specific reference genome. The results suggest that the identified lncRNAs in both genotypes vary slightly, and further functional analysis may prove their association with drought tolerance. We assigned classes for identified true lncRNAs and studied their biogenesis from transposable elements (TEs). BTX642 identified 6485 true lncRNAs on the sorghum reference genome, of which 5457 (84.14%) were assigned various classes and 4198 (67.73%) showed biogenesis from TEs. Among these 6485 lncRNAs, the antisense, exonic, and intergenic classes were most prevalent, while LTR-, TIR-, and Helitron-derived lncRNAs were also predominant **(Supplemental Figure S1(j)-S1(k))**. We identified 2982 true lncRNAs in BTX642 on the cultivar-specific reference genome, of which 2734 (91.68%) were assigned to various classes, with antisense exonic and intergenic RNAs being the most abundant **(Supplemental Figure S1(l))**. Similarly, RTX430 identified 6787 lncRNAs on the sorghum reference genome, of which 5603 (82.55%) were assigned classes and 4624 (68.13%) showed biogenesis from TEs. Our results indicated that the classes antisense exonic and intergenic, as well as LTR-, TIR-, and Helitron-derived lncRNAs, were the most prevalent among these 6787 **(Supplemental Figure S1(m)-S1(n))**. RTX430 identified 2764 true lncRNAs on cultivar-specific reference genome, of which 2537 (91.78%) were assigned classes, which suggests antisense exonic and intergenic lncRNAs were most predominant **(Supplemental Figure S1(o))**. The results infer that most of the lncRNAs we identified were transcribed from the opposite strand of coding genes and intergenic regions. Additionally, we reported biogenesis of the most predominant lncRNAs from TEs such as LTR, TIR, and Helitrons. These suggest that they could be valid **(Table 3(a)-3(b); Supplemental Data S4-S5)**.

### Downstream Analysis

#### 1. LncRNA conservation analysis

Our Slncky results showed that out of the total selected transcripts with desired class codes on the sorghum reference genome, 12344 (24.30%) in BTX642 and 117722 (22.76%) in RTX430 were orthologous to maize **(Supplemental Data S6).** These included 784 (12.08%) true lncRNAs in BTX642 and 958 (14.11%) true lncRNAs in RTX430 that were orthologous to maize. Additionally, we identified 600 and 750 long intergenic non-coding RNAs (lincRNAs) in BTX642 and RTX430 that were orthologous to maize **(Supplemental Table S1)**. Results suggest that the number of lncRNA orthologs was marginally higher in RTX430 than in BTX642. Furthermore, we searched for an exact match of the intron chain for identified lncRNAs on the sorghum reference genome in BTX642 and RTX430. This led to the identification of 1528 conserved lncRNAs between these genotypes, indicating that they were conserved and the remaining lncRNAs were novel **(Figure 2(a))**. The results suggest that fewer lncRNAs existed between these two genotypes with exact intron chain matches, and the rest could be spliced from the same gene or transcribed from intergenic regions **(Supplemental Data S6, Sheet 6)**. BTX642 reported a total of 6485 and 2982 lncRNAs in the sorghum reference genome and the cultivar reference genome, respectively. Among them, 2782 (42.90%) and 1055 (35.38%) lncRNAs corresponded to entries in the PLncDB, whereas the rest were novel. Similarly, RTX430 reported 6787 and 2764 lncRNAs in the sorghum and cultivar reference genomes, respectively. Among these, 3286 (48.42%) and 903 (3267%) lncRNAs were found to match in the PLncDB, with the rest being novel. The results suggest that most of the lncRNAs identified using a dual reference scheme were novel and could have an association with drought tolerance in both sorghum genotypes **(Figure 2(b); Supplemental Data S6, Sheet 7)**.

#### 2. LncRNAs Target Predictions

Cis-targets of lncRNAs were reported across tissue types in both genotypes. In BTX642, 469 lncRNAs in leaf tissues were found to be linked with 1329 target genes located 100 kb upstream/downstream. Whereas, 651 lncRNAs in root tissues were associated with 1662 target genes within the same distance. Similarly, RTX430 has 283 lncRNAs in leaf tissues and was found to be linked with 765 target genes located 100 kb upstream/downstream. In the root tissues of RTX430, 537 lncRNAs with 1537 100 kb upstream/downstream target genes were reported **(Table 4(a)).** We found that major TFs, such as C2H2, WD40-like, WRKY, NAC, MADS-MIKC, bZIP, PHD, MYB, GARP-G2-like, bHLH, and B3, were cis targets of those lncRNAs in the leaf and root tissues of BTX642. Similarly, we reported that cis-targets for some lncRNAs, which included TFs: C2H2, bHLH, WRKY, WD40-like, MYB, CCHC(Zn), B3-ARF, AP2/ERF-ERF, bZIP, FAR1, and NAC in the leaf and root tissues of RTX430 ( **Supplemental Data S7)**. To predict lncRNA binding sites, we employed a triplexator, a tool that reports Hoogsteen base pairing with DNA (minimum score = 15 and guanine rate >50%). The promoter region of a gene is defined as the sequence ranging from 1000 bp upstream to 200 bp downstream of the TSS (Yu et al. 2015). Hence, we investigated the formation of DNA:RNA triplexes within this region. In BTX642, Triplexator identified 1148 lncRNAs that formed 14321 triplexes at the promoter regions of 975 genes on the same strand. While 1453 lncRNAs formed 30554 triplexes at the promoter regions of 1759 genes on the opposite strand.

Similarly, RTX430 has 1175 lncRNAs that formed 12424 triplexes at the promoter region of 868 genes on the same strand, and 1466 lncRNAs that formed 28898 triplexes at the promoter region of 1715 genes on the opposite strand **(Table 4(b); Supplemental Data S8).** The given gene was considered a trans-target for lncRNAs when it met the given parameters (PCC > |0.9| and p-value < 0.01) and formed a triplex at the gene’s promoter region. We reported 15 lncRNAs, with 25 and 12 lncRNAs and 19 trans-target genes in the leaf and root tissues of BTX642, respectively. In RTX430, we reported 6 lncRNAs, with 12 trans-target genes and 7 lncRNAs and 17 trans-target genes in leaf and root tissues, respectively. These genes also included several TFs such as C2H2, AP2-ERF/ERF, MYB, bHLH, bZIP, and PAZ-Argonaute targeted by lncRNAs in the leaf and root tissues of both genotypes (**Table 4(c); Supplemental Data S9)**. The results demonstrate that the lncRNA binding sites were higher in BTX642 than in RTX430, which could be correlated with the stay-green phenotype. However, RTX430 has a comparatively smaller number of genes with lncRNA binding sites, which suggests that these regions were either missing or carrying mutations because lncRNAs do not form triplexes.

#### 3. Functional annotation of NPCTs

Among the 476 NPCTs, 418 (87.82%) were annotated using Trinotate in leaf and root tissues of both cultivars, and the rest were unknown protein-coding transcripts. These transcripts were not previously annotated on the sorghum reference or cultivar reference genomes. The NPCTs identified on the sorghum reference genome were considered common; however, the NPCTs exclusively located in the cultivar reference genome were cultivar-specific. Their functional annotation suggested that they encoded elements such as UDP-glycosyltransferase, zinc finger BED domain-containing protein, ribonuclease H protein, receptor-like protein, receptor-like protein kinase, pentatricopeptide repeat-containing protein, putative disease resistance protein, and retrovirus-related Pol polyproteins (RE-1, RE-2, and TNT-1) in both genotypes in varied numbers. Compared with BTX642, the RTX430 reported more NPCTs, most of which were located on the sorghum reference genome, and very few were on the cultivar reference genome. These include some zinc finger proteins, putative disease resistance proteins, putative ribonuclease H, lactate dehydrogenase, cyclic nucleotide-gated ion channels, TNT-1, RE-1, and RE-2, which may be involved in pre-flowering drought tolerance. On the other hand, BTX642 contained NPCTs for a wall-associated receptor kinase, U-box domain-containing protein, polyubiquitin, FAR-Red impaired response protein, and calcium-transporting ATPase. This suggests that these NPCTs might have a hidden role in the drought response and the stay-green phenotype **(Figure 3; Supplemental Data S10)**.

### Time-ordered transcriptomics and gene coexpression studies

#### 1. Transcription factor (TF) prediction

Several transcription factors (TFs) were predicted across the sorghum reference genome and cultivar reference genomes using the iTAK, PlantTFCat, and PlantTFDB online servers. These numbers were 4410, 4217, and 4094 for the sorghum reference genome and the BTX642 and RTX430 cultivar reference genomes, respectively **(Supplemental Data S11).** Among all three reference genomes, PlantTFCat reported the greatest number of predicted TF genes **(Supplemental Table S2).** We found that the sorghum reference genome has more predicted TFs than the cultivar reference genomes, and loss of TF genes might be associated with drought tolerance in these genotypes.

#### 2. Differential gene expression and count normalization

We performed DESeq2 analysis to identify differentially expressed genes, which revealed 17080 genes in leaf tissues and 15754 genes in root tissues of BTX642, showing significant (P-value < 0.05) changes in expression under control, pre-flowering, and post-flowering drought conditions. These included 2834 lncRNAs in leaf and 2894 lncRNAs in root tissues. Of these 2834 lncRNAs in BTX642 leaf tissues, 1299 (45.84%) were upregulated and 1535 (54.16%) were downregulated. Similarly, BTX642 root tissues reported 1341 (46.50%) and 1543 (53.50%) lncRNAs up- and downregulated, respectively. We identified the top 10 most significant lncRNAs in leaf and root tissues. In leaf tissue, 8 of the 10 lncRNAs were downregulated under post-flowering drought conditions compared to the control **(Figure 4(a))**. The remaining two lncRNAs, BTX642.nov01G4214.2 and BTX642.nov10G5336.2, were upregulated under post-flowering drought conditions. We identified target genes for these two lncRNAs. Specifically, BTX642.nov01G4214.2 positively regulates C2H2-110, Ras-related protein (RAB-18), Methyltransferase (PMT-14), Domain of Unknown Function (DUF-677), and other putative genes, while BTX642.nov10G5336.2 positively regulates a putative zinc finger domain and negatively regulates electron carrier/ protein disulfide oxidoreductase genes. In the root tissues of BTX642, all 10 of the most significant lncRNAs were downregulated under post-flowering drought conditions compared to the control **(Figure 4(b))**. The results showed that in BTX642 leaf and root tissues, most of the lncRNAs were downregulated under post-flowering drought conditions, which could be associated with the stay-green phenotype in BTX642. However, we found that the lncRNAs BTX642.nov01G4214.2 and BTX642.nov10G5336.2 were upregulated in leaf tissues under post-flowering drought, suggesting a potential role in the stay green phenotype in BTX642.

In RTX430, we reported 19237 genes in leaf and 16309 genes in root tissues were differentially expressed (P-value < 0.05). These included 3021 lncRNAs in leaf and 2782 lncRNAs in root tissues. Of these 3021 lncRNAs in leaf tissues, 1420 (45.50%) were upregulated and 1701 (54.50%) were downregulated. In the root tissues of RTX430, we found 1374 (49.21%) lncRNAs were upregulated and 1418 (50.78%) lncRNAs were downregulated. We listed the top 10 most significant lncRNAs, and of these, 8 lncRNAs exhibited lower expression levels in leaf tissue under pre-flowering drought compared to the control. However, their expression levels increased when watering was restored **(Figure 4(c))**. The remaining two lncRNAs, RTX430.nov02G7151.1 and RTX430.nov06G17608.1, were upregulated under pre-flowering drought compared to the control. We identified target genes for these lncRNAs, which showed that RTX430.nov06G17608.1 positively regulates the L-Ascorbate Peroxidase-S (APX-S) gene, while RTX430.nov02G7151.1 positively regulates genes such as B3-20, DUF-4378 and Lanosterol Synthase (LS) genes. These results suggest that the upregulation of genes like APX-S, B3-20, LS, and DUF-4378 in leaf tissues of RTX430 could be associated with pre-flowering drought tolerance. In root tissues, we found that all of the top 10 significant lncRNAs were downregulated under pre-flowering drought conditions compared to the control, and their expression levels increased when watering was restored. This suggests that these lncRNAs could be associated with the recovery of RTX430 from pre-flowering drought **(Figure 4(d); Supplemental Table S3(a), Supplemental Data S12)**.

We compared control versus pre- and post-flowering drought conditions for BTX642 and RTX430 to identify differentially expressed lncRNAs (DELs) (P-value < 0.05) with significant log2fold change (| log2fold| > 2) in these genotypes. In BTX642 leaf tissues, we identified 499 DELs, of which 197 showed significant upregulation (log2fold > 2) and 302 showed a significant downregulation (log2fold < −2) under post-flowering drought. Similarly, pre-flowering drought identified 177 DELs, of which 72 were significantly upregulated and 105 remained significantly downregulated **(Supplemental Figure S4(a))**. These results infer that BTX642 leaf tissues reported more DEL numbers with significant up- and downregulation under post-flowering drought than pre-flowering drought conditions, which could be correlated with the stay green phenotype. In the BTX642 root tissues, we found 383 DELs, of which 130 and 253 showed significant up- and downregulation, respectively, under post-flowering drought conditions. We reported 359 DELs under pre-flowering drought conditions, of which 231 were significantly upregulated and 128 remained significantly downregulated **(Supplemental Figure S4(b))**. The results suggested that the number of DELs with significant up- and downregulation in the BTX642 root tissues differs between drought stages, and the number of downregulated lncRNAs under post-flowering drought could be associated with the stay green phenotype.

In RTX430 leaf tissues, we found only 50 DELs, of which 30 showed significant upregulation and the remaining 20 showed significant downregulation under pre-flowering drought conditions. While 315 DELs were identified under post-flowering drought, of which 78 were significantly upregulated and 237 remained significantly downregulated **(Supplemental Figure S4(c))**. The results suggest that reduced DELs in leaf tissues of RTX430 could be associated with pre-flowering drought tolerance. In the root tissues of RTX430, we found 323 under pre-flowering drought, of which 212 were significantly upregulated and 111 were significantly downregulated. Similarly, 289 DELs were identified under post-flowering drought conditions, of which 111 showed significant upregulation and the remaining 178 showed significant downregulation **(Supplemental Figure S4(d))**. The results indicate that the root tissues of RTX430 exhibit upregulation of 212 DELs under pre-flowering drought, which could be associated with its tolerance to pre-flowering drought **(Supplemental Table S3(b))**.

#### 3. Time-ordered gene coexpression and network visualization

To report coexpressed genes, we performed a TO-GCN analysis comparing control conditions with pre- and post-flowering drought for RTX430 and BTX642, respectively. In BTX642 leaf tissues, we identified 12 coexpression levels with varying numbers of TFs and lncRNAs for both control and post-flowering drought conditions **(Figure 5(a))**. These included 471 lncRNAs and 111 distinct transcription factors (TFs) that were coexpressed under the control condition. Major TFs in this category included C2H2 (145), AP2-ERF/ERF (58), WRKY (54), bHLH (53), MYB (48), WD40-like (41), PHD (37), bZIP (30), and others. We reported 431 lncRNAs, and 100 distinct TFs were coexpressed under post-flowering drought conditions, including C2H2 (124), AP2-ERF/ERF (58), WRKY (49), bHLH (48), MYB (30), WD40-like (43), PHD (28), bZIP (24), and others **(Figure 5(b))**. The results suggest that the coexpressed TFs and lncRNAs were decreased under post-flowering drought conditions in the BTX642 leaf tissues, which may be associated with the stay-green phenotype. In particular, the numbers of C2H2, MYB and lncRNA were significantly reduced under post-flowering drought conditions compared to the control, indicating their potential involvement in post-flowering drought tolerance **(Supplemental Data S12, sheet 2)**. In root tissues, we reported 13 coexpression levels for control and only 7 coexpression levels for post-flowering drought conditions **(Figure 5(c))**. These comprised 325 lncRNAs that were coexpressed with 88 distinct TFs under control conditions and 153 lncRNAs that were coexpressed with 70 different TFs in post-flowering drought conditions. We listed the top TFs (control/post-flowering drought) involved in the TO-GCN network, which included C2H2 (143/73), bHLH (54/48), AP2/ERF-ERF (45/40), WRKY (45/23), MYB (42/29), bZIP (34/10), NAC-like (32/23), and others **(Figure 5(d); Supplemental Data S13, sheet 3)**. The results suggest that the BTX642 root tissues exhibited reduced major TFs and lncRNA numbers under post-flowering drought, which may be associated with post-flowering drought tolerance. Especially, TFs such as C2H2, WRKY, MYB, bZIP, and NAC numbers were reduced under post-flowering drought, highlighting the role of these TFs and lncRNAs in post-flowering drought regulation.

For RTX430 leaf tissues, we identified 489 lncRNAs coexpressed with 95 different TFs in 14 levels under control conditions and 508 lncRNAs coexpressed with 94 distinct TFs in 8 levels under pre-flowering drought conditions **(Figure 5(e))**. These included major TFs (control/pre-flowering drought) such as C2H2 (164/167), AP2/ERF-ERF (65/73), WRKY (59/52), bHLH (55/57), WD40-like (46/44), MYB (44/41), NAC (43/46), and others **(Figure 5(f); Supplemental Data S13, sheet 4)**. Results infer that pre-flowering drought conditions showed slightly higher lncRNA and C2H2, AP2/ERF-ERF, and NAC numbers in leaf tissues, which could be related to pre-flowering drought tolerance and recovery in RTX430. In the root tissues of RTX430, we identified 14 coexpression levels for control and 12 coexpression levels for pre-flowering drought conditions **(Figure 5g))**. These included 362 lncRNAs coexpressed with 88 different TFs under control conditions and 339 lncRNAs coexpressed with 83 different TFs under pre-flowering drought conditions. We found major TFs (control/pre-flowering drought), including C2H2 (135/141), WRKY (50/60), AP2/ERF-ERF (67/57), bHLH (58/49), MYB (46/49), NAC (46/47), bZIP (37/33), PHD (35/32), and others that were coexpressed in root tissues **(Figure 5(h); Supplemental Data S13, sheet 5)**. In particular, select lncRNAs, C2H2, and WRKY numbers were slightly higher in pre-flowering drought conditions than in control conditions, highlighting their involvement in pre-flowering drought tolerance and recovery.

### Weighted Gene Coexpression Network Analysis (WGCNA)

We performed WGCNA analysis to detect potential coexpression modules associated with pre- and post-flowering drought tolerance in RTX430 and BTX642, respectively. BTX642 identified 15 coexpression modules in leaf tissues **(Figure 6a))**. We found that coexpression modules “Black”, “Magenta”, and “Tan” were associated with post-flowering drought conditions when performing module-trait relationship and module eigengenes analysis **(Figure 6(b); Supplemental Figure S6(b))**. Furthermore, we found that “Black”, “Magenta”, and “Tan” included several coexpressed lncRNAs and genes **(Supplemental Table S4)**. Gene Ontology (GO) enrichment analysis of these modules shows that they were associated with small molecule metabolic process, oxoacid metabolic process, cellular lipid metabolic process, phosphorus metabolic process, etc **(Supplemental Figure S6(c))**. Similarly, BTX642 root tissues reported 14 coexpression modules **(Figure 6c))**, of which four coexpression modules, namely “Green”, “Magenta”, “Pink”, and “Salmon”, were associated with post-flowering drought tolerance **(Figure 6(d); Supplemental Figure S6(e))**. The GO enrichment analysis revealed that those genes were involved in transport, membrane, multicellular organism development, developmental process, carbohydrate metabolic process, and anatomical structure development **(Supplemental Figure S6(f))**. In RTX430 leaf tissues, we reported 20 coexpression modules **(Figure 6e))**. Out of the 20 coexpression modules in RTX430 leaf tissues, four modules, “Green”, “Grey”, “Magenta”, and “Lightcyan”, exhibited a substantial correlation with the pre-flowering drought response **(Figure 6(f); Supplementary Figure S6(h))**. The GO enrichment analysis of these four modules suggests that they contained genes involved in photosystem II, photosynthetic membrane, photosynthesis, light harvesting, heat shock protein binding, hydrolase activity, and electron transfer activity **(Supplemental Figure S6(i))**. Similarly, we identified 11 compression modules in RTX430 root tissues **(Figure 6g))**. We selected four modules, “Black”, “Blue”, “Pink”, and “Red” from the 11 available, based on their association with pre-flowering drought **(Figure 6(h); Supplemental Figure S6(k))**. Our GO enrichment analysis reported that they were involved in xenobiotic transmembrane transport, signaling, microtubule motor activity, oxidoreductase activity, peroxidase activity, phospholipid metabolic process, glycosyltransferase activity, hydrolase activity, carbohydrate metabolic process, and antioxidant activity **(Supplemental Figure S6(l); Supplemental Data S14)**.

## Discussion

The present analysis used 389 public RNA-Seq datasets from leaf and root tissues of two sorghum genotypes to investigate the role of lncRNAs in regulating responses to pre- and post-flowering drought conditions. We identified known and cultivar-specific lncRNAs in sorghum using a dual-reference genome scheme. In BTX642, we identified a total of 9467 lncRNAs using LncRAnalyzer, of which 2982 (31.49%) were cultivar-specific. While RTX430 has a total of 9551 lncRNAs, of which 2764 (28.93%) were cultivar-specific. Results suggest that cultivar-specific lncRNA numbers were slightly higher in BTX642, and could be associated with post-flowering drought tolerance. Past studies reported that some cultivar-specific lncRNAs were associated with heat-tolerant stress tolerance in wheat (Babaei et al. 2024).

We performed lncRNA conservation analysis using Slncky, which showed that 12344 and 11772 class code selected transcripts were orthologous with maize in BTX642 and RTX430, respectively. They included 784 and 958 conserved lncRNAs in BTX642 and RTX430. Results inferred that the RTX430 has comparatively higher numbers of conserved lncRNAs with maize than BTX642. Previous research indicated that 21943 sorghum transcript isoforms, including non-coding genes, exhibited conservation with maize (Wang et al. 2018). We compared the genomic coordinates of lncRNAs identified in BTX642 and RTX430 with the sorghum reference genome. Our analysis revealed that 1528 lncRNAs shared identical intron chains, indicating conserved structures across both cultivars. The remaining lncRNAs were classified as novel. We found that 45.72% of the lncRNAs in the sorghum reference genome and 34.08% of the lncRNAs in the cultivar reference genome shared similarities with PLncDB sequences in both genotypes. This implies that many cultivar-specific lncRNAs in both genotypes were novel and might be induced by drought response. A previous study on lncRNA conservation using BLASTN reported that lncRNAs exhibit lower sequence conservation than the protein-coding genes (Sang et al. 2021). Similar results for lncRNA conservation were previously reported in many plant species, which supports our findings (Hezroni et al. 2015; Ulitsky 2016; Deng et al. 2018). The lower conservation of lncRNAs might be due to extensive mRNA splicing and transcription of intergenic regions. Previous research has demonstrated that lncRNAs are products and precursors of alternative splicing in cancer, influencing disease progression through cancer-related splice variants (Ouyang et al. 2022).

We reported several lncRNAs that were cis- and trans-acting targets for several TFs, such as AP2/ERF, NAC, MYB, WRKY, bZIP, bHLH, B3-ARF, C2CH2, GRAS, etc. Previous studies reported that these TFs were associated with the drought stress response in plants (Hu et al. 2022). Additionally, we identified several lncRNAs forming triplex structures within 1000 bp upstream and 200 bp downstream of the transcription start sites (TSSs) of key TFs, including A20-like, bHLH, bZIP, C2H2, and MYB. These findings underscore the critical role of lncRNAs in regulating TF activity. However, the number of triplex-forming sites was reduced in RTX430, suggesting that the mutations and genome organization of RTX430 might influence triplex formation. Research has shown that lncRNA:DNA triplexes form specific chromosomal regions and are essential for predicting topologically associating domains (TADs) (Soibam and Zhamangaraeva 2021). Therefore, lncRNA:DNA triplexes affect both gene regulation and the organization of three-dimensional chromatin architecture.

A total of 476 NPCTs were reported in the two genotypes, and these sequences were transcribed and not previously annotated in the genomes. Most of these genes encode elements such as UDP-glycosyltransferase, zinc finger BED domain-containing protein, ribonuclease H protein, receptor-like protein, receptor-like protein kinase, pentatricopeptide repeat-containing protein, putative disease resistance protein, and retrovirus-related Pol polyproteins (RE-1, RE-2, and TNT-1). Our results suggested that their expression is associated with the drought stress response in sorghum. Previous analyses revealed that TNT-1 expression is associated with abiotic factors such as wounding, freezing, and others, and is known to induce plant defense mechanisms (Mhiri et al. 1997). Moreover, the activity of transposable elements (TEs) is influenced by stress and leads to increased tolerance in plants (Thieme et al. 2022). These supported our findings, suggesting that the relationship between transposase activity and drought tolerance in sorghum can be further explored.

We found that the lncRNAs BTX642.nov01G4214.2 and BTX642.nov10G5336.2 were upregulated under post-flowering drought conditions in BTX642 leaf tissues. These lncRNAs regulate C2H2-110, RAB-18, DUF-677, putative zinc finger domain, protein disulfide oxidoreductase, and other putative genes, suggesting their involvement in post-flowering drought tolerance. Previous studies reported that the C2H2 zinc finger protein increased drought tolerance in plants (Han et al. 2020; Bouard and Houde 2022; Chen et al. 2024b). Additionally, stomatal closure in plant leaves is regulated by Ras-related small GTP-binding proteins through ABA signaling, which improves their drought tolerance (Ambastha et al. 2021; Chen et al. 2021). Past analysis reported DUF proteins were highly conserved in eukaryotes and participate in abscisic acid and cytokinin pathways, which enhance drought tolerance in rice (Chen et al. 2023; Jayaraman et al. 2023; Yu et al. 2024). In the roots of BTX642, we found that most lncRNAs were downregulated under post-flowering drought, which could be correlated with the stay-green phenotype. RTX430 leaf tissues reported two major candidates, RTX430.nov02G7151.1 and RTX430.nov06G17608.1, which showed upregulation under pre-flowering drought conditions. These lncRNAs positively regulate B3-20, APX-S, LS, and DUF-4378 genes. Previous research has suggested that B3 domain proteins enhance the drought tolerance of plants by altering stomatal shape and reducing stomatal density in leaves (Liu et al. 2015b; Verma and Bhatia 2019; Wang et al. 2024). In plant species, ascorbate peroxidase (APX) and glutathione peroxidases (GPX) are major ROS-scavenging enzymes responsible for H2O2 reduction and preventing potential cellular damage resulting from H2O2 (Ozyigit et al. 2016). LS is a sterol molecule that acts as a structural component of plasma membranes and precursor of steroidal hormones (Suzuki et al. 2006). A past study reported that the putative LS encoding gene was upregulated in the leaf tissues of *Atractylodes chinensis* under drought stress (Ma et al. 2024). These emphasized the significance of lncRNA-regulated genes in the drought stress responses. In RTX430 root tissues, the most significant 10 lncRNAs were downregulated under pre-flowering drought. However, 12 DELs exhibited significant upregulation under pre-flowering drought in RTX430 root tissues, which could be correlated with pre-flowering drought tolerance.

Time-ordered transcriptomics of leaf and root tissues of BTX642 and RTX430 revealed time point coexpression of TFs and lncRNAs at different levels. In BTX642 leaf tissues, we found that coexpressed lncRNAs and TFs were reduced, with particular emphasis on C2H2 under post-flowering drought. This observation could be related to post-flowering drought tolerance in BTX642. In previous research, it was determined that the C2H2 zinc finger domain negatively regulates drought tolerance by increasing the ROS accumulation in plant leaves (Yang et al. 2021). In BTX642 root tissue, we observed that coexpressed lncRNA and C2H2, WRKY, MYB, bZIP, and NAC numbers were significantly decreased under post-flowering drought. This highlights their involvement in the stay-green phenotype in BTX642. Past studies reported that C2H2 zinc finger candidates negatively regulate drought tolerance in Arabidopsis (Tian et al. 2024) and maize (Liu et al. 2024). In addition to this, some bZIP (Liu et al. 2012), MYB (Li et al. 2020; Peng et al. 2023), NAC (Li et al. 2023; Fan et al. 2024), and WRKY (Ahammed et al. 2020; Jia et al. 2024) TFs were found to be negatively regulating drought tolerance in plants, which supports our findings. In RTX430 leaf tissues, we found slightly higher coexpressed lncRNA, C2H2, AP2/ERF-ERF, and NAC numbers under pre-flowering drought than control, which could be correlated with pre-flowering drought tolerance and recovery. As previously reported, the overexpression of the C2H2 zinc finger domain enhances rice’s drought tolerance by facilitating the breakdown of ABA (Wang et al. 2022). Moreover, overexpression of selected members of C2H2 (Cheuk et al. 2020; Bouard and Houde 2022; Chen et al. 2024b), AP2/ERF (Jisha et al. 2015; Kong et al. 2023), and NAC (Hu et al. 2006; Zheng et al. 2009; Hong et al. 2016) also enhanced drought tolerance in plants. In the root tissues of RTX430, we identified more coexpressed lncRNAs and TF numbers under pre-flowering drought, with a particular emphasis on C2H2 and WRKY. Previous research demonstrated that drought stress tolerance in plants was improved by overexpressing WRKY candidates in wheat (Gao et al. 2018), tomato (Chen et al. 2024a), and Arabidopsis (Zhang et al. 2022; Bai et al. 2023). Our TO-GCN analysis examined the functional relationships between lncRNA and TF. The results indicated that post-flowering drought tolerance in BTX642 is negatively correlated with certain lncRNAs, as well as C2H2, WRKY, MYB, bZIP, and NAC members. Conversely, pre-flowering drought tolerance in RTX430 is positively correlated with certain lncRNAs and specific members of C2H2, AP2/ERF-ERF, NAC, and WRKY. Our WGCNA analysis identified numerous modules linked to drought tolerance. They identified genes that are involved in transport, membrane, electron transfer, carbohydrate metabolism, photosynthesis, and heat shock in leaf tissues. However, root tissues contained genes that were involved in carbohydrate metabolism, antioxidant activity, signaling, microtubule motor activity, oxidoreductase activity, and peroxidase activity.

## Conclusion

The present study identified lncRNA and NPCTs using public RNA-Seq datasets of sorghum BTX642 and RTX430 under control, pre-, and post-flowering drought conditions. Using a dual reference genome scheme, we reported 9467 (3349% cultivar-specific) and 9551 (28.93% cultivar-specific) lncRNAs in BTX642 and RTX430, respectively. The expression patterns of the top 10 most significant lncRNAs in both genotypes illustrated that drought stress upregulated only two candidates. These lncRNAs were positive regulators of B3-20, C2H2-110, RAB-18, DUF-677, APX-S, LS, and DUF-4378 in BTX642 and RTX430, highlighting their role in drought stress response. With gene co-expression analysis, we found that some lncRNAs, along with C2H2, WRKY, MYB, bZIP, and NAC members, showed a negative association with post-flowering drought in BTX642. While some lncRNAs, along with C2H2, AP2/ERF-ERF, NAC, and WRKY members, showed positive association with pre-flowering drought in RTX430.

## Availability of data and materials

The Python and R codes used for differential gene expression analysis and generating figures will be provided upon request. The supplemental data and tables have been submitted along with the manuscript.

## Acknowledgments

We thank the SRM Institute of Science and Technology, Kattankulathur, Chennai, India, for offering fellowships and the required facilities to conduct the present research. We also acknowledge SRM HPCC for providing the computational analytics infrastructure.

## Author Contributions

1. Conceptualization: SN, RNS
2. Methodology: SN, HSM
3. Investigation: SN, RNS
4. Visualization: SN, RNS, HSM
5. Supervision: RNS, HSM
6. Writing-original draft: SN
7. Writing-review & editing: SN, RNS

## Appendix S1-Figures

**Figure 1(a):**
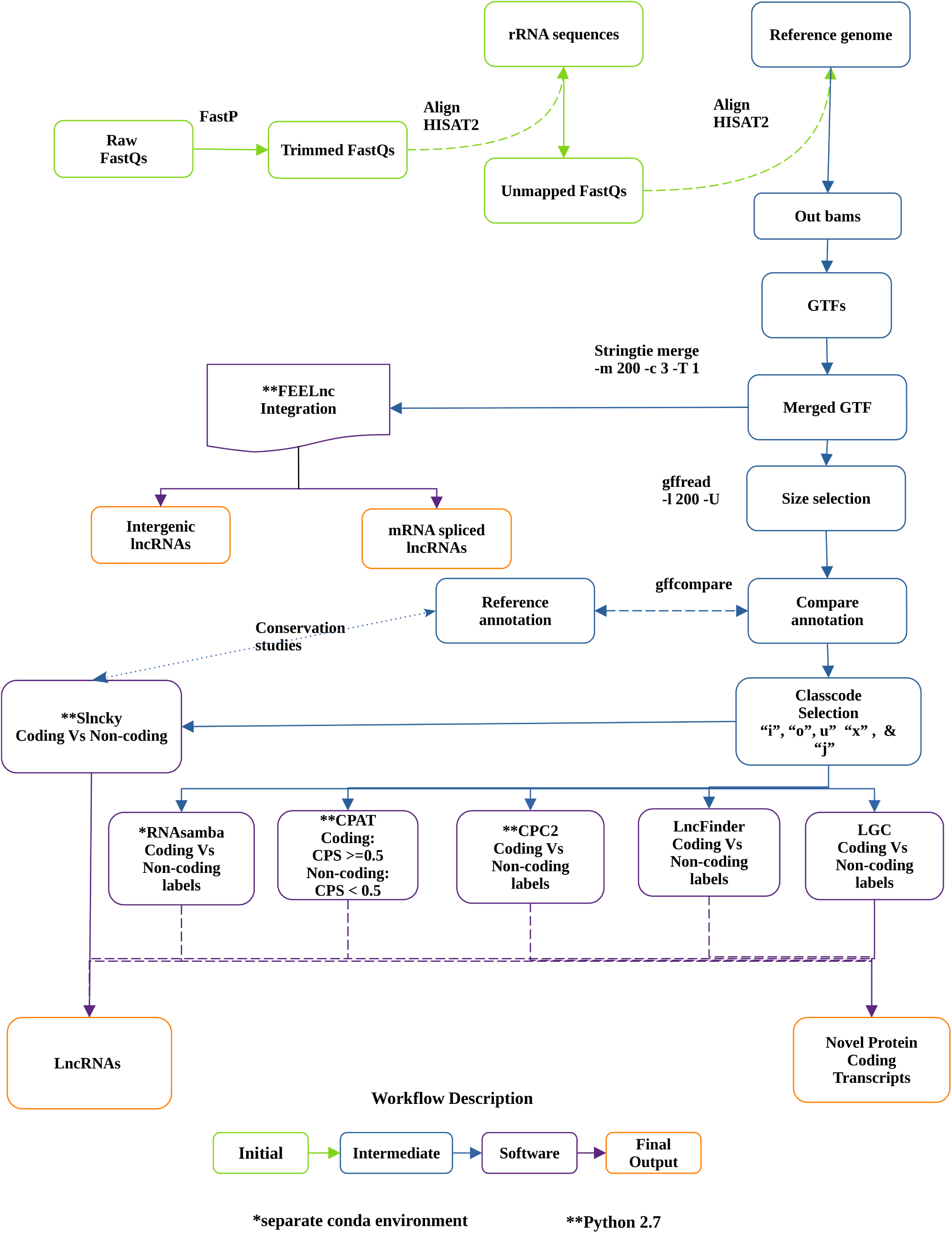
LncRAnalyzer workflow showing the integration of CPC2, CPAT, FEELnc, LGC, LncFinder,, PfamScan, RNAsamba, and Slncky tools in the automated pipeline for lncRNAs and NPCTs identification using RNA-seq datasets.

**Figure 1(b):**
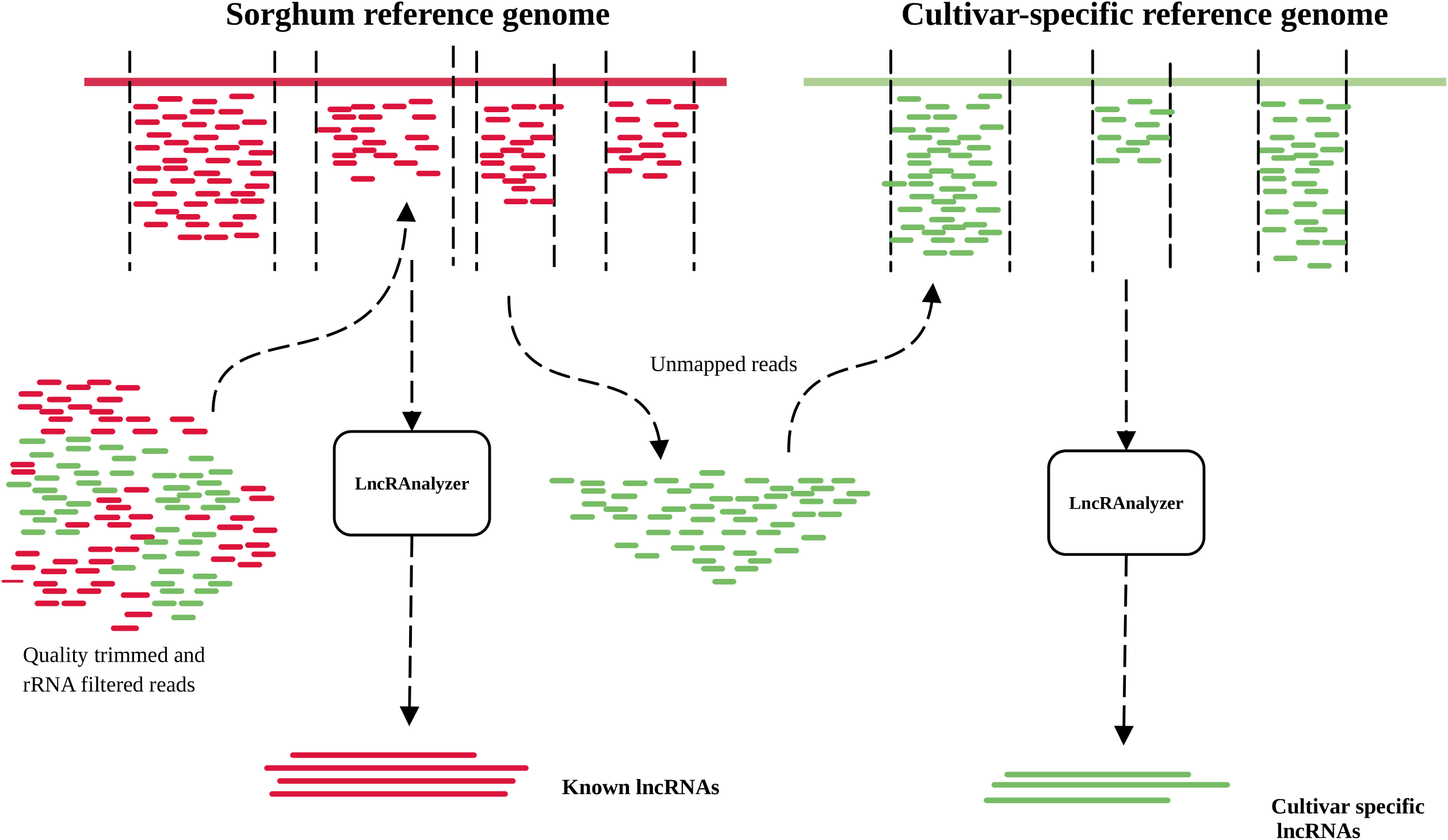
LncRAnalyzer implementation for identification of conserved and cultivar-specific lncRNAs using 389 RNA-seq accessions of BTX642 and RTX430.

**Figure 2(a):**
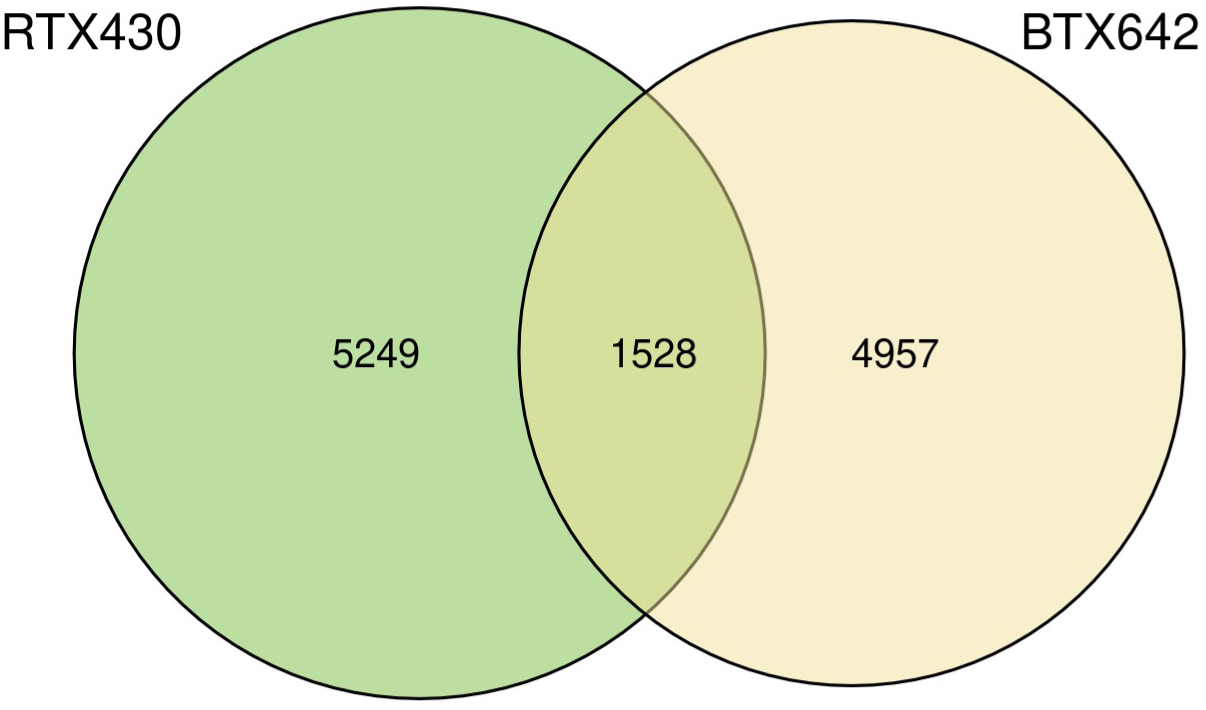
LncRNAs conservation analysis between BTX642 and RTX430 reports only 1528 lncRNAs were common between these genotypes.

**Figure 2(b):**
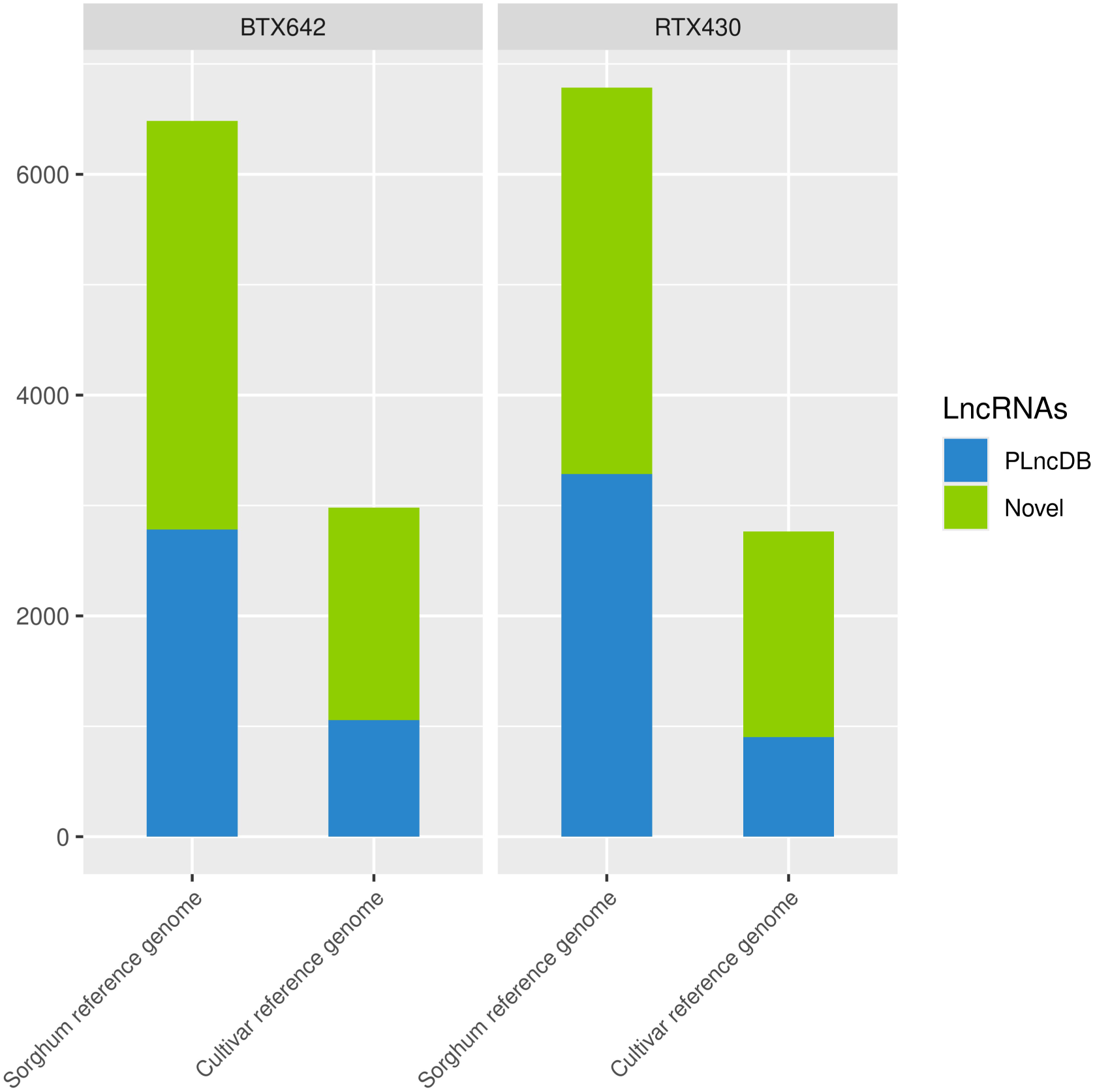
Conserved lncRNAs with PLncDB in BTX642 and RTX430 were identified using BLASTN (e-value < 10^-5^), which reports several novel lncRNAs in these genotypes.

**Figure 3:**
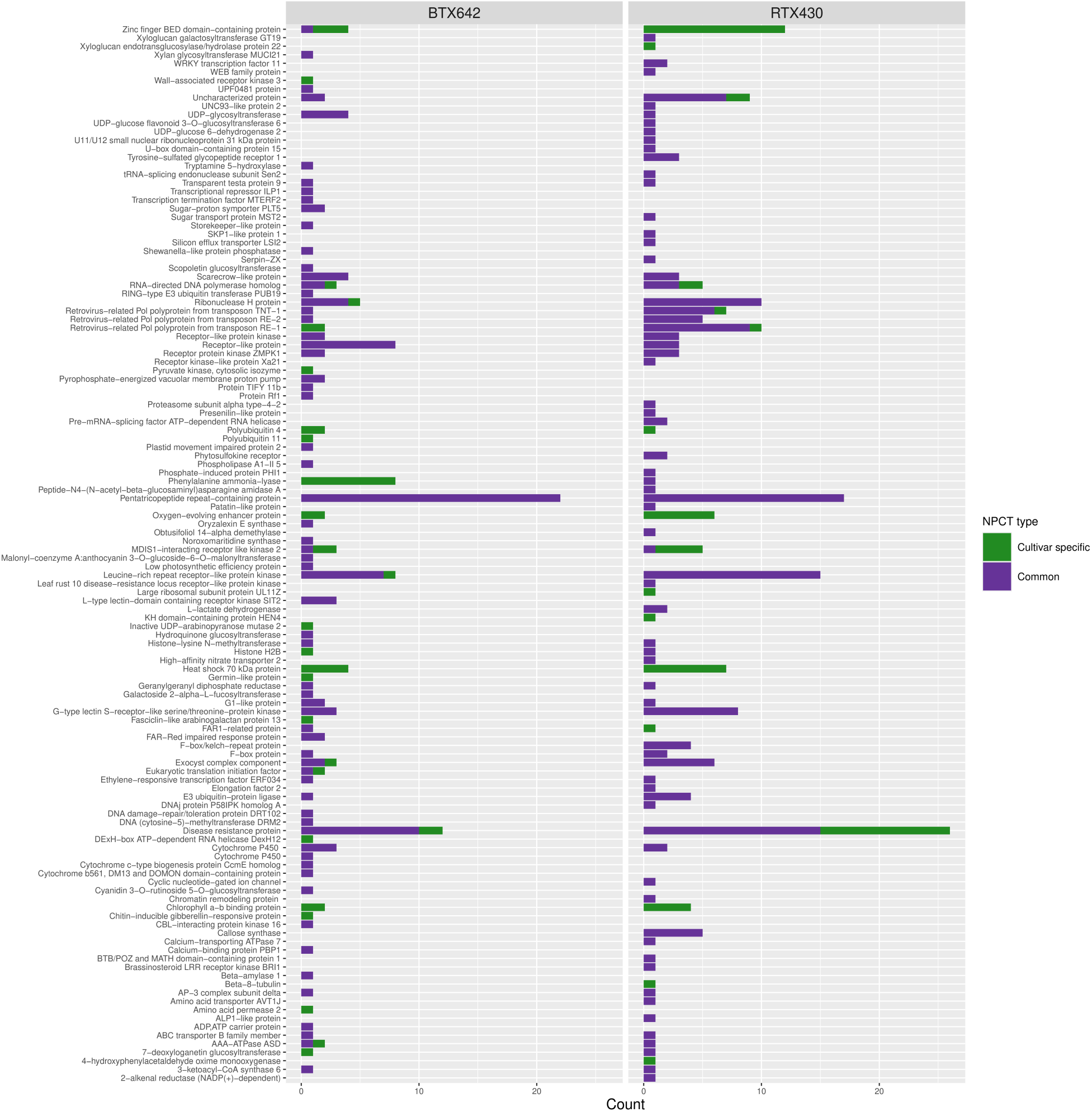
Functional annotations of NPCTs using Trinotate reports common and cultivar-specific NPCTs in BTX642 and RTX430.

**Figure 4(a):**
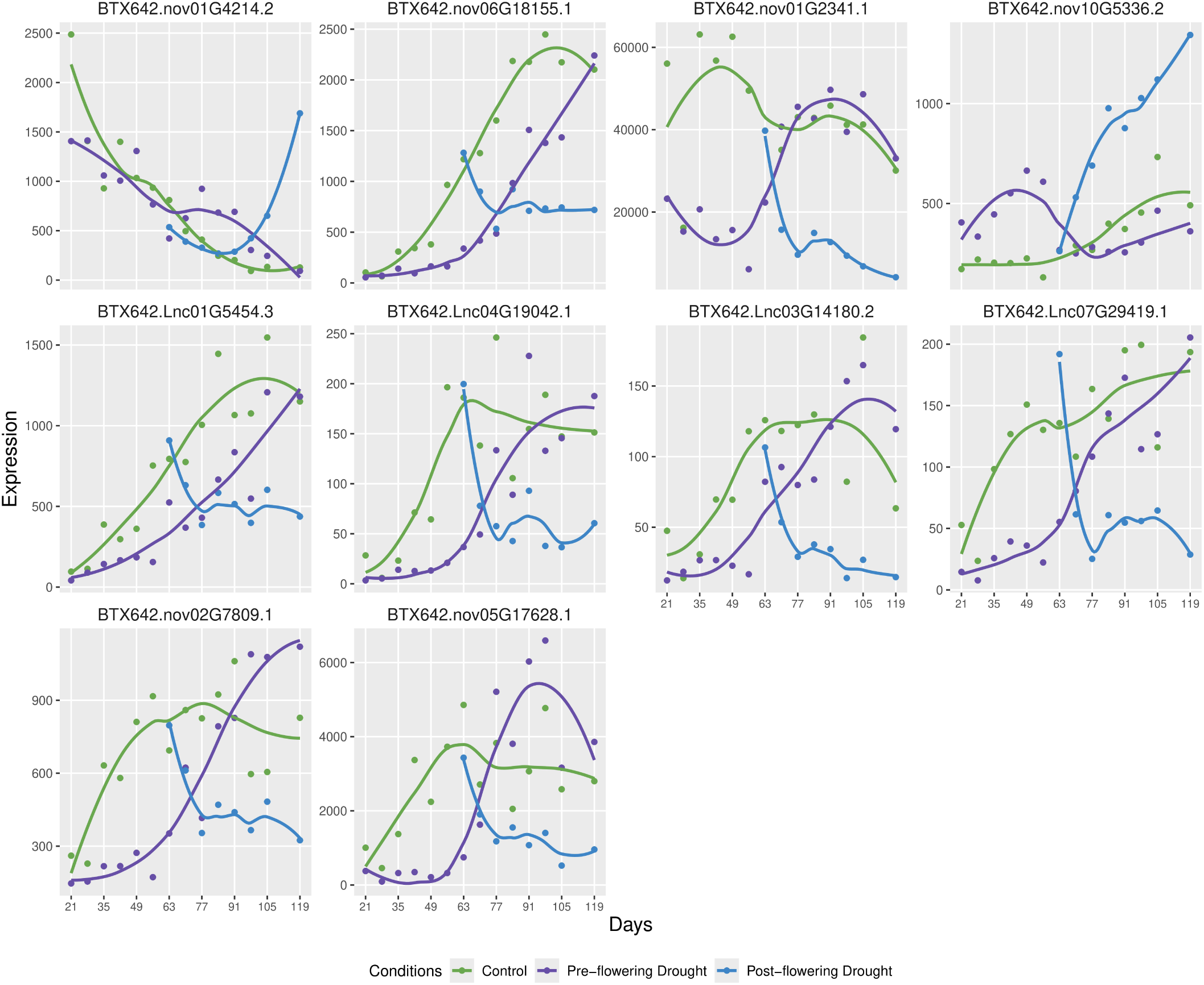
Expression patterns of the top 10 most significant lncRNAs in BTX642 leaf tissues under control, pre-, and post-flowering drought.

**Figure 4(b):**
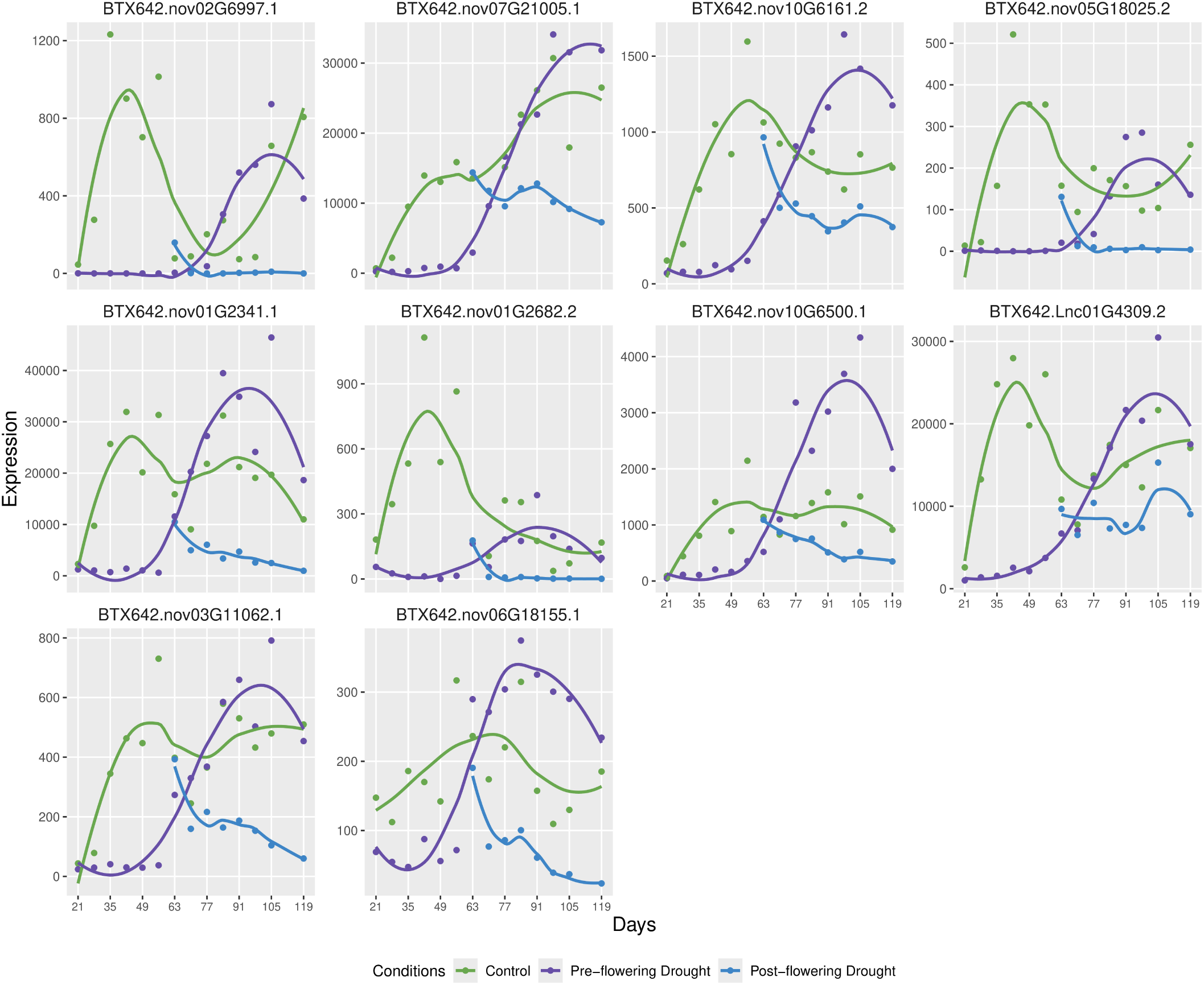
Expression patterns of the top 10 most significant lncRNAs in BTX642 root tissues under control, pre-, and post-flowering drought.

**Figure 4(c):**
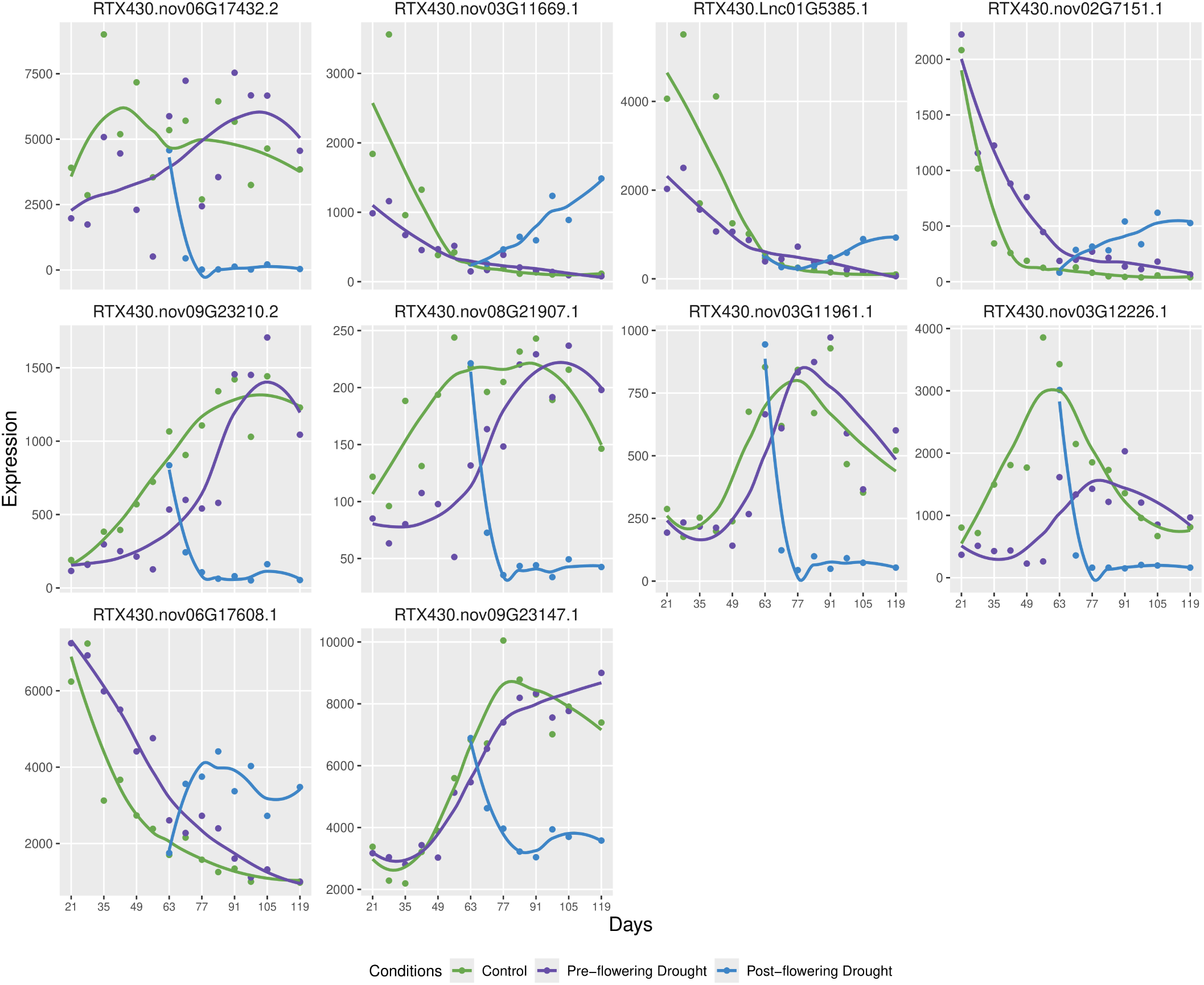
Expression patterns of the top 10 most significant lncRNAs in RTX430 leaf tissues under control, pre-, and post-flowering drought.

**Figure 4(d):**
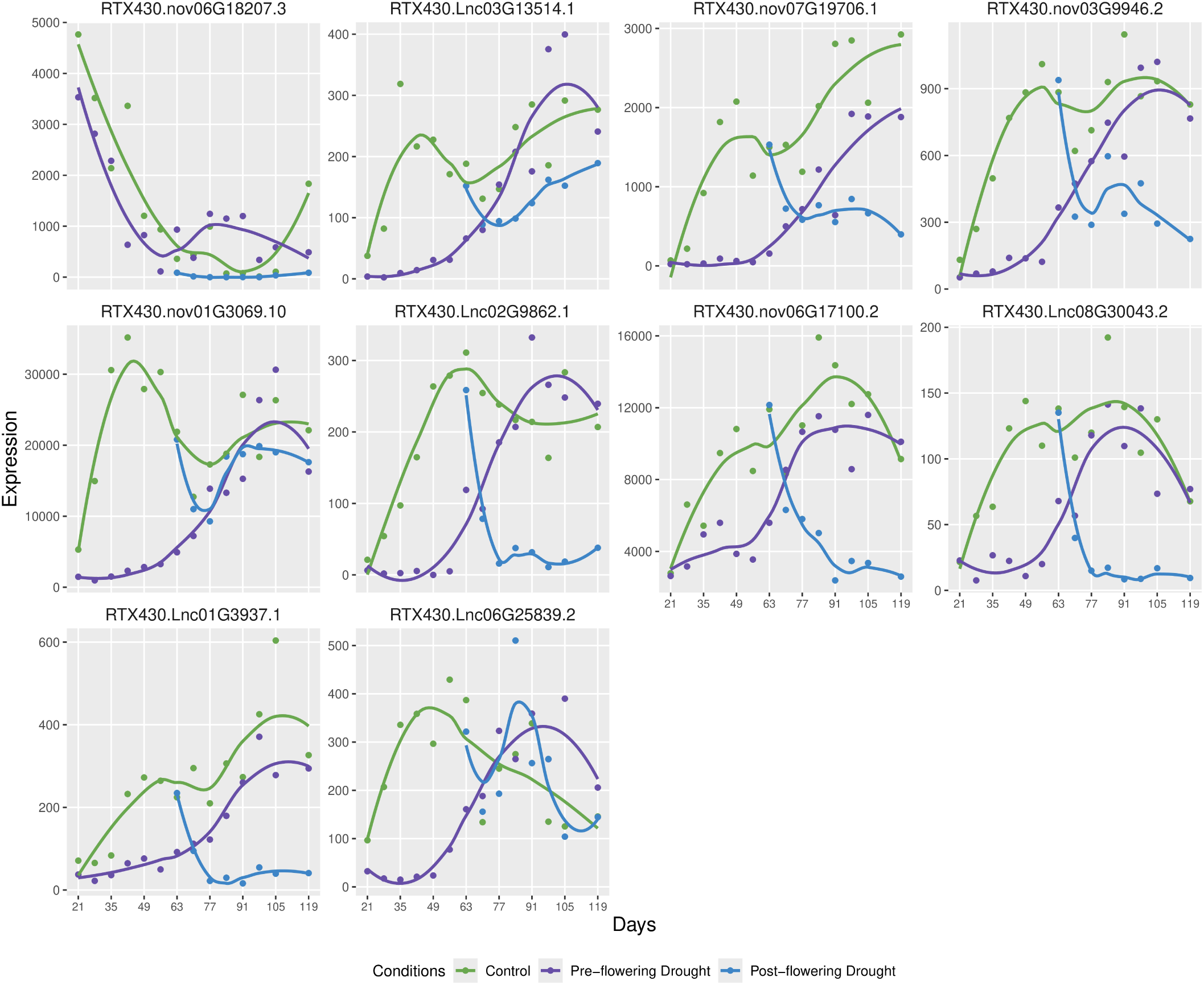
Expression patterns of the top 10 most significant lncRNAs in RTX430 root tissues under control, pre-, and post-flowering drought.

**Figure 5(a):**
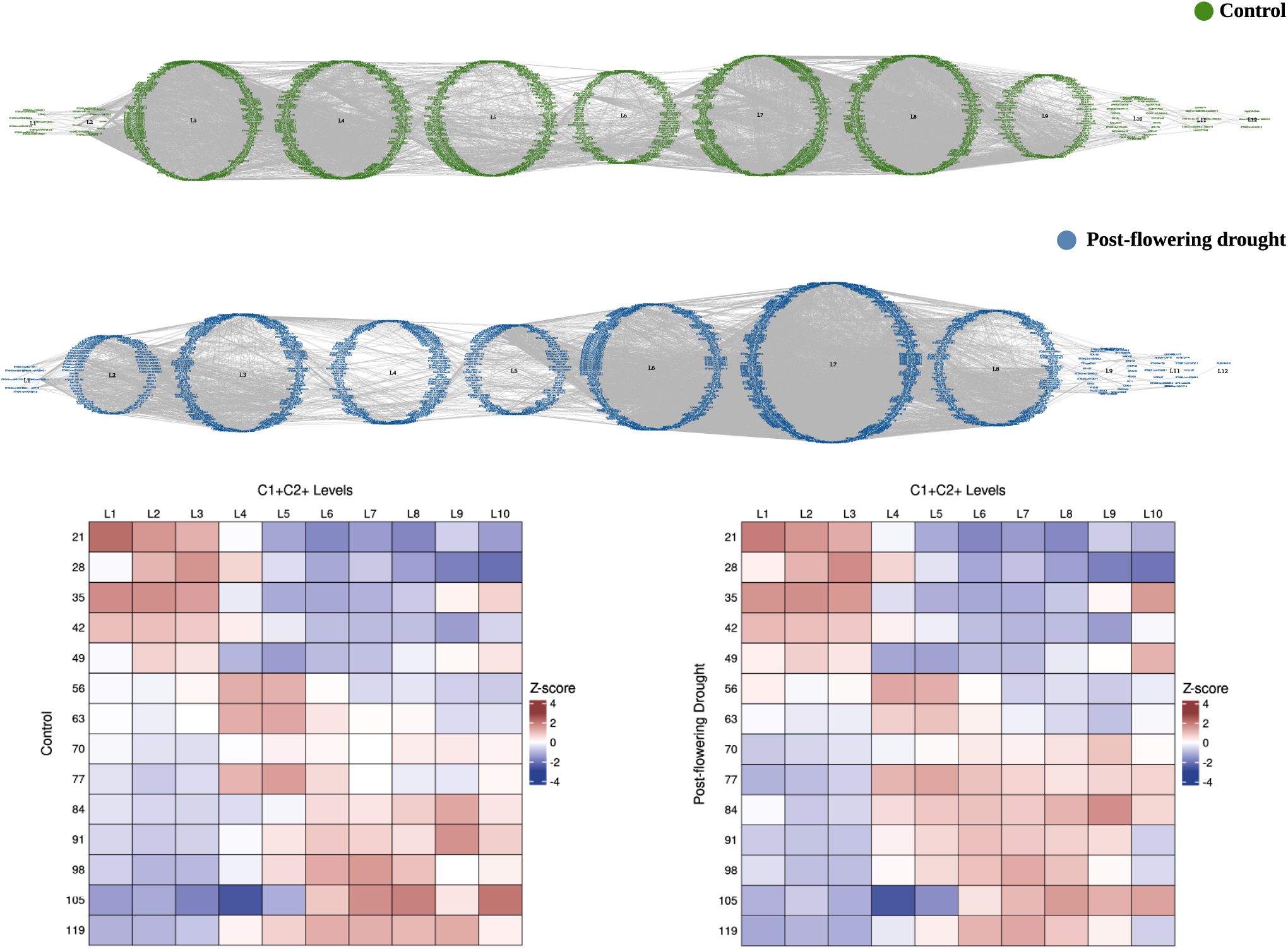
Time-ordered gene coexpression network showing co-expressed TFs and lncRNAs in different levels under control and post-flowering drought conditions in BTX642 leaf tissues.

**Figure 5(b):**
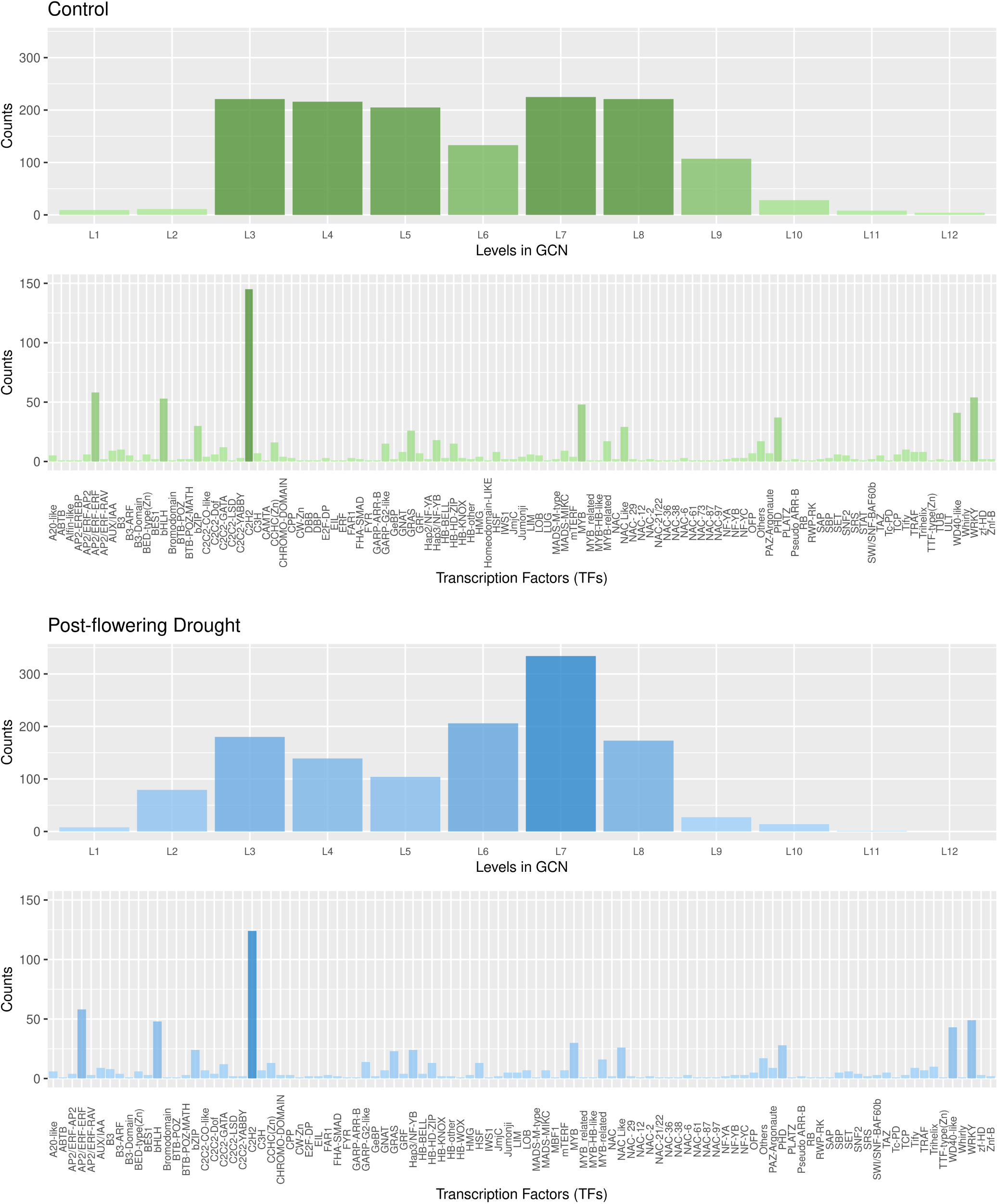
Gene coexpression analysis of BTX642 leaf under control and post-flowering drought conditions. i. Numbers of co-expressed TFs and lncRNAs in TO-GCN levels under control conditions. ii. Numbers of various TFs co-expressed under control conditions. iii. Numbers of co-expressed TFs and lncRNAs in TO-GCN levels under post-flowering drought conditions. iv. Numbers of various TFs co-expressed under post-flowering drought conditions.

**Figure 5(c):**
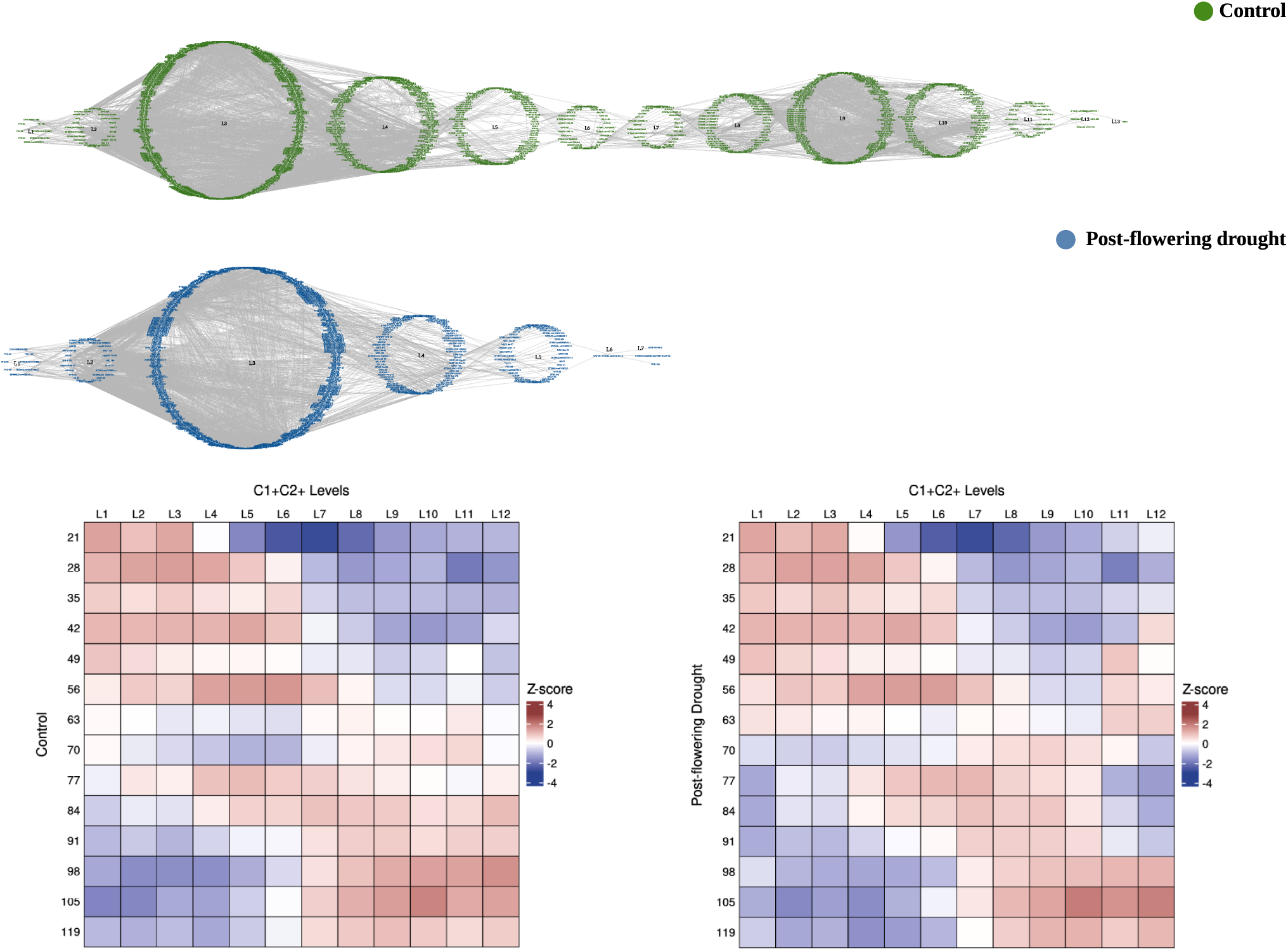
Time-ordered gene coexpression network showing co-expressed TFs and lncRNAs in different levels under control and post-flowering drought conditions in BTX642 root tissues.

**Figure 5(d):**
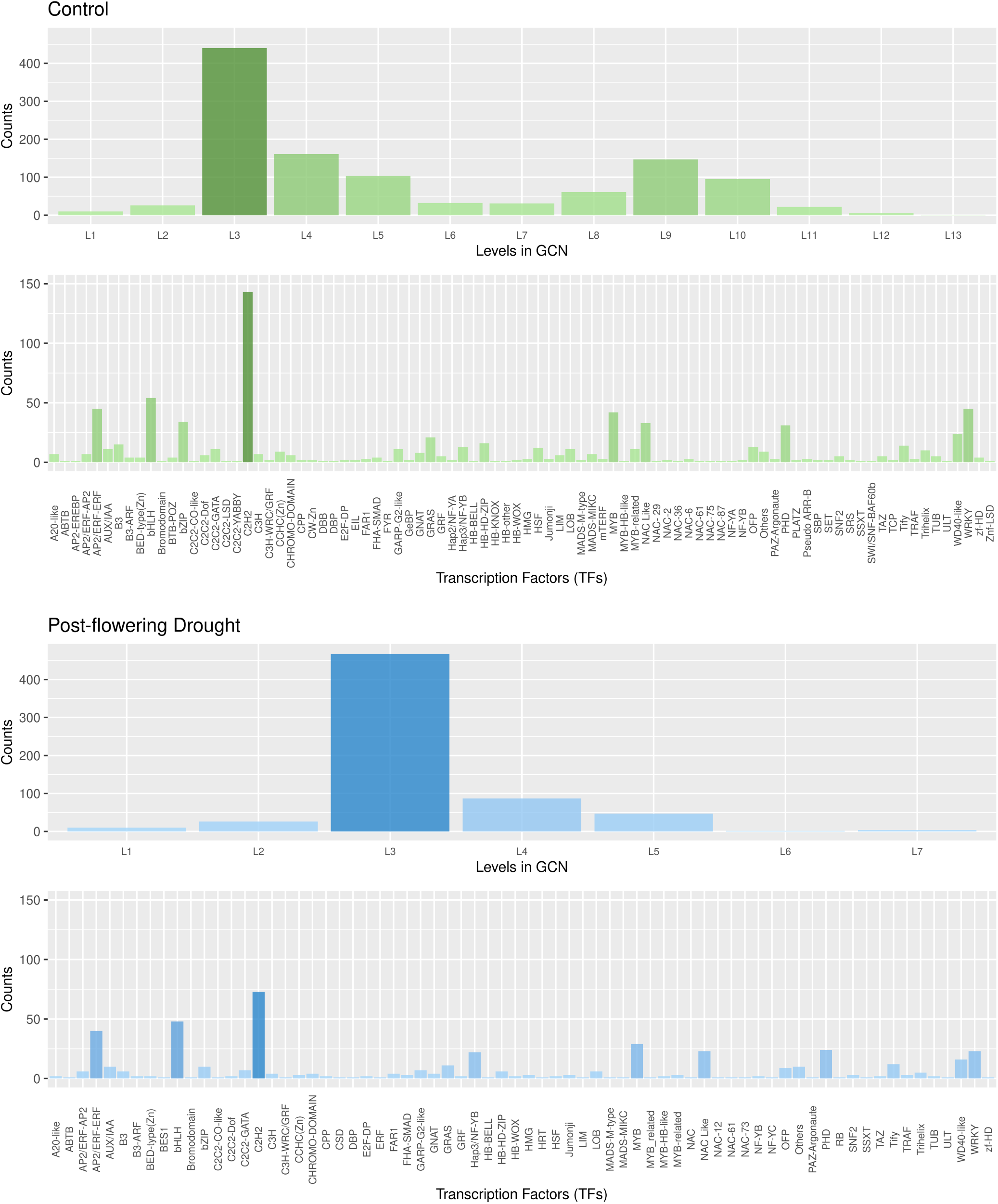
Gene coexpression analysis of BTX642 roots under control and post-flowering drought conditions. i. Numbers of co-expressed TFs and lncRNAs in TO-GCN levels under control conditions. ii. Numbers of various TFs co-expressed under control conditions. iii. Numbers of co-expressed TFs and lncRNAs in TO-GCN levels under post-flowering drought conditions. iv. Numbers of various TFs co-expressed under post-flowering drought conditions.

**Figure 5(e):**
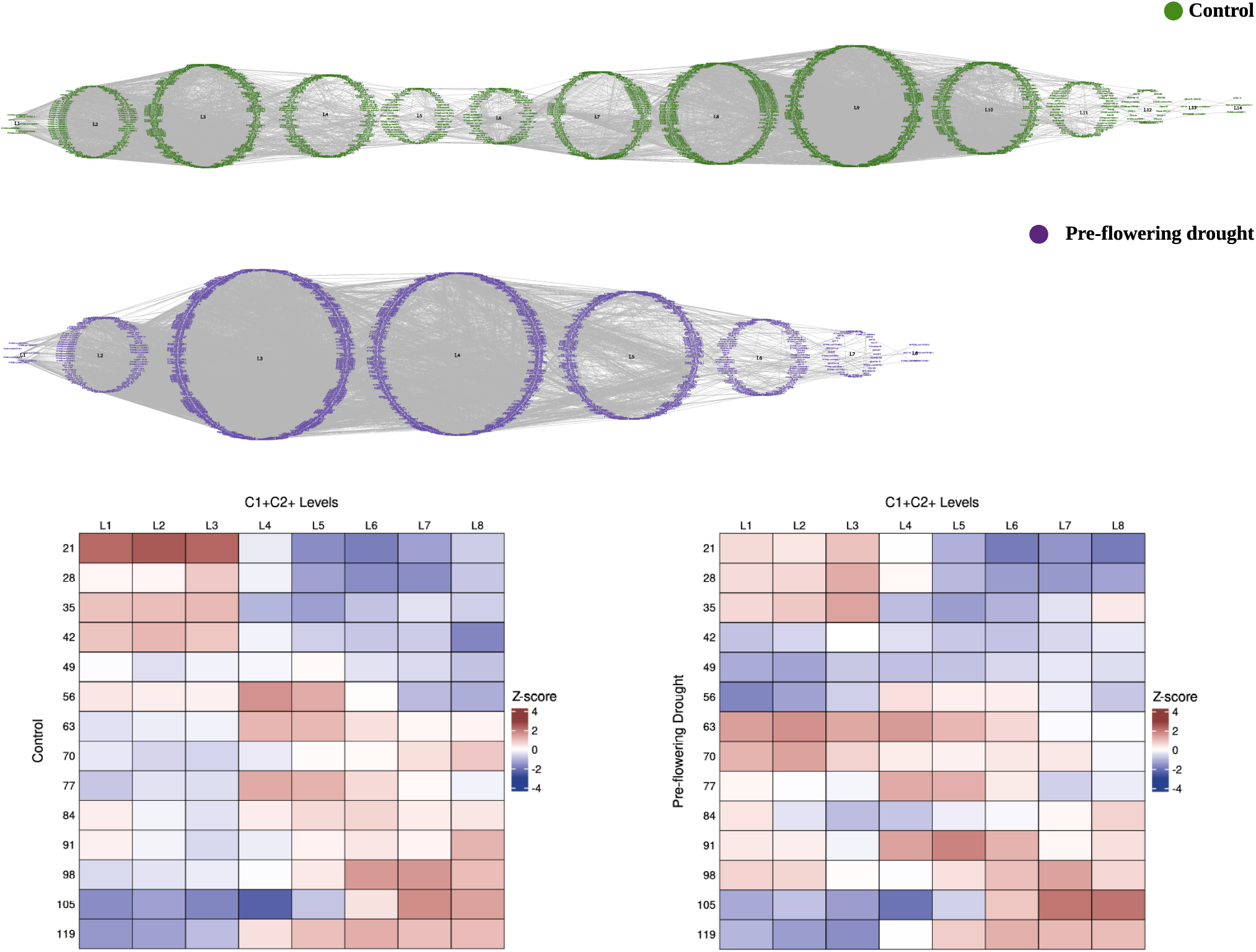
Time-ordered gene coexpression network showing co-expressed TFs and lncRNAs in different levels under control and pre-flowering drought conditions in RTX430 leaf tissues.

**Figure 5(f):**
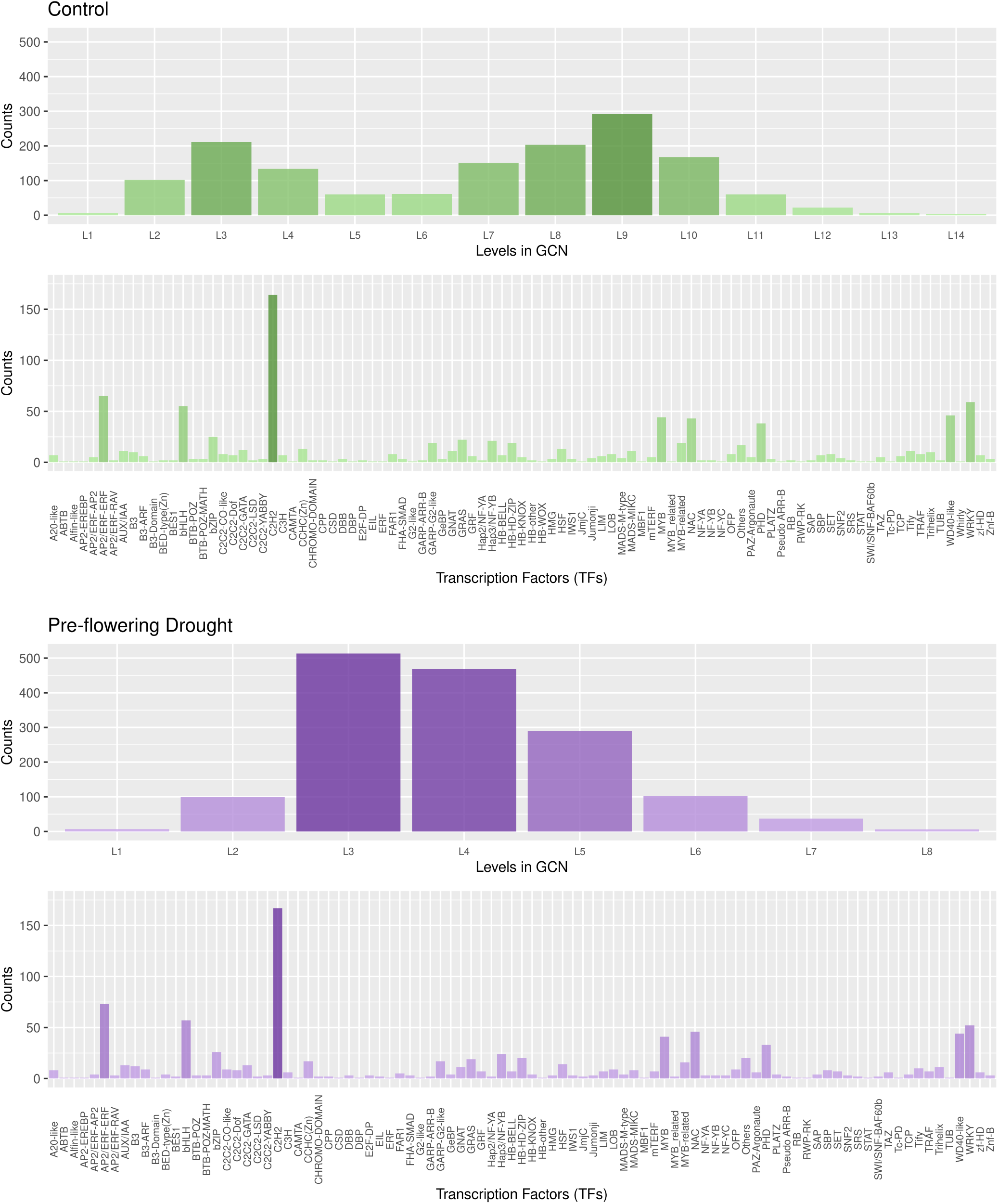
Gene coexpression analysis of RTX430 leaf under control and pre-flowering drought conditions. i. Numbers of co-expressed TFs and lncRNAs in TO-GCN levels under control conditions. ii. Numbers of various TFs co-expressed under control conditions. iii. Numbers of co-expressed TFs and lncRNAs in TO-GCN levels under pre-flowering drought conditions. iv. Numbers of various TFs co-expressed under pre-flowering drought conditions.

**Figure 5(g):**
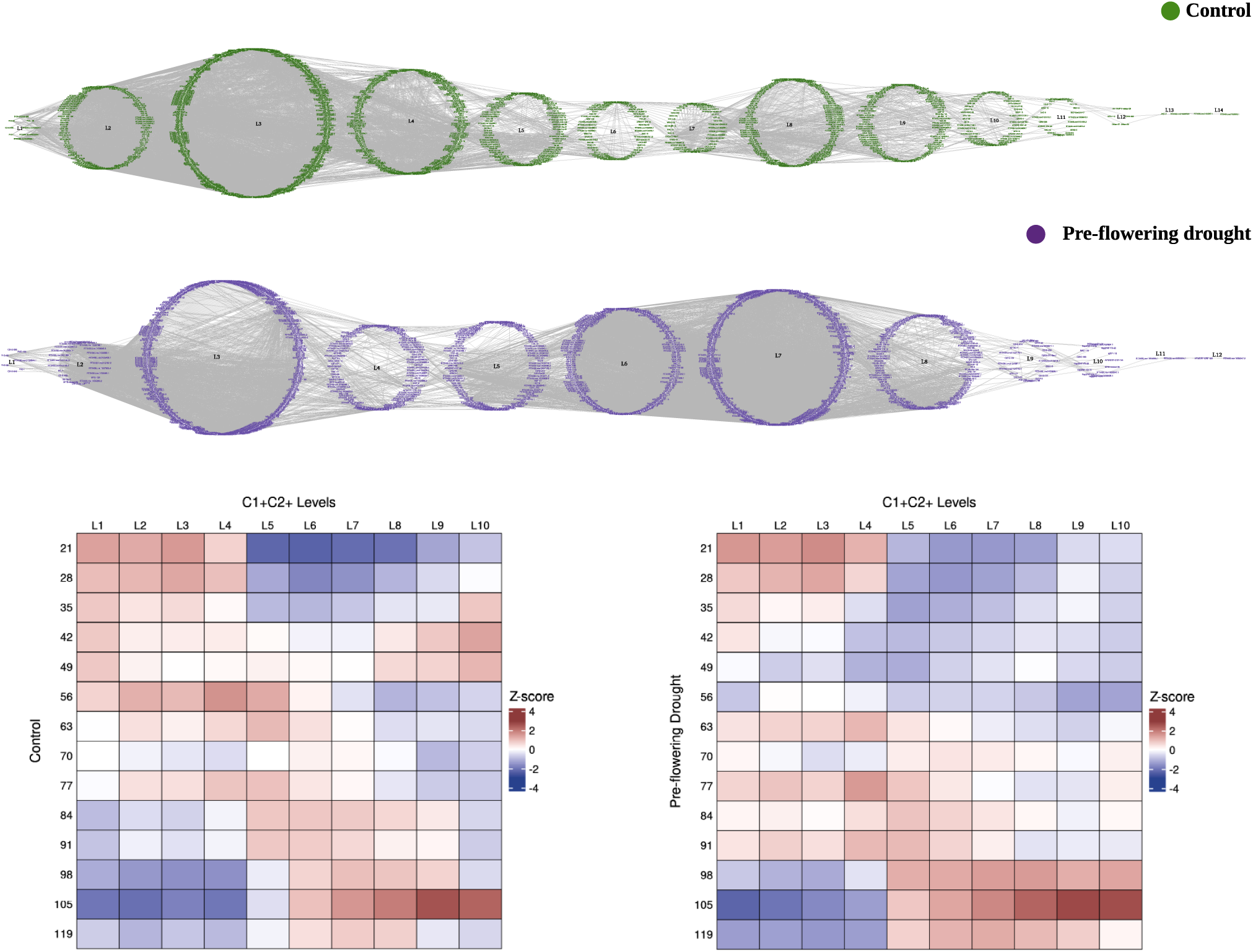
Time-ordered gene coexpression network showing co-expressed TFs and lncRNAs in different levels under control and pre-flowering drought conditions in RTX430 root tissues.

**Figure 5(h):**
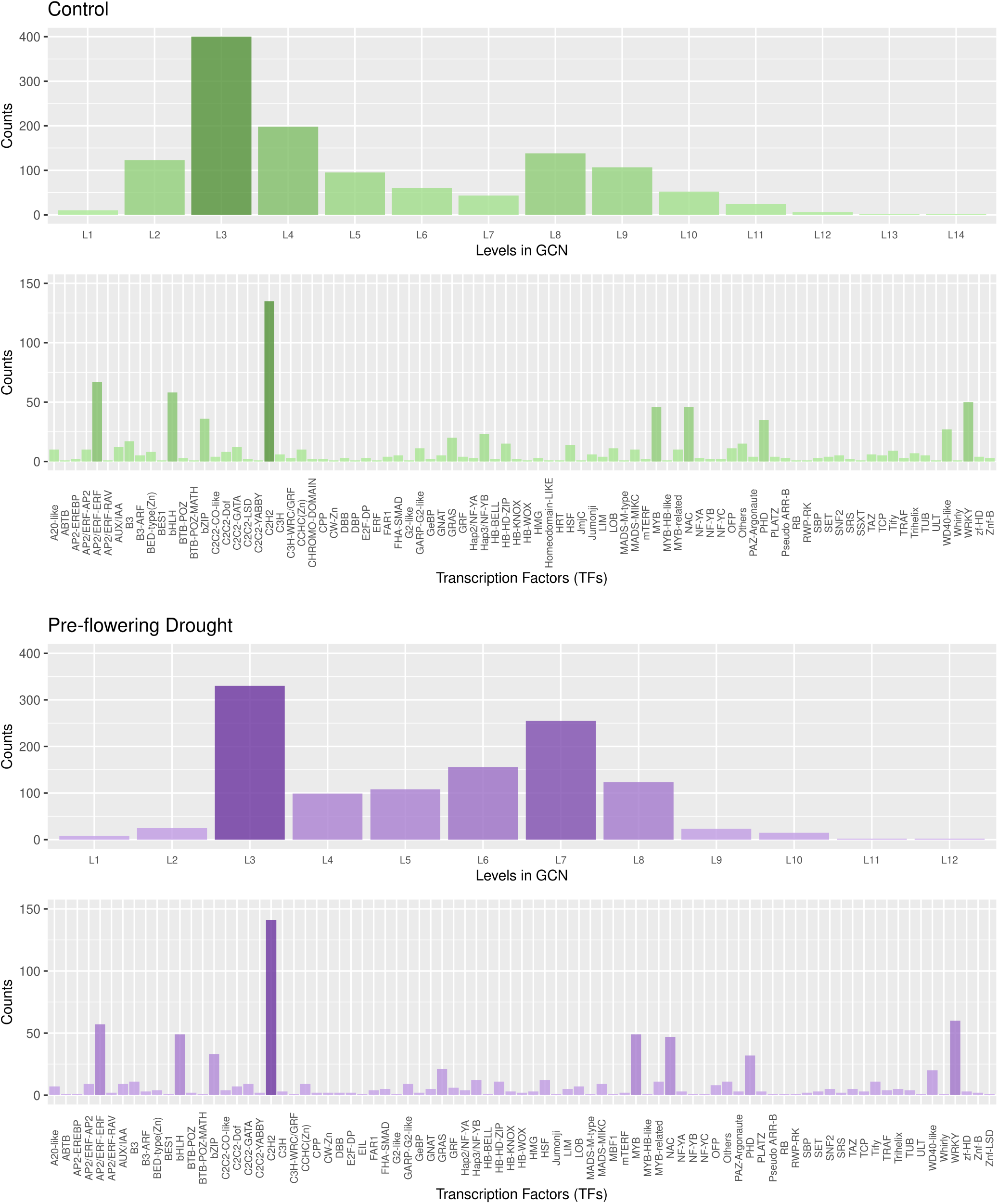
Gene coexpression analysis of RTX430 roots under control and pre-flowering drought conditions. i. Numbers of co-expressed TFs and lncRNAs in TO-GCN levels under control conditions. ii. Numbers of various TFs co-expressed under control conditions. iii. Numbers of co-expressed TFs and lncRNAs in TO-GCN levels under pre-flowering drought conditions. iv. Numbers of various TFs co-expressed under pre-flowering drought conditions.

**Figure 6(a):**
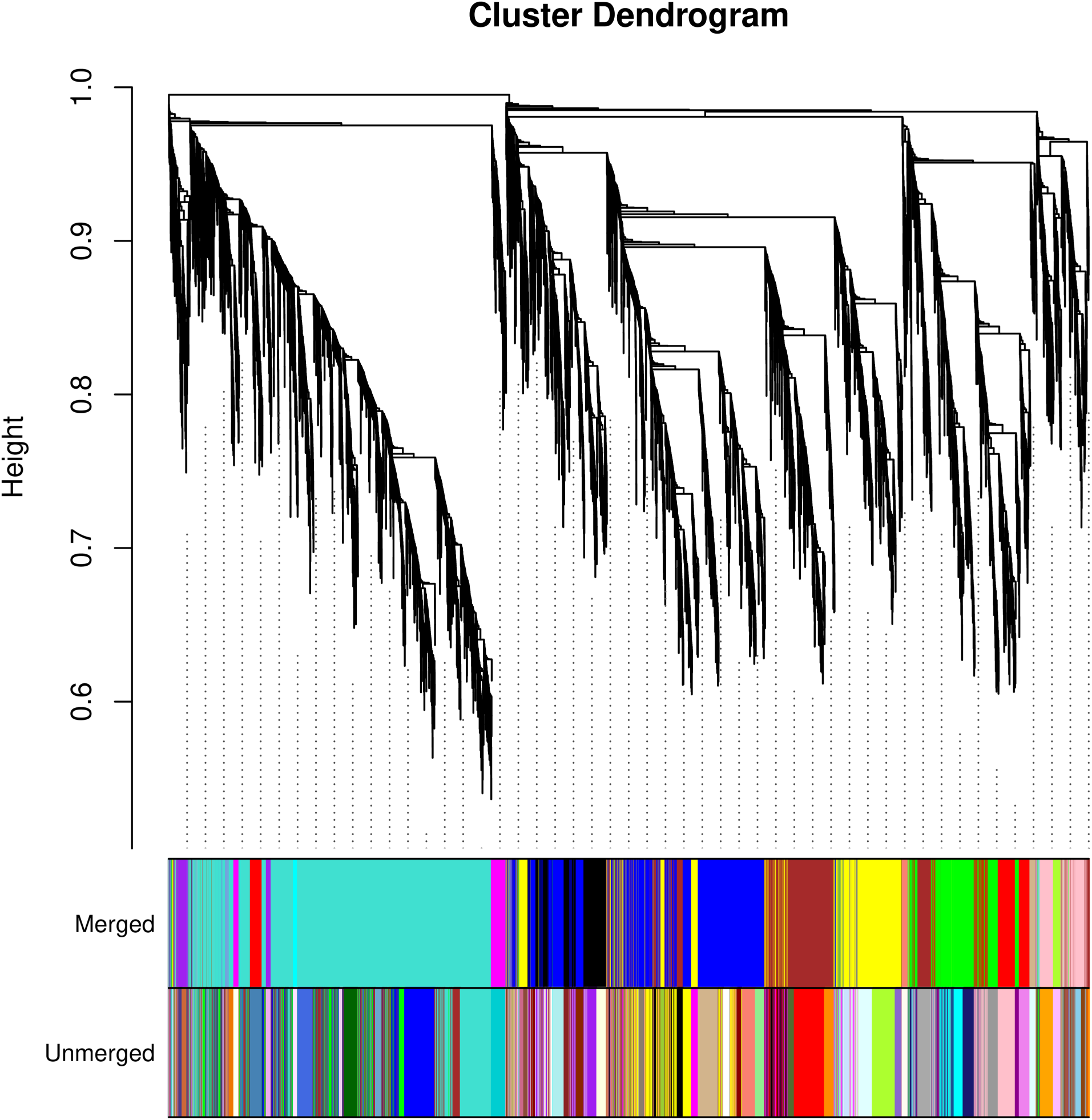
WGCNA cluster dendrogram showing various merged and unmerged modules in BTX642 leaf tissues.

**Figure 6(b):**
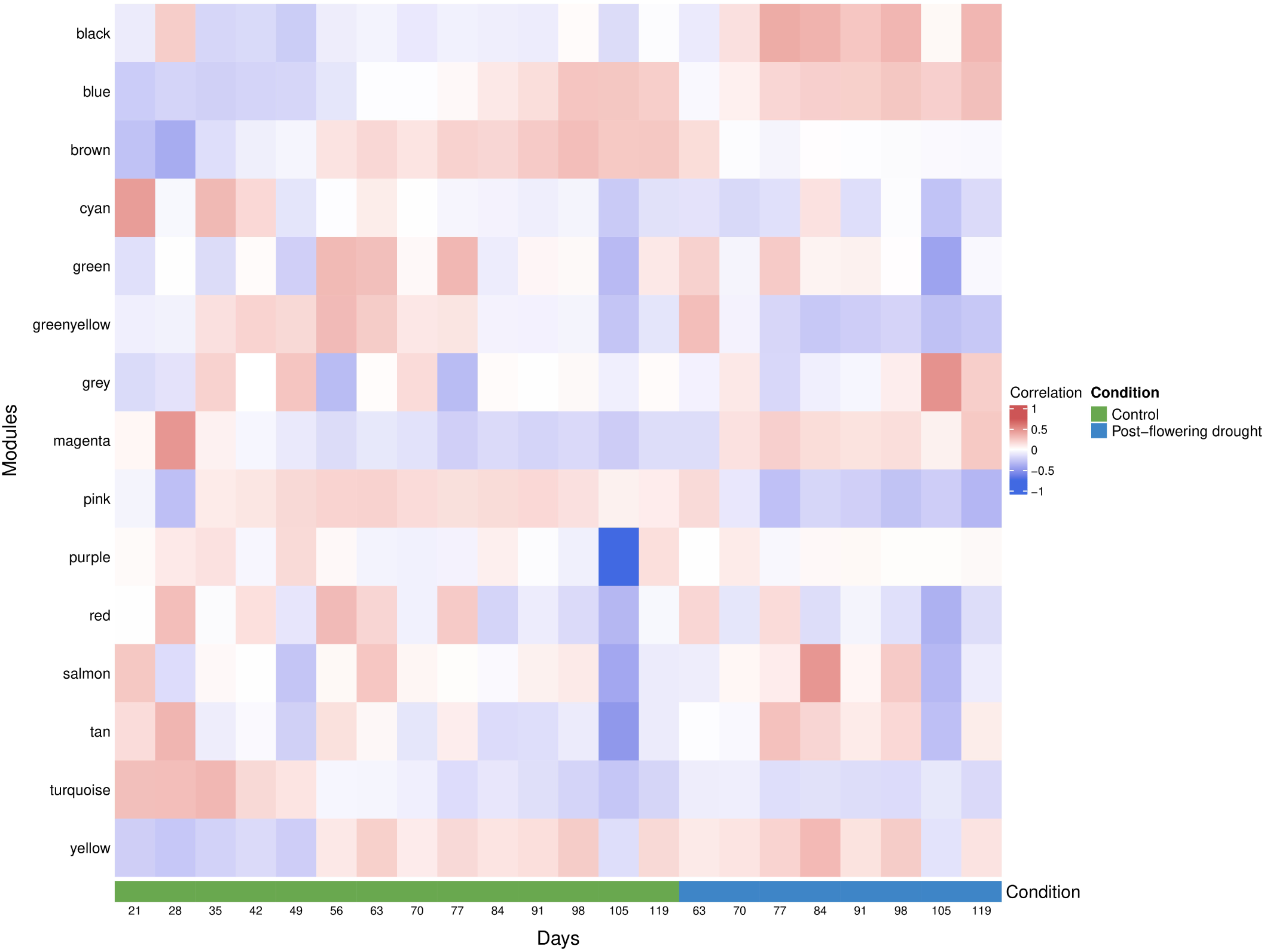
WGCNA module-trait relationship showing “Black”, “Magenta”, and “Tan” coexpression modules in BTX642 leaf tissues, which were associated with post-flowering drought tolerance.

**Figure 6(c):**
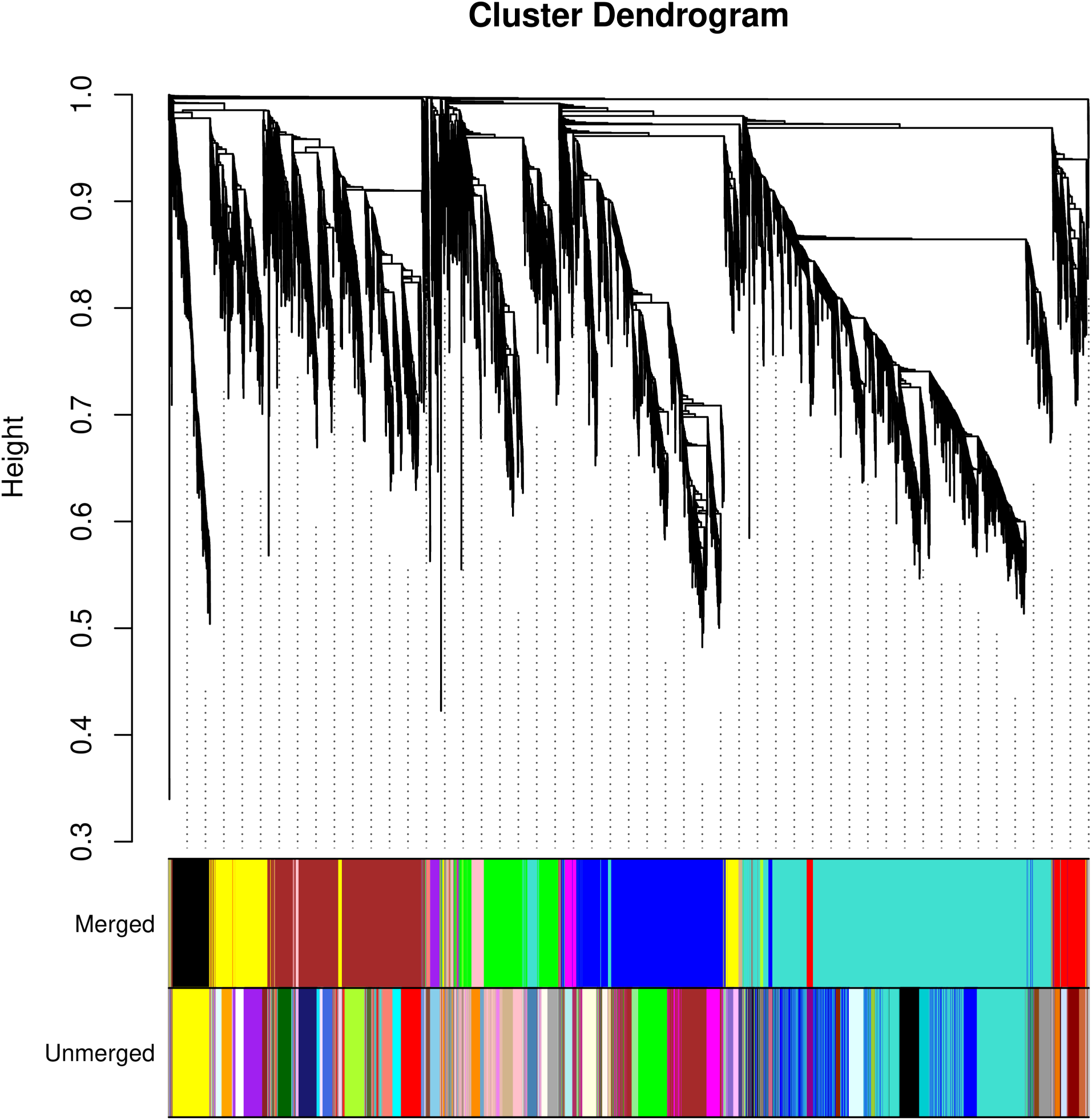
WGCNA cluster dendrogram showing various merged and unmerged modules in BTX642 root tissues.

**Figure 6(d):**
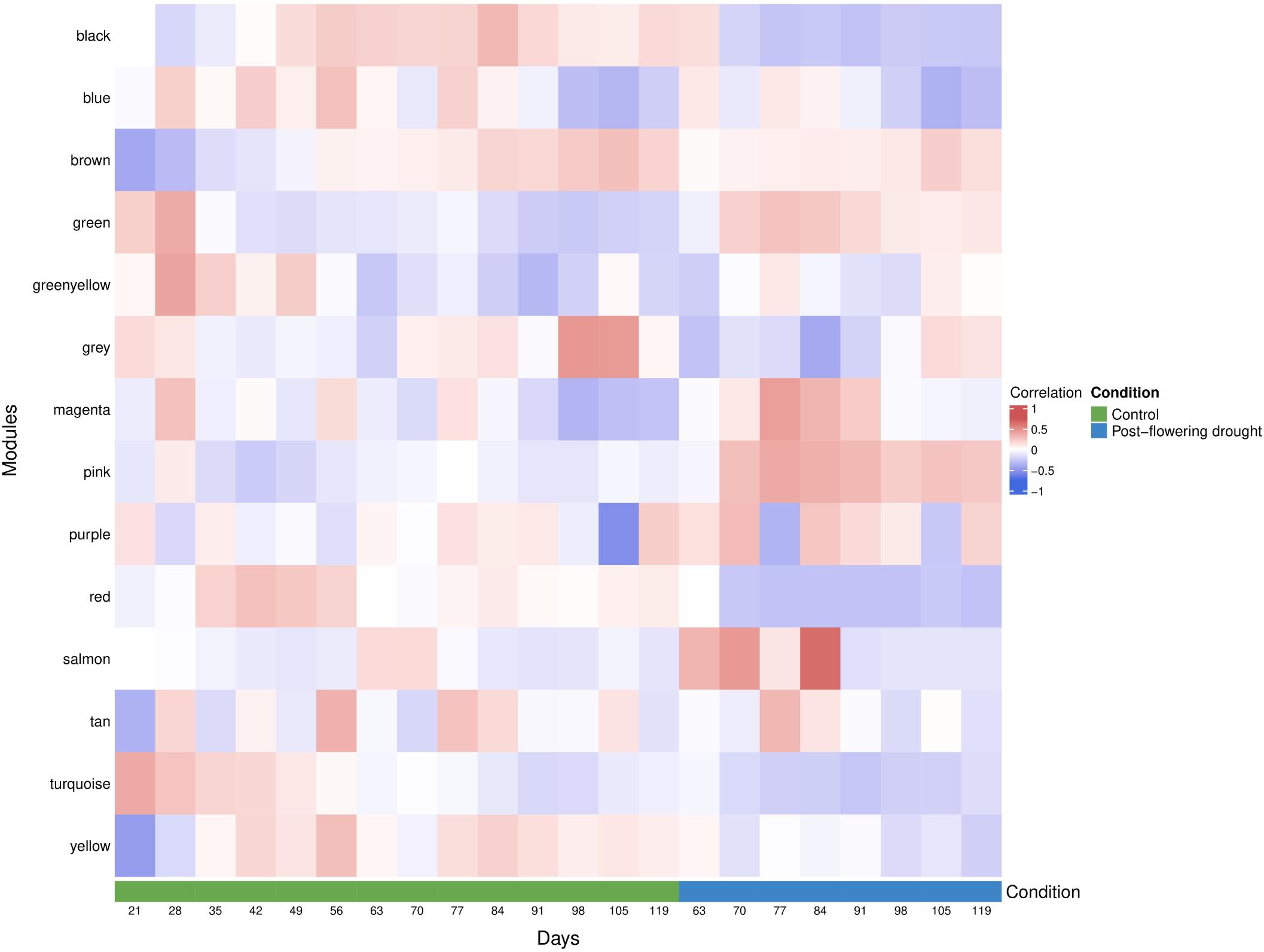
WGCNA module-trait relationship showing “Green”, “Pink”, “Magenta”, and “Salmon” coexpression modules in BTX642 root tissues, which were associated with post-flowering drought tolerance.

**Figure 6(e):**
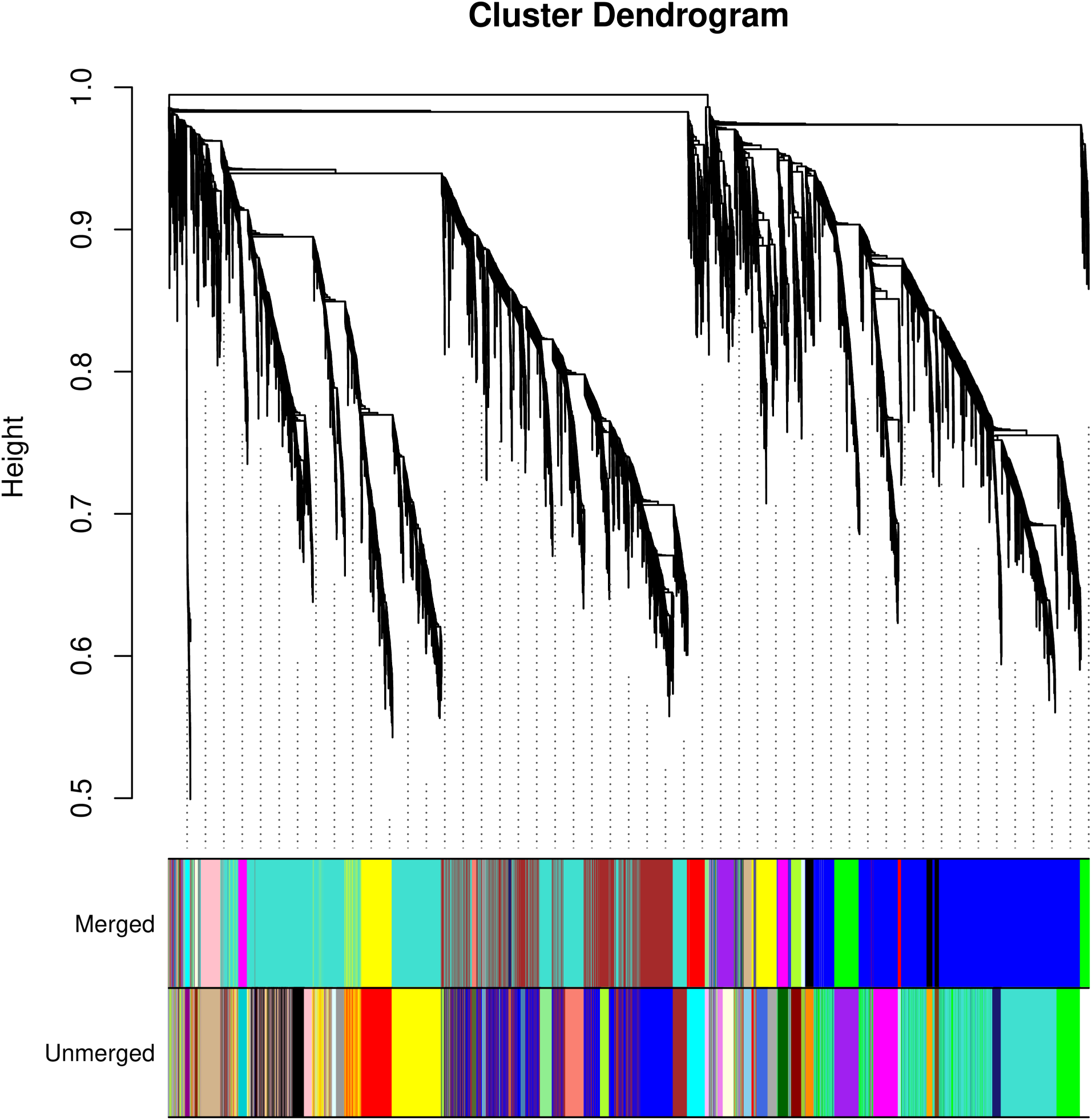
WGCNA cluster dendrogram showing various merged and unmerged modules in RTX430 leaf tissues.

**Figure 6(f):**
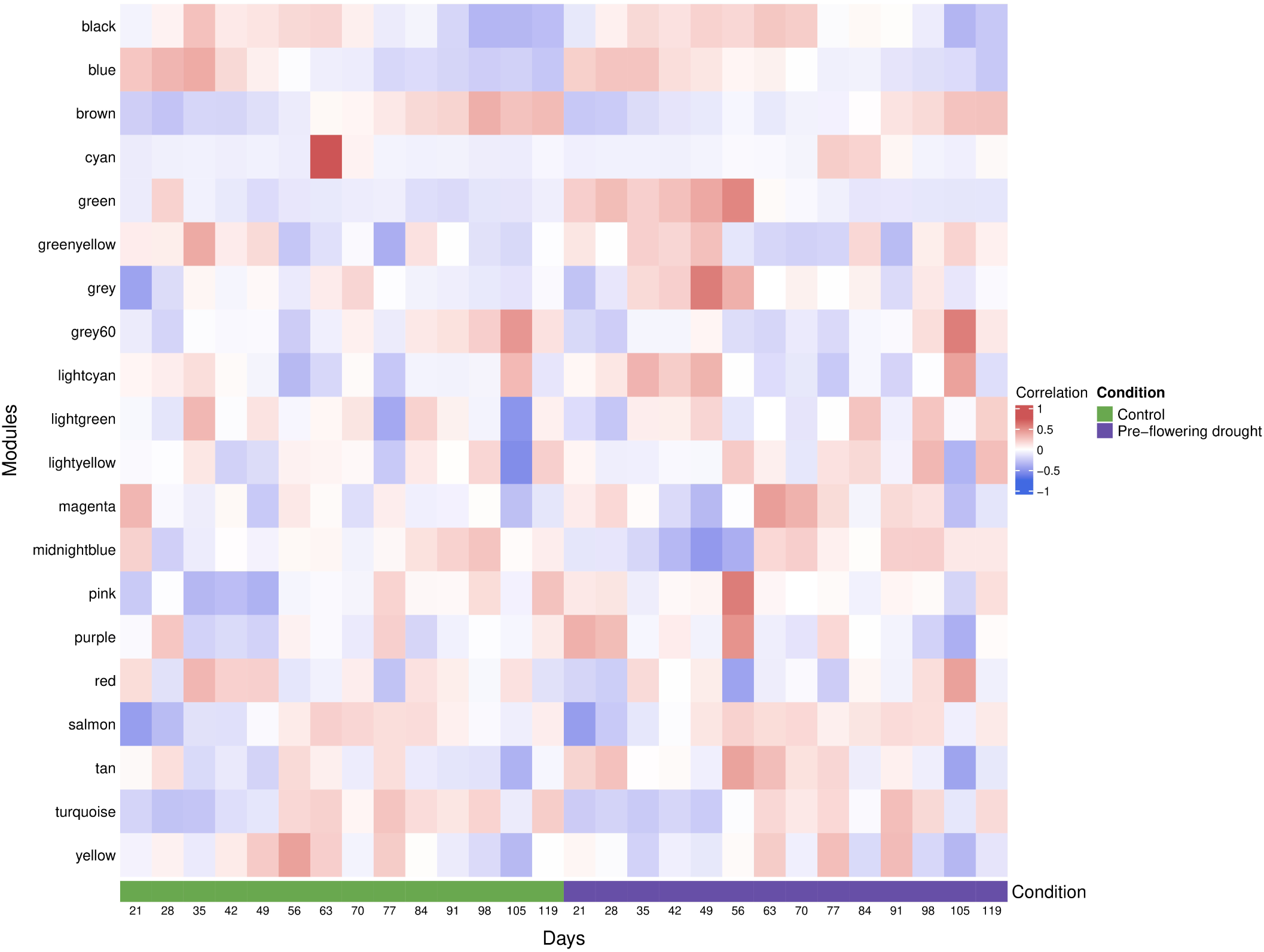
WGCNA module-trait relationship showing “Green”, “Grey”, “Magenta”, and “Lightcyan” coexpression modules in RTX430 leaf tissues, which were associated with pre-flowering drought tolerance.

**Figure 6(g):**
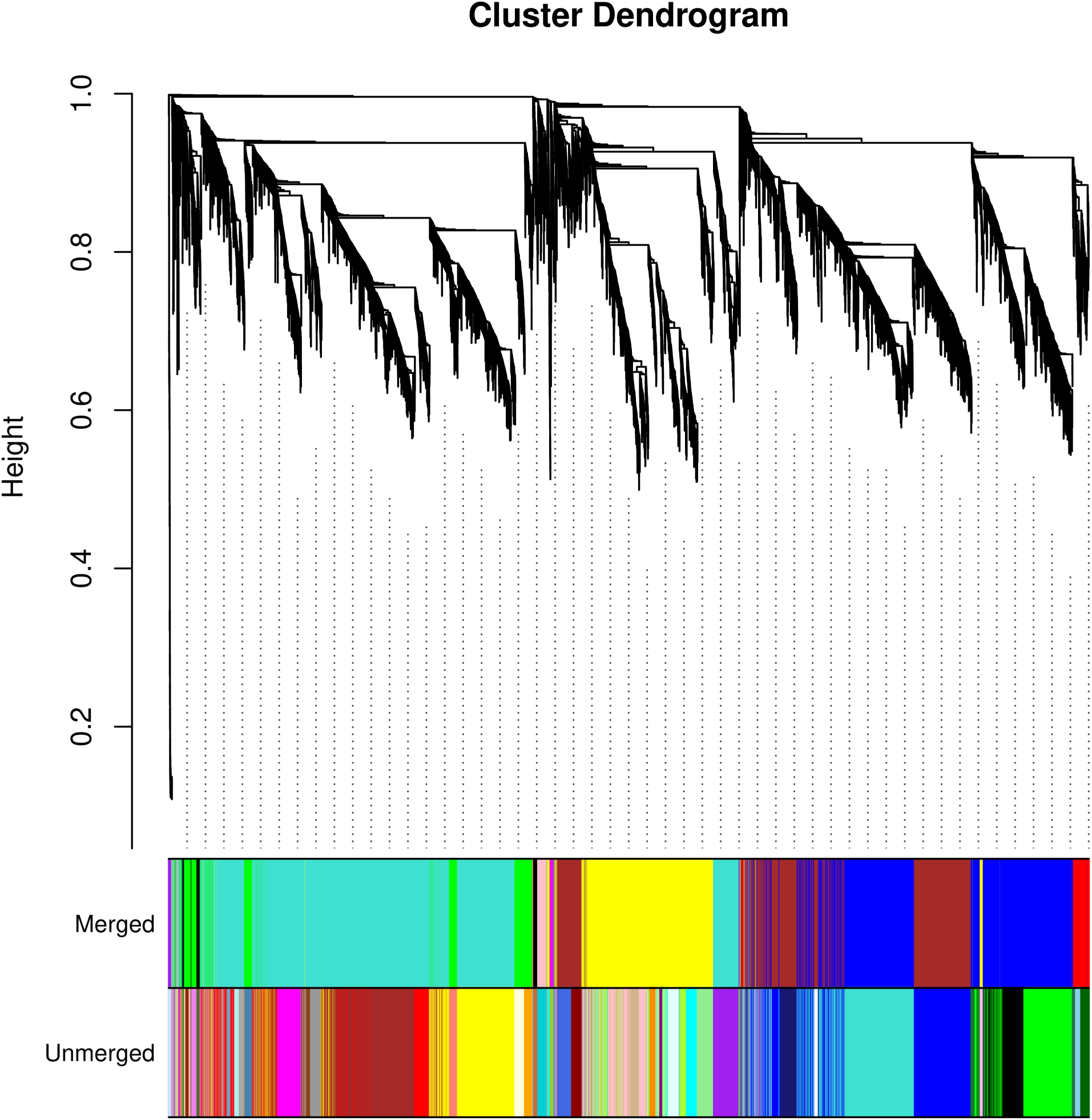
WGCNA cluster dendrogram showing various merged and unmerged modules in RTX430 root tissues.

**Figure 6(h):**
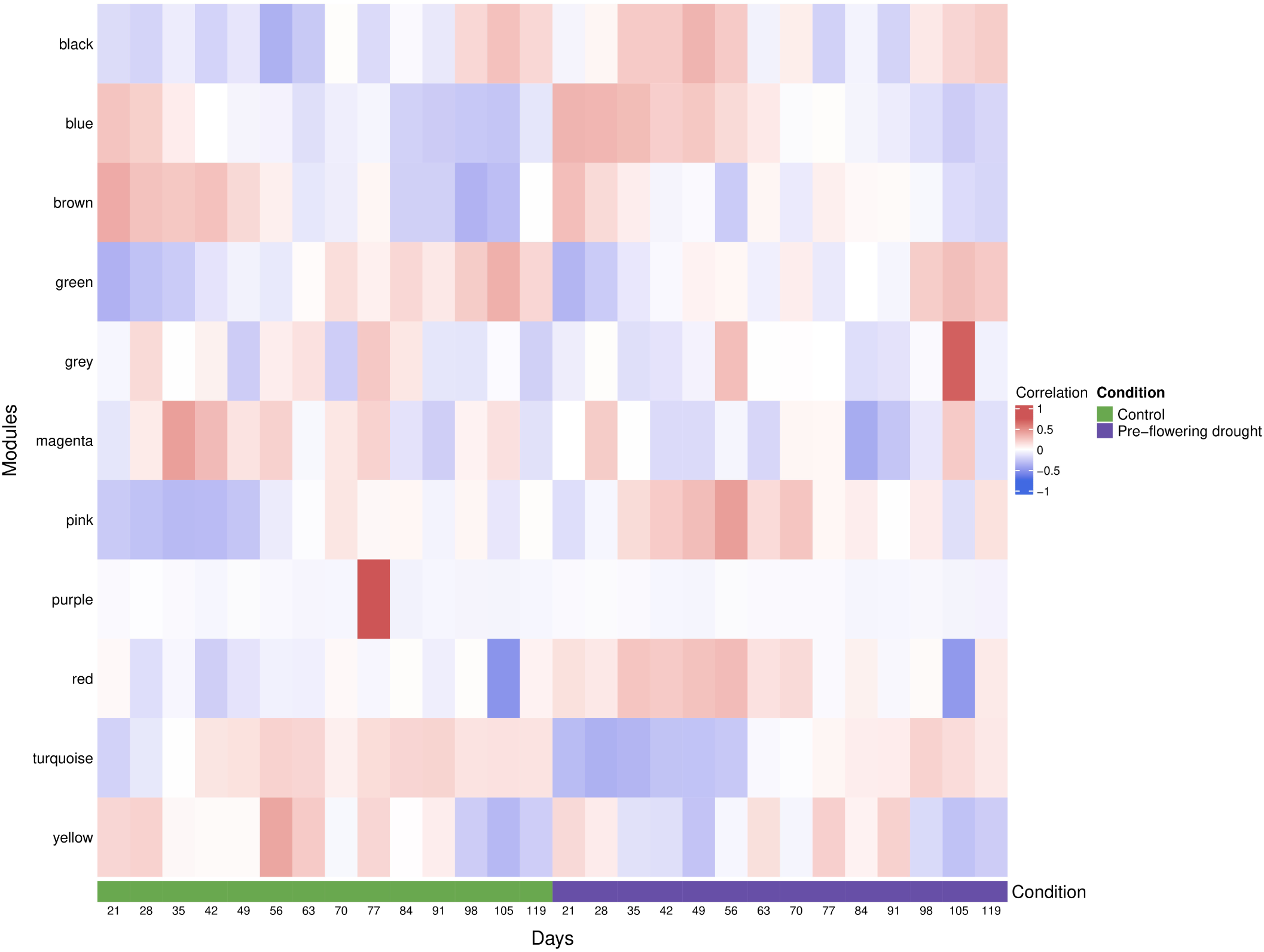
WGCNA module-trait relationship showing “Black”, “Blue”, “Pink”, and “Red” coexpression modules in RTX430 root tissues, which were associated with pre-flowering drought tolerance.

