## Appendix S1- Supplemental Figures for "Uncovering Long Non-Coding RNAs and Exploring Gene Coexpression Patterns in Sorghum Genomics"

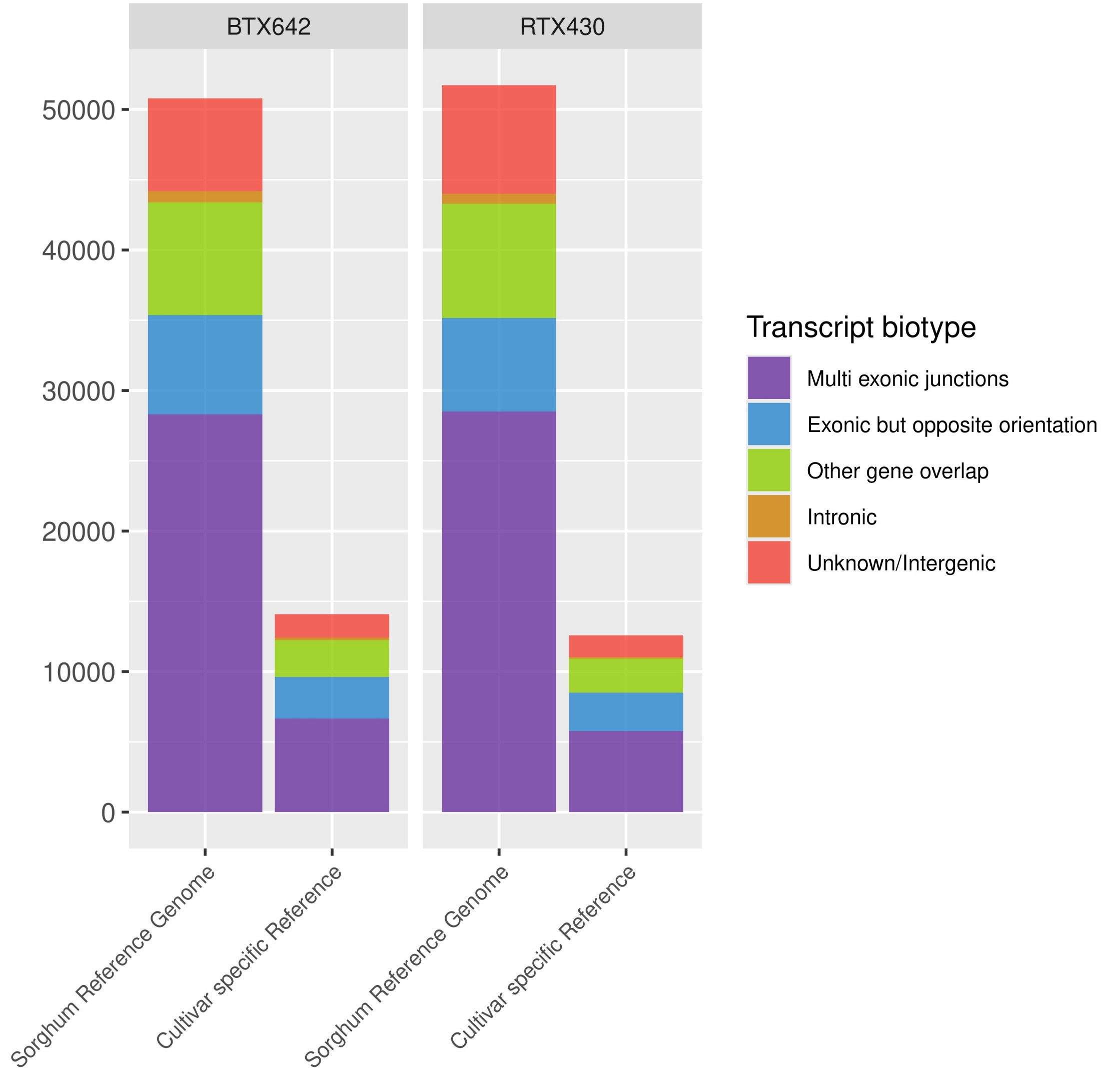


**Supplemental Figure S1(a)**: The number of transcripts belonging to class codes ‘i’, ‘o’, ‘x’, ‘j’, and ‘u’ were selected for the identification of lncRNAs and NPCTs on the sorghum reference genome and the cultivar reference genome.

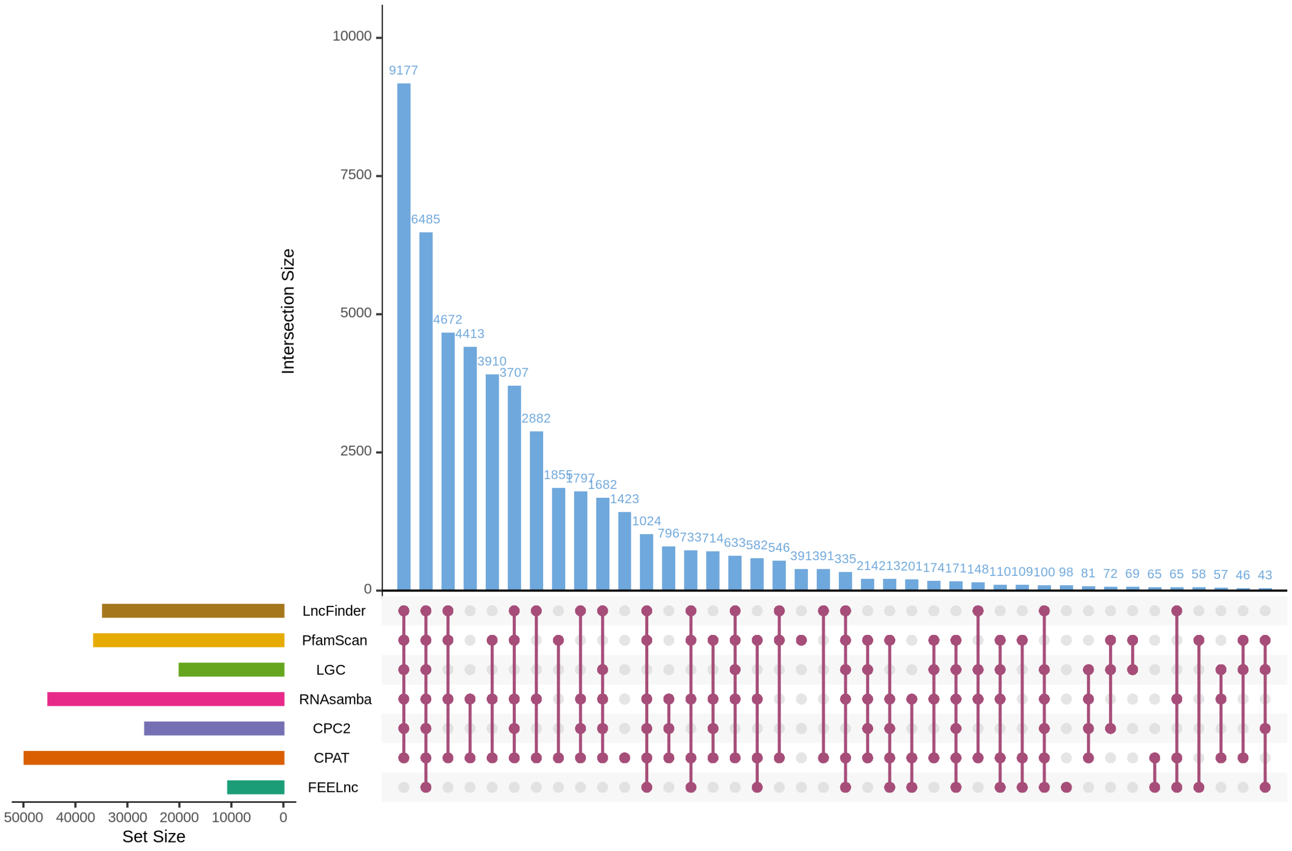


**Supplemental Figure S1(b)**: The 6485 true lncRNAs were identified by LncRAnalyzer in BTX642 using seven methods on the sorghum reference genome v3.1.

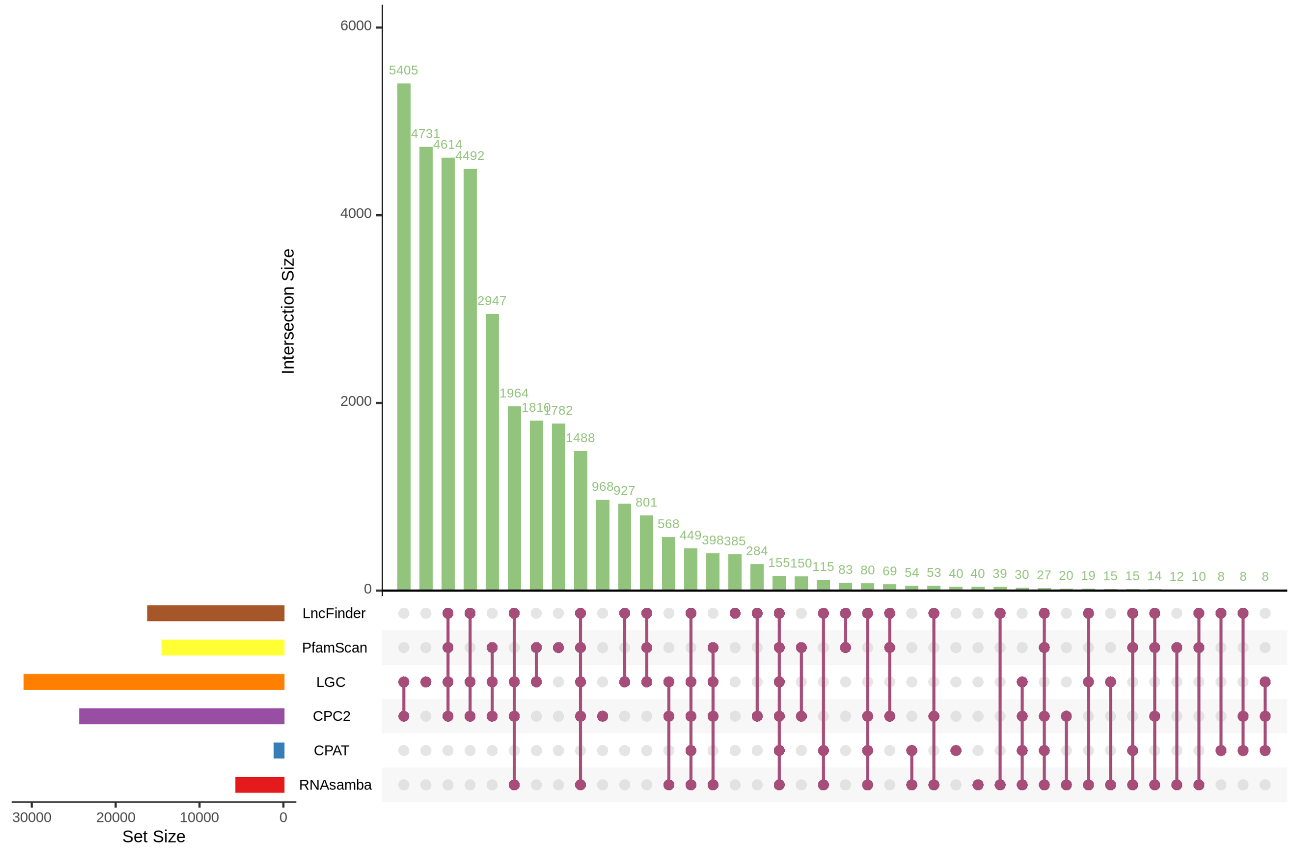


**Supplemental Figure S1(c)**: The 155 true NPCTs were identified by LncRAnalyzer in BTX642 using six methods on the sorghum reference genome v3.1.

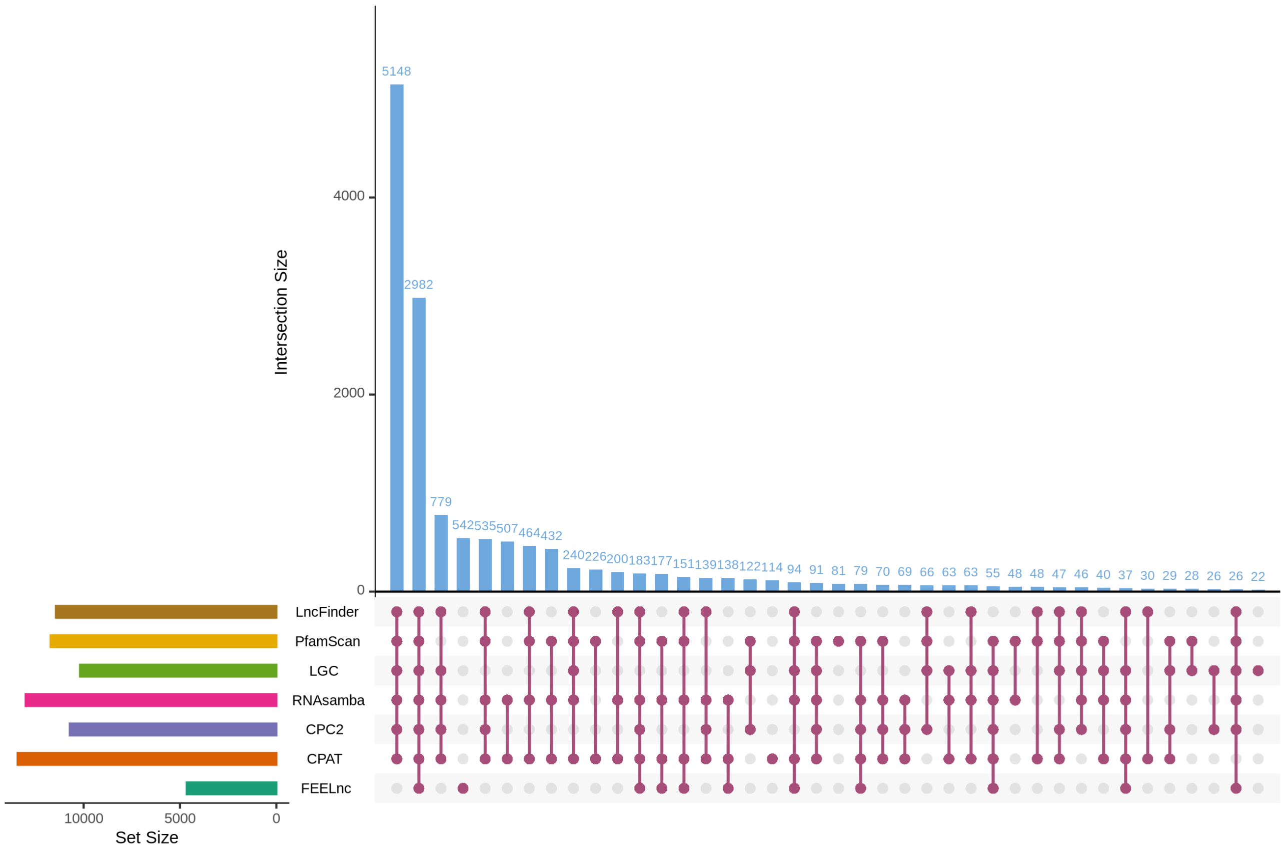


**Supplemental Figure S1(d)**: The 2982 true lncRNAs were identified by LncRAnalyzer in BTX642 using seven methods on the cultivar-specific reference genome.

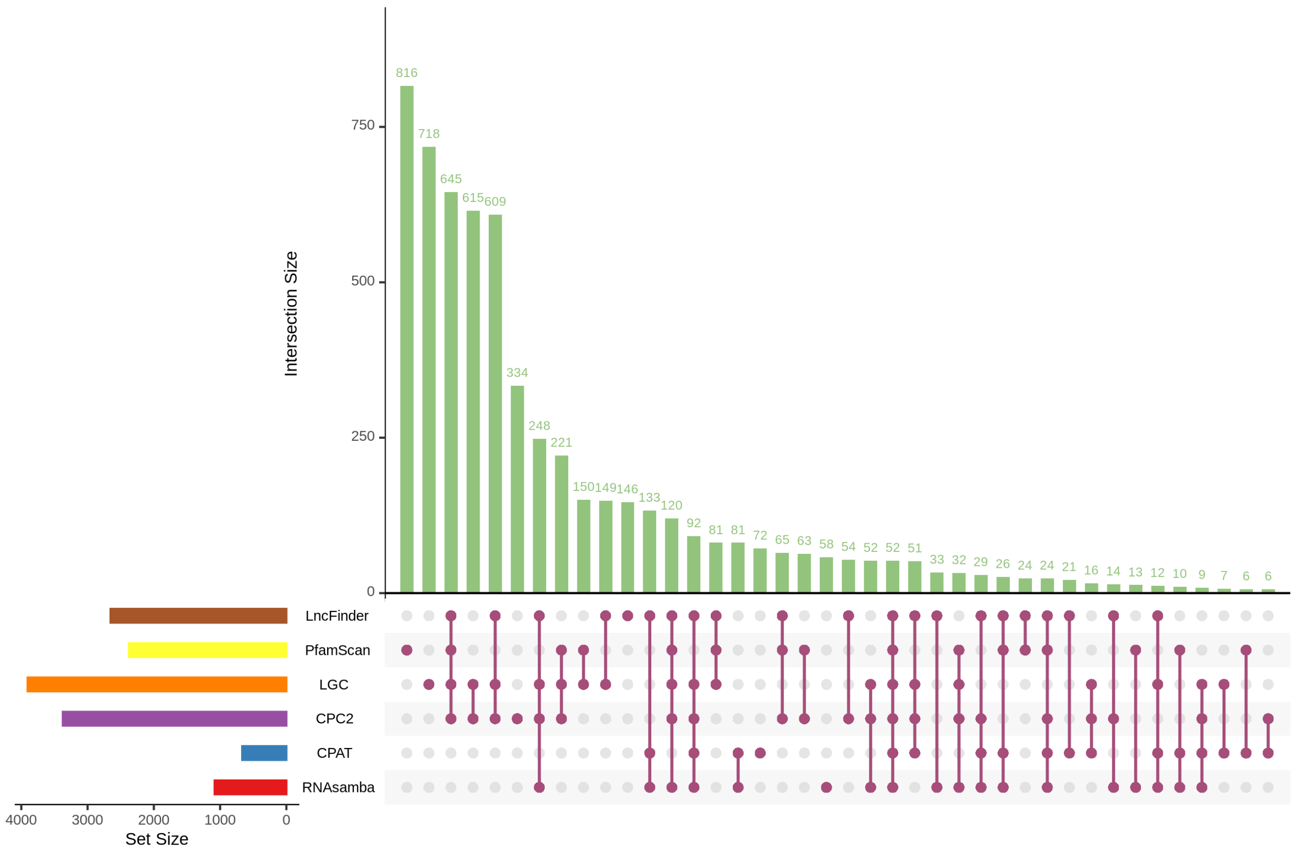


**Supplemental Figure S1(e)**: The 52 true NPCTs identified by LncRAnalyzer in BTX642 using six methods on the cultivar-specific reference genome.

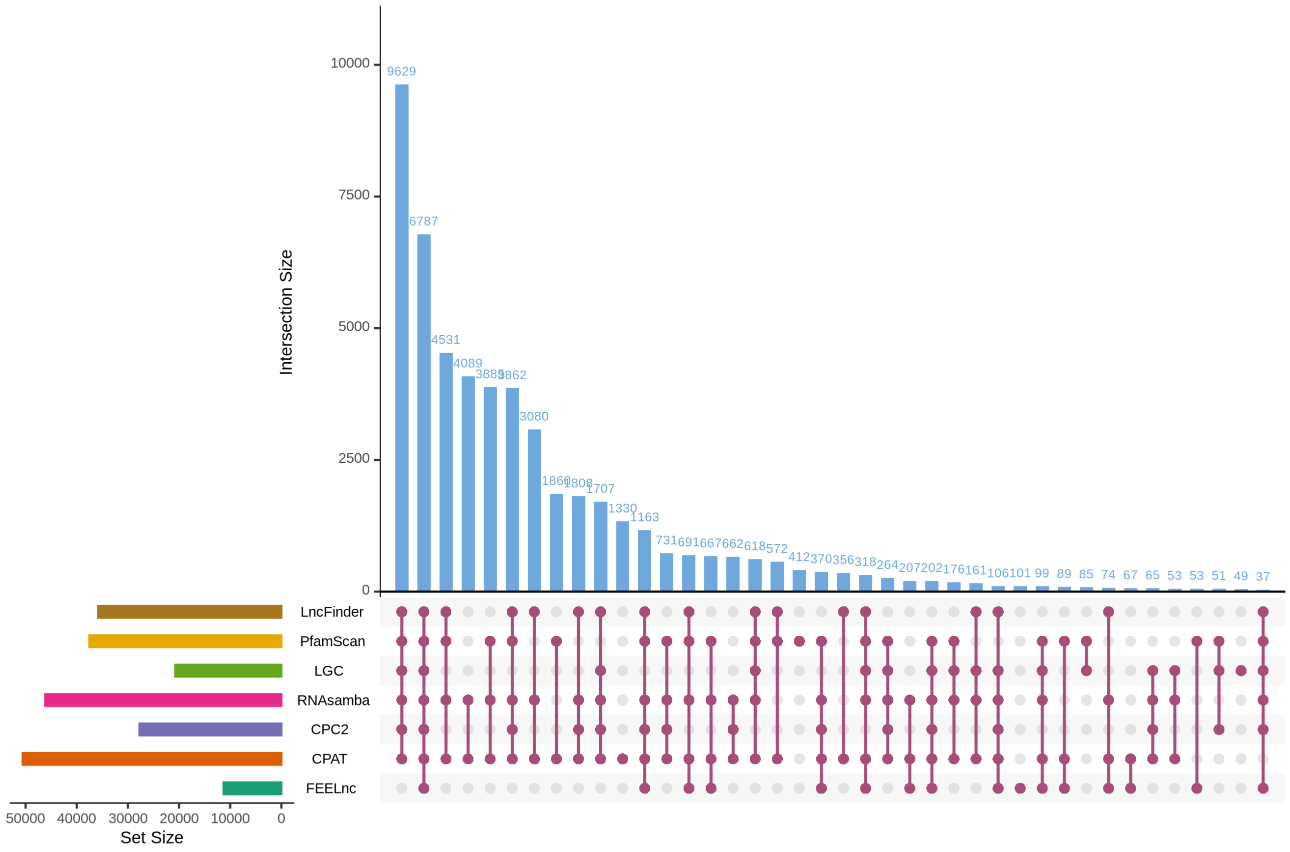


**Supplemental Figure S1(f)**: The 6787 true lncRNAs were identified by LncRAnalyzer in RTX430 using seven methods on the sorghum reference genome v3.1.

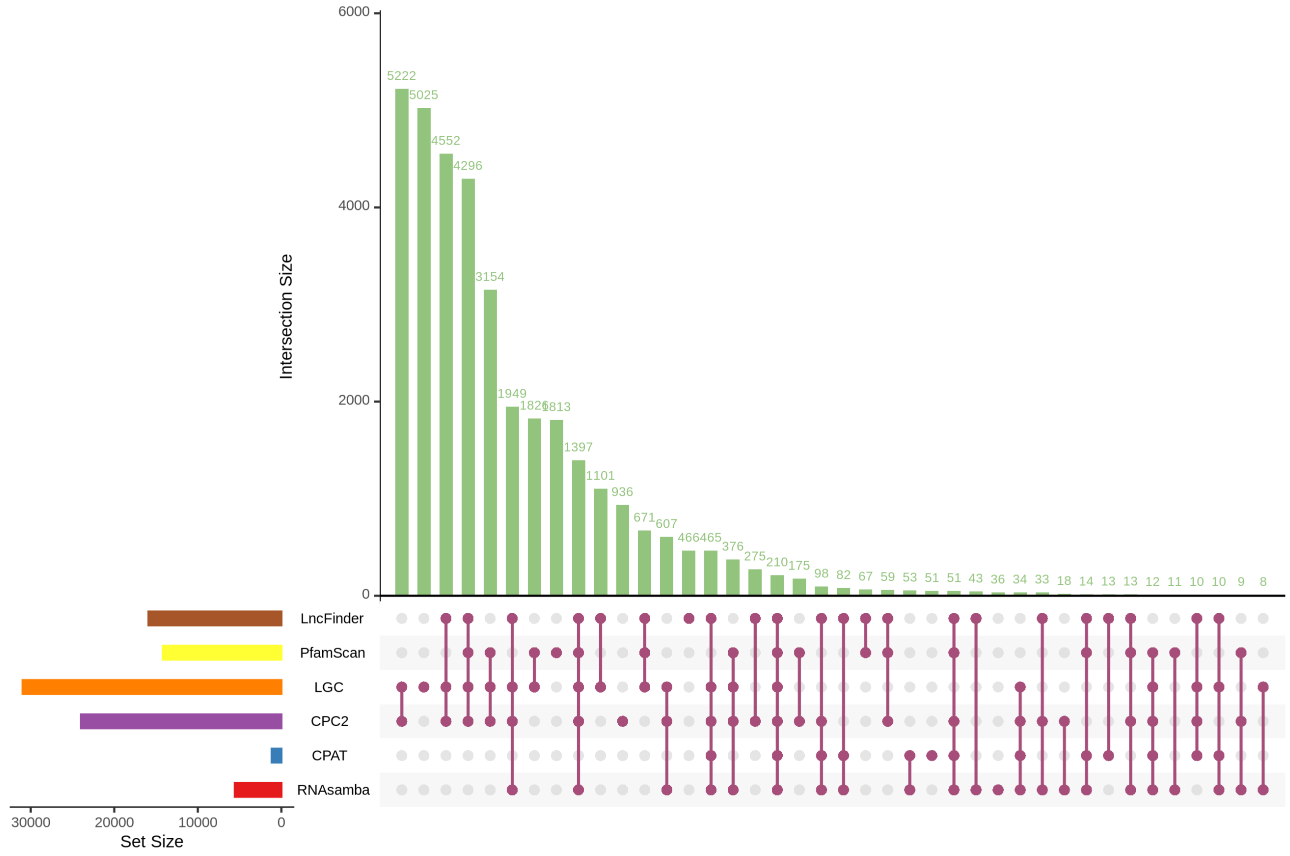


**Supplemental Figure S1(g)**: The 210 true NPCTs were identified by LncRAnalyzer in RTX430 using six methods on the sorghum reference genome v3.1.

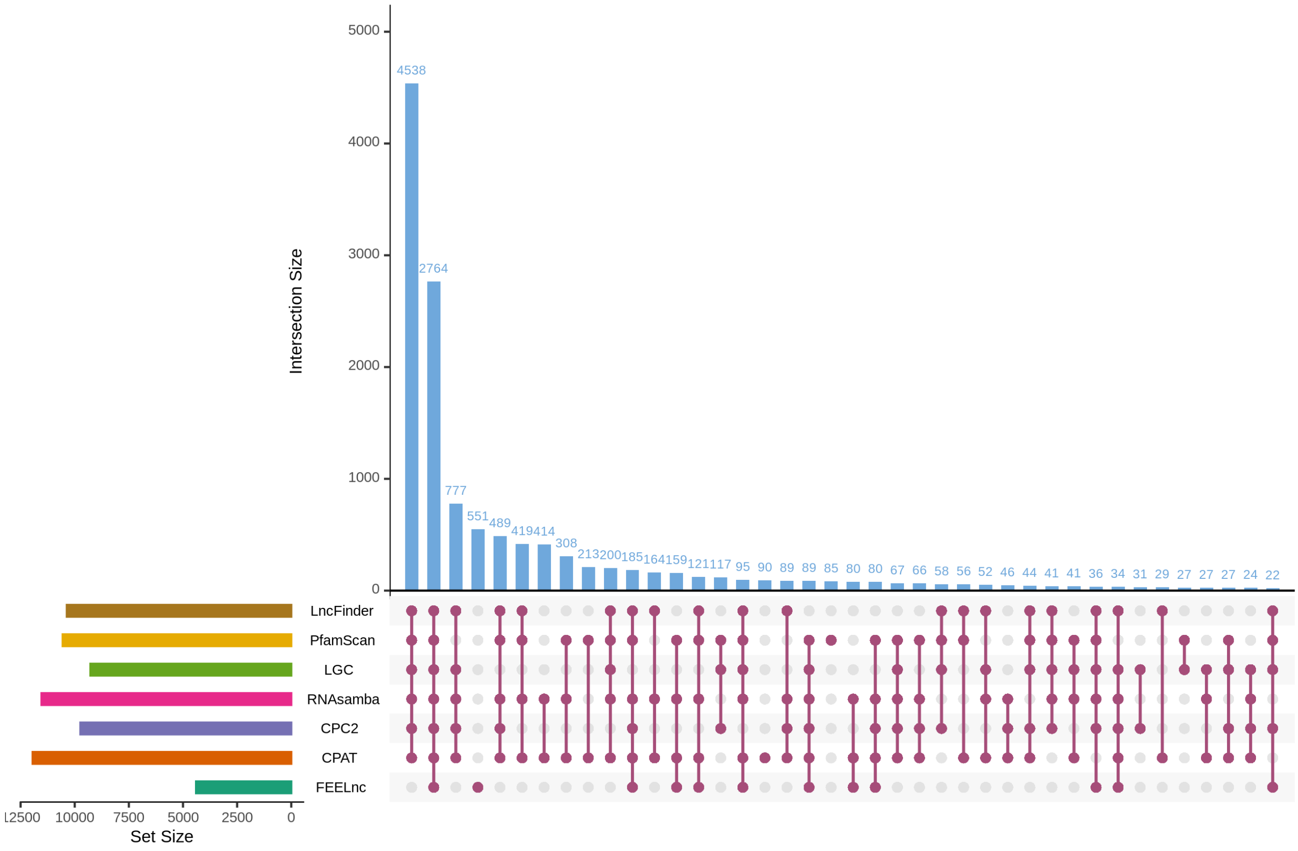


**Supplemental Figure S1(h)**: The 2764 true lncRNAs identified by LncRAnalyzer in RTX430 using seven methods on the cultivar-specific reference genome.

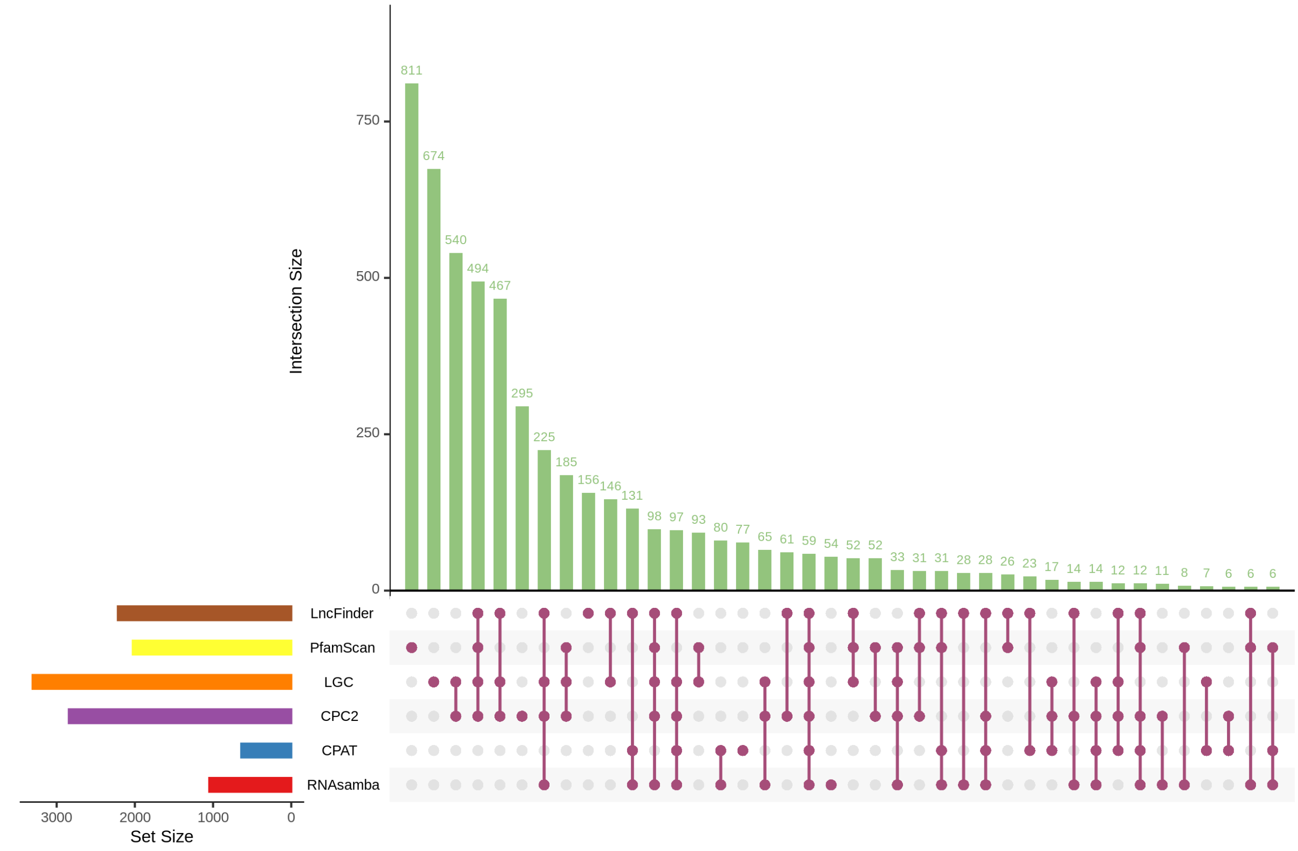


**Supplemental Figure S1(i)**: The 59 true NPCTs identified by LncRAnalyzer in RTX430 using six methods on the cultivar-specific reference genome.

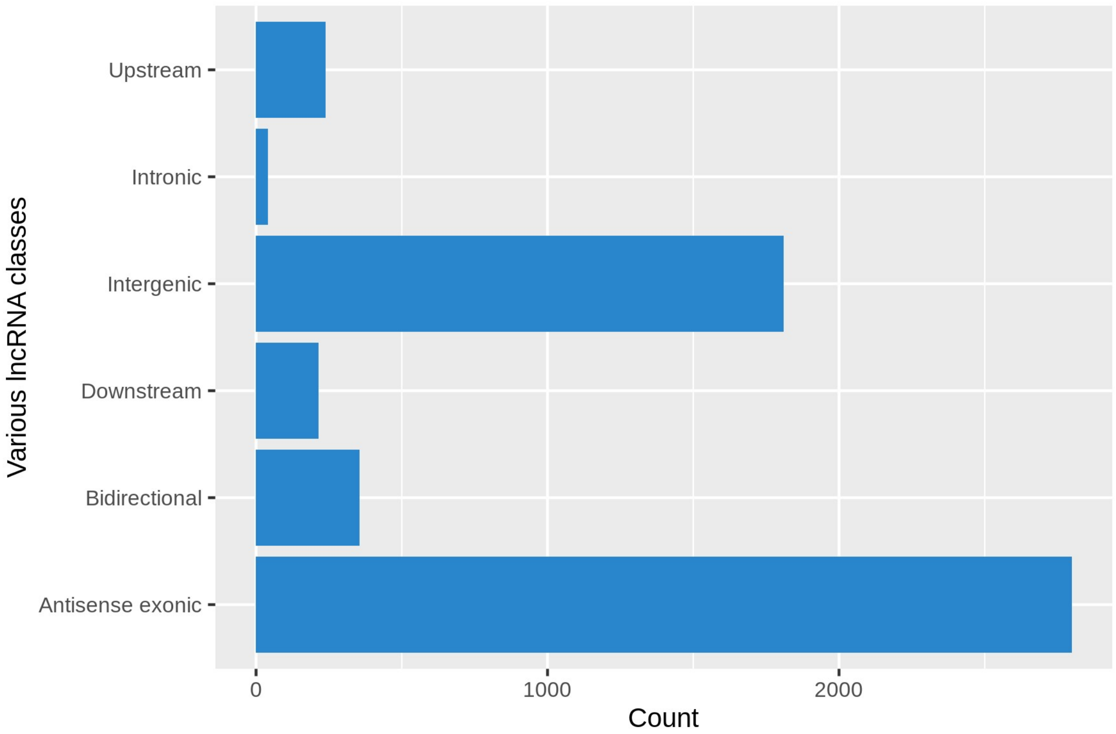


**Supplemental Figure S1(j)**: BTX642 identified 6485 true lncRNAs on the sorghum reference genome, of which 5457 (84.14%) were assigned classes based on their orientation, type, genomic locations, and distance from protein-coding genes.

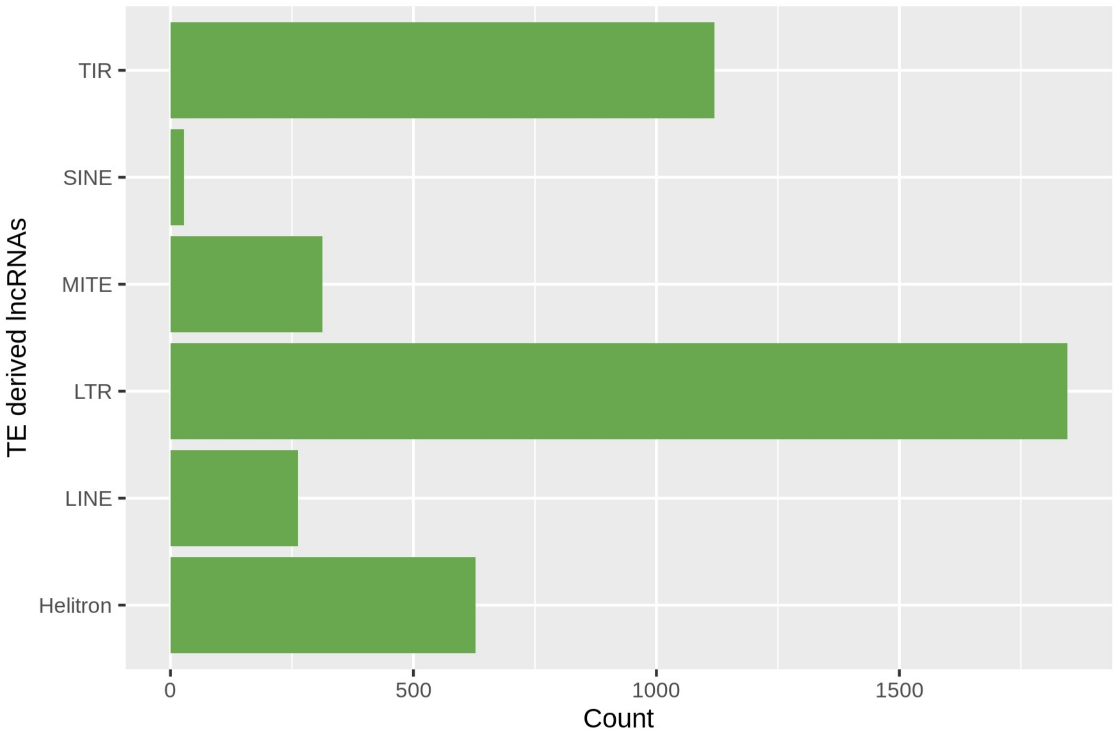


**Supplemental Figure S1(k)**: BTX642 identified 6485 true lncRNAs on the sorghum reference genome, of which 4198 (64.73%) were derived from various Transposable Elements (TEs).

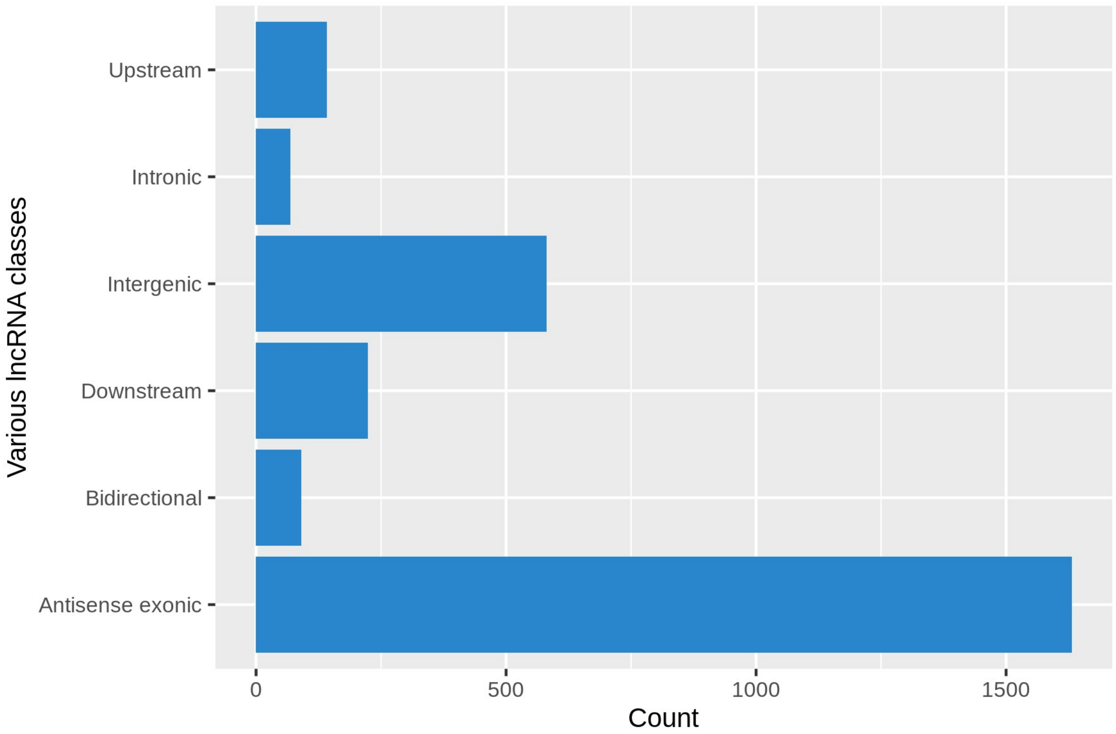


**Supplemental Figure S1(l)**: BTX642 identified 2982 true lncRNAs on cultivar-specific reference genome, of which 2734 (91.68%) were assigned classes based on their orientation, type, genomic locations, and distance from protein-coding genes.

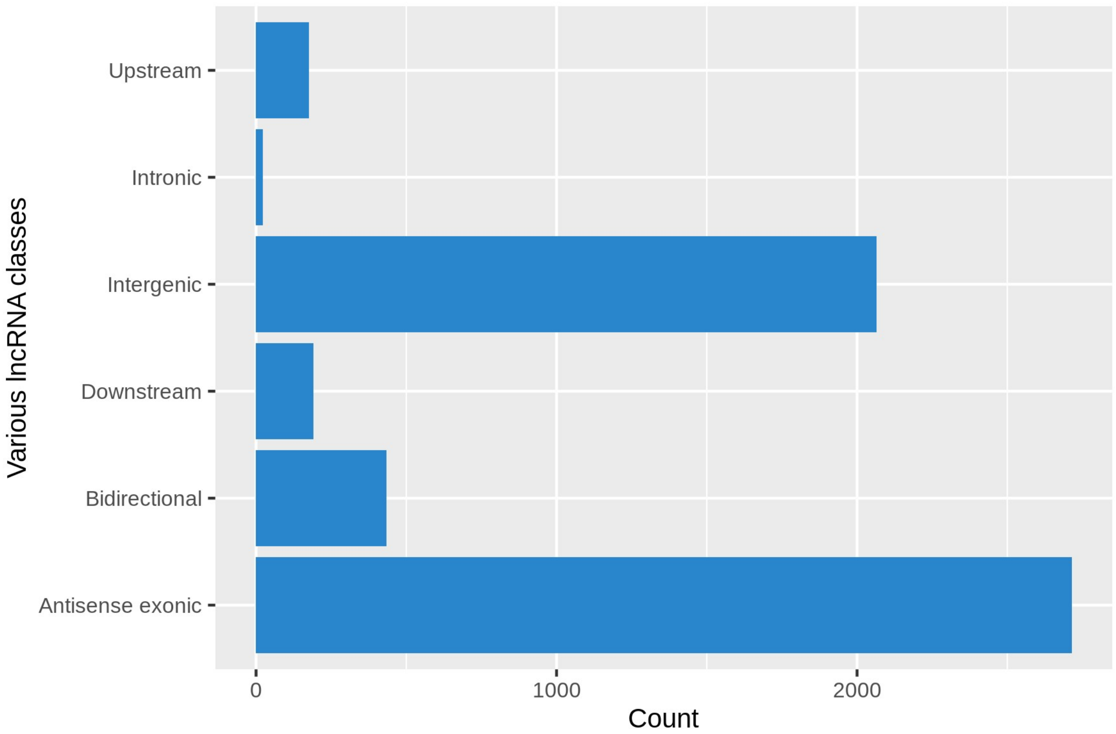


**Supplemental Figure S1(m)**: RTX430 identified 6787 true lncRNAs on the sorghum reference genome, of which 5603 (82.55%) were assigned classes based on their orientation, type, genomic locations, and distance from protein-coding genes.

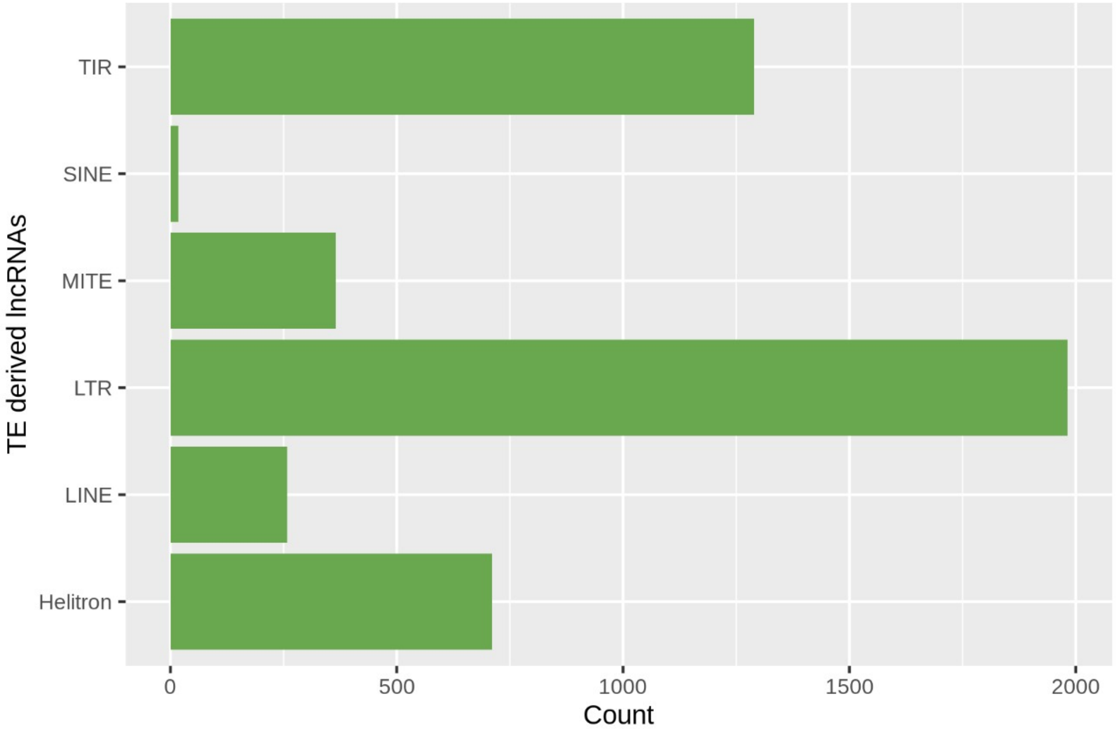


**Supplemental Figure S1(n)**: RTX430 identified 6787 true lncRNAs on the sorghum reference genome, of which 4624 (68.13%) were derived from various Transposable Elements (TEs).

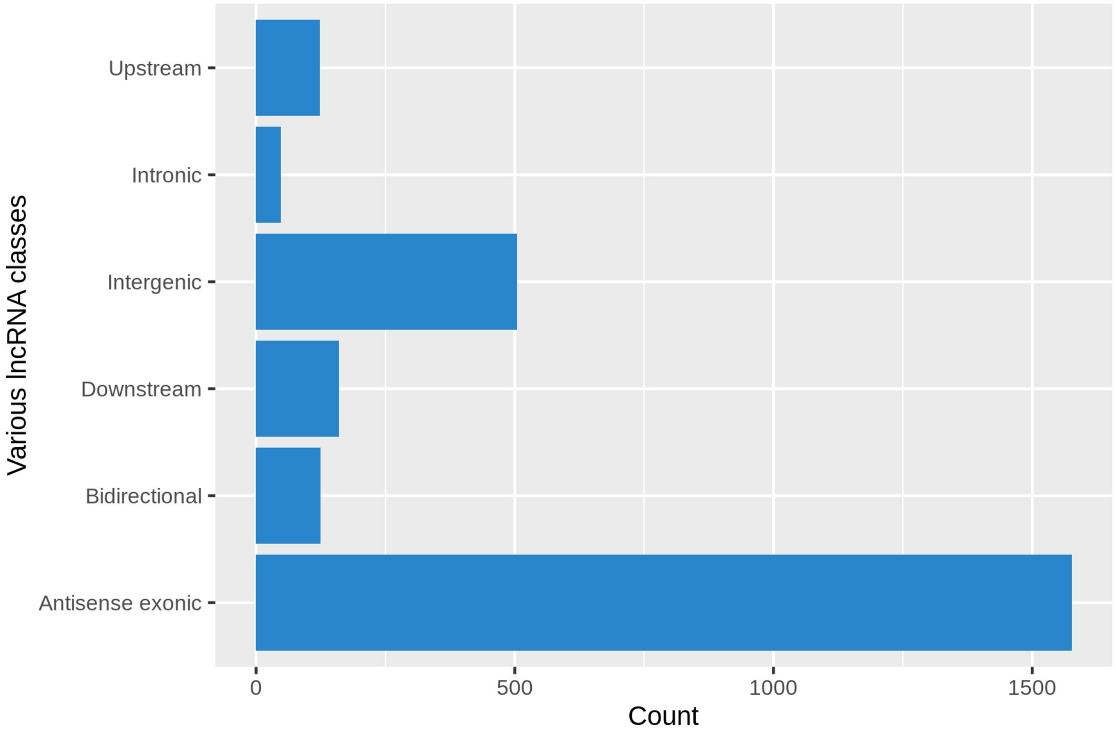


**Supplemental Figure S1(o)**: RTX430 identified 2764 true lncRNAs on cultivar-specific reference genome, of which 2537 (91.78%) were assigned classes based on their orientation, type, genomic locations, and distance from protein-coding genes.

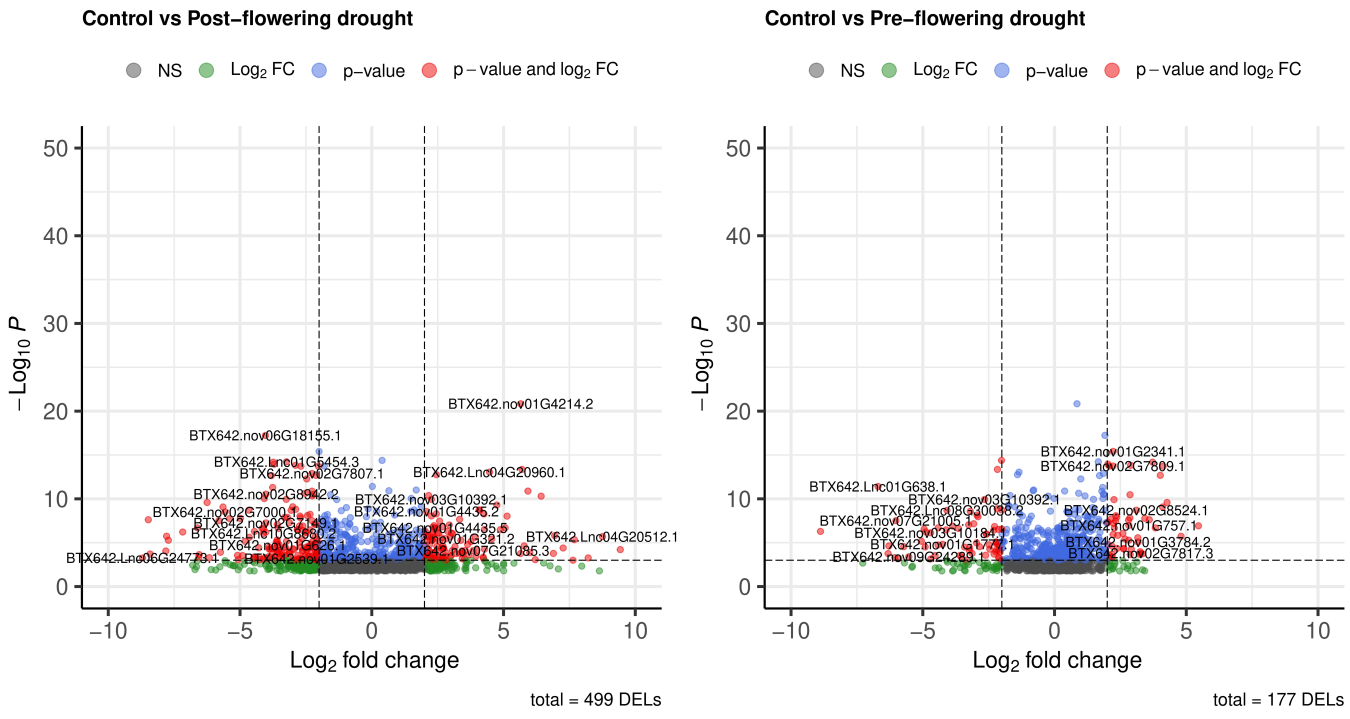


**Supplemental Figure S4(a)**: Number of DELs with P-value < 0.05 and log2fold > |2| in BTX642 leaf tissues under control, pre-, and post-flowering drought conditions.

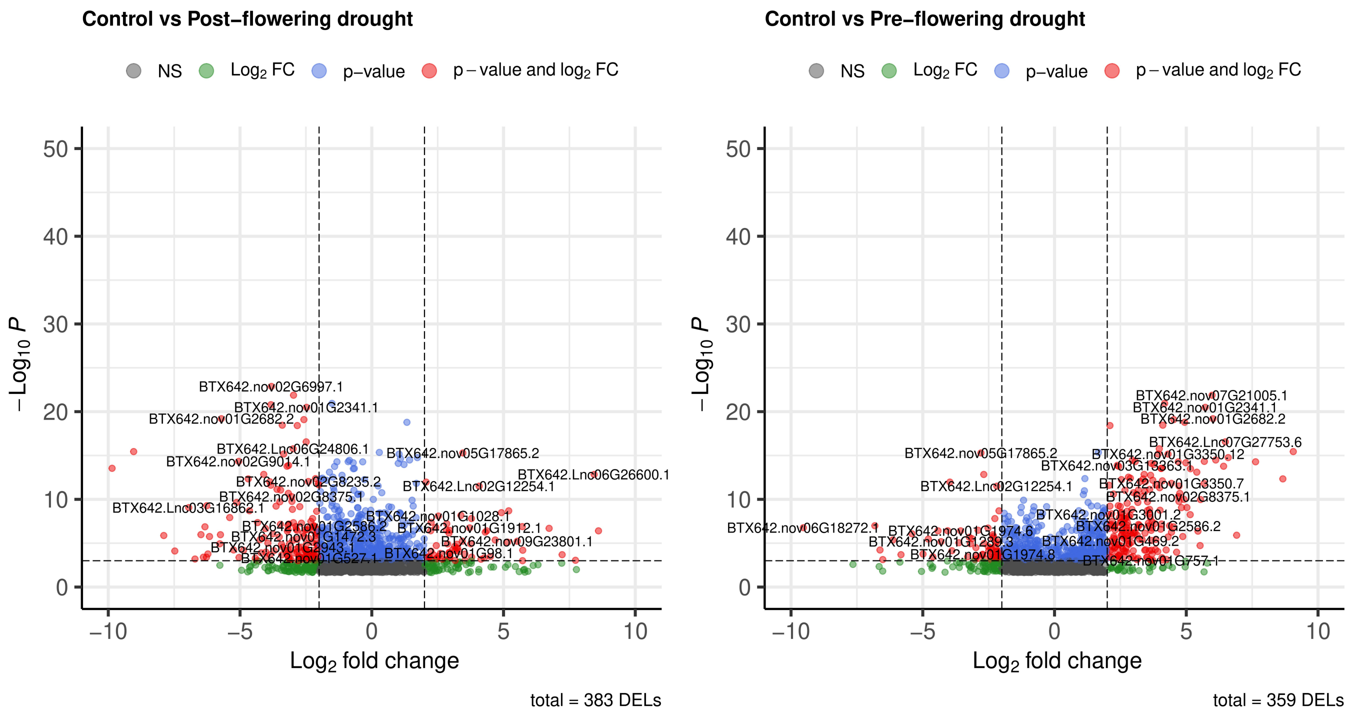


**Supplemental Figure S4(b)**: Number of DELs with P-value < 0.05 and log2fold > |2| in BTX642 root tissues under control, pre-, and post-flowering drought conditions.

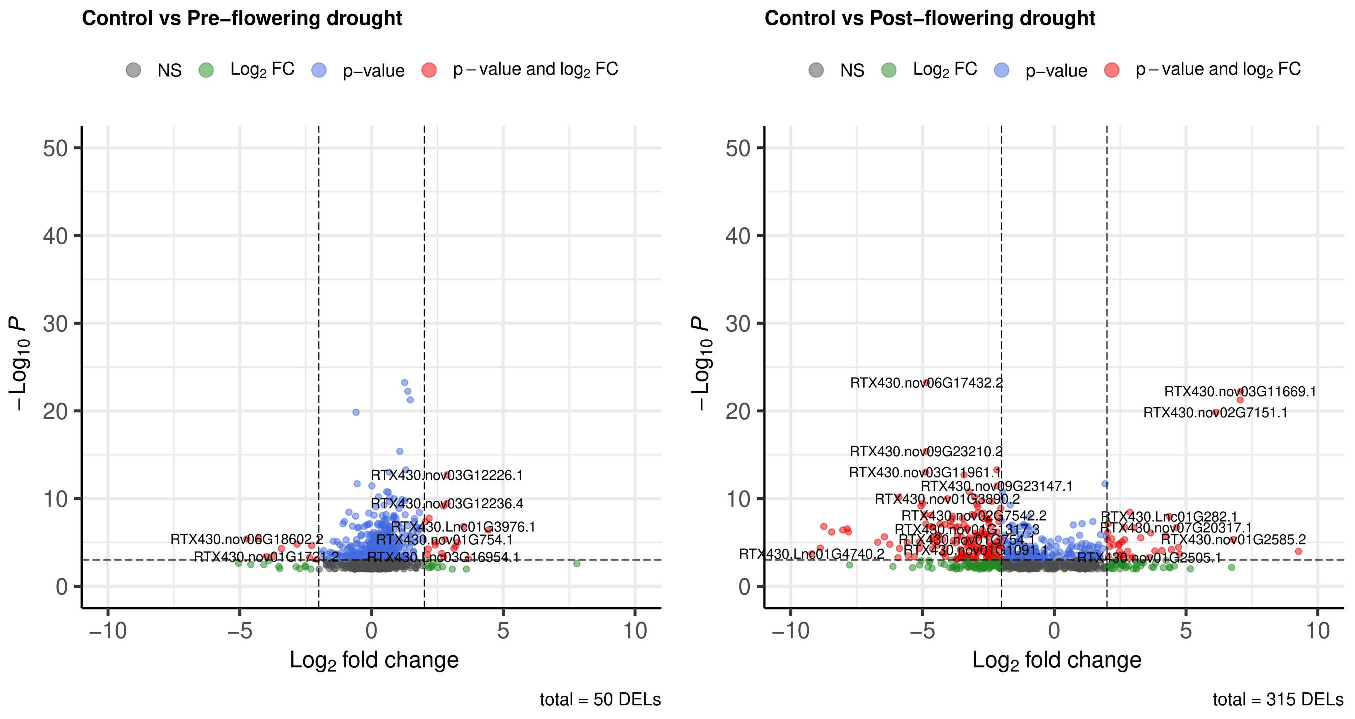


**Supplemental Figure S4(c)**: Number of DELs with P-value < 0.05 and log2fold > |2| in RTX430 leaf tissues under control, pre-, and post-flowering drought conditions.

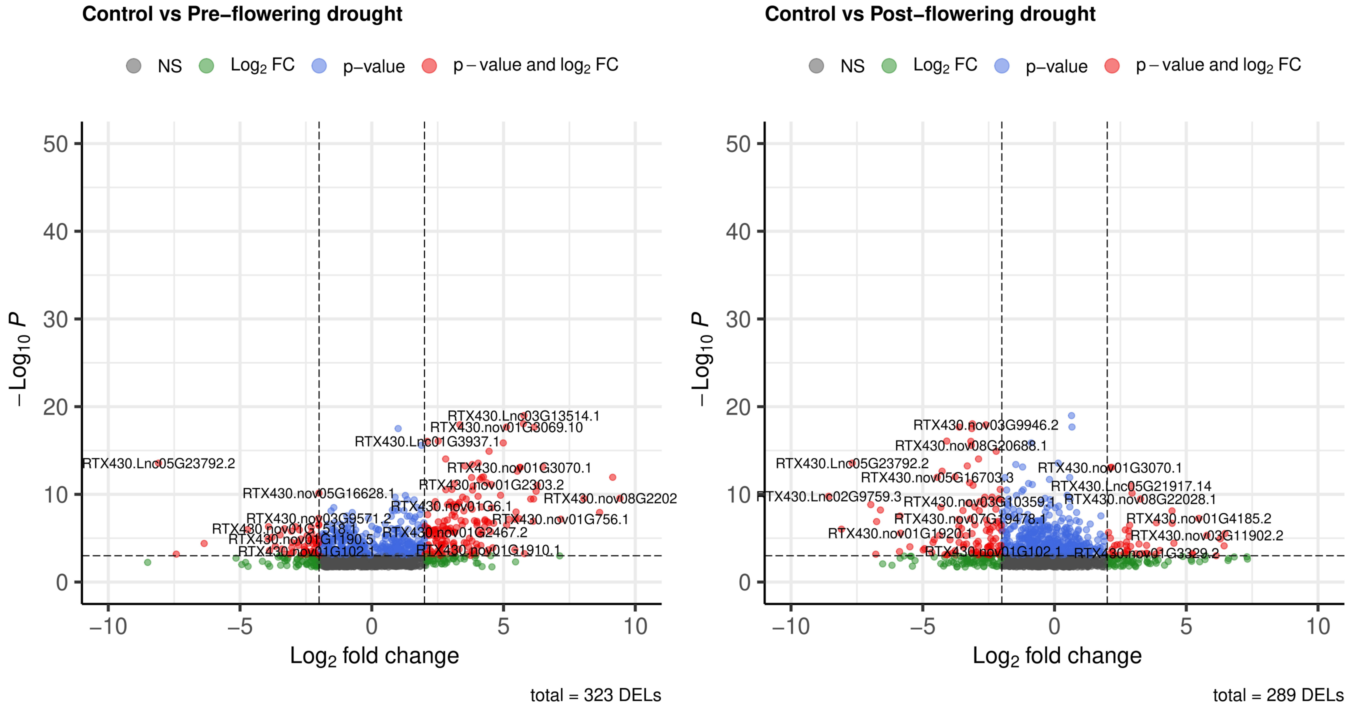


**Supplemental Figure S4(d):** Number of DELs with P-value < 0.05 and log2fold > |2| in RTX430 root tissues under control, pre-, and post-flowering drought conditions.

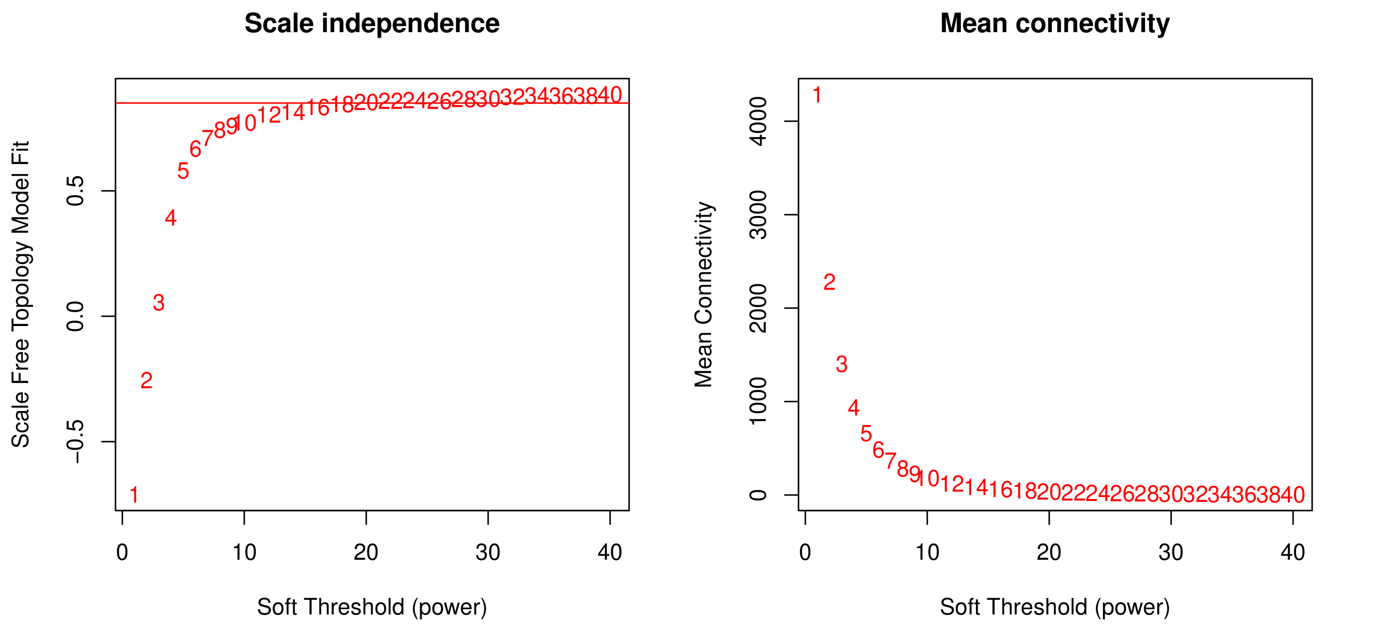


**Supplemental Figure S6(a):** WGCNA scale independence and mean connectivity analysis were performed to get the soft threshold (powers) ($R^2 > 0.85$) for BTX642 leaf tissues.

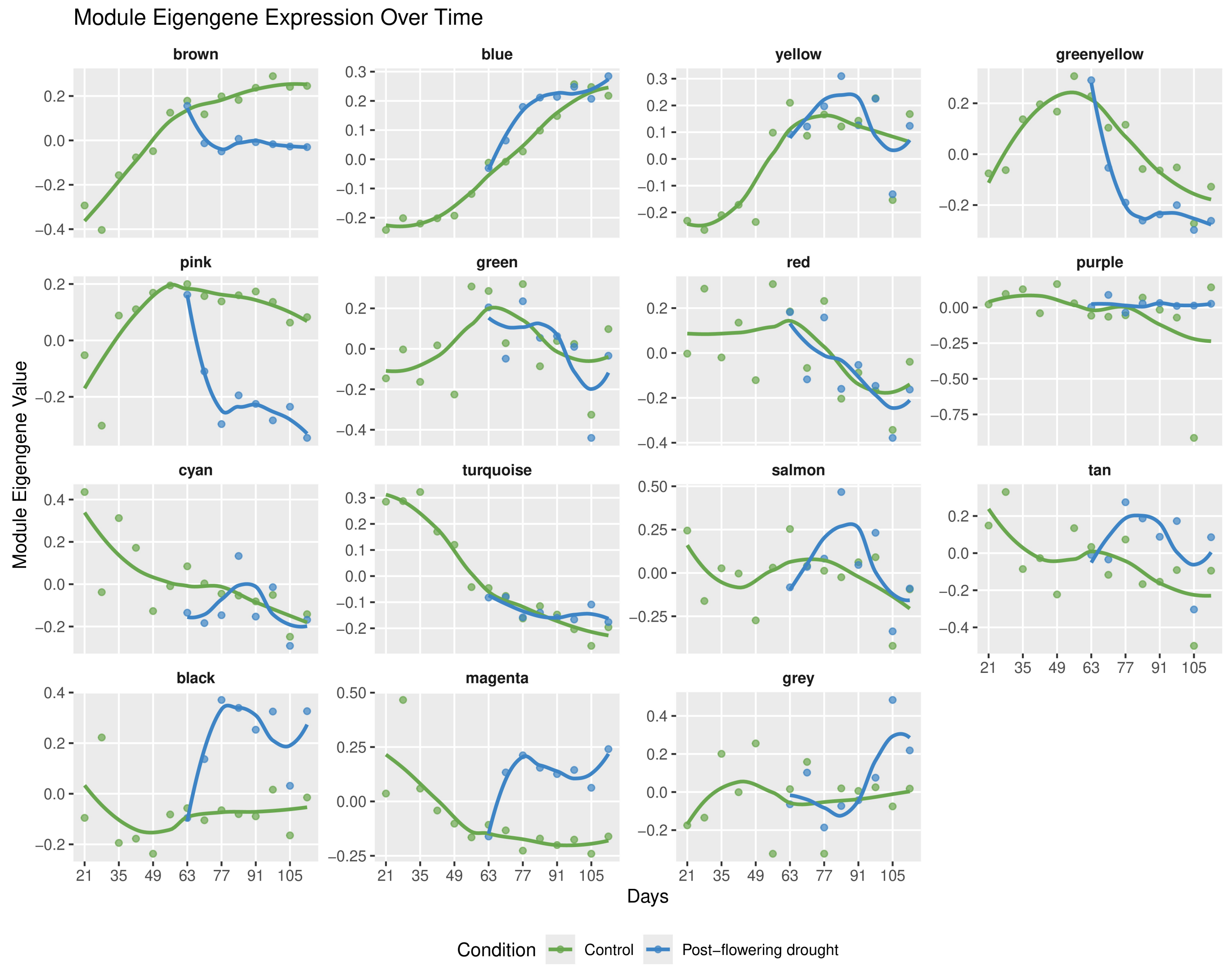


**Supplemental Figure S6(b):** WGCNA module eigengene analysis was performed to capture the overall expression trends of the different coexpression modules in BTX642 leaf tissues.

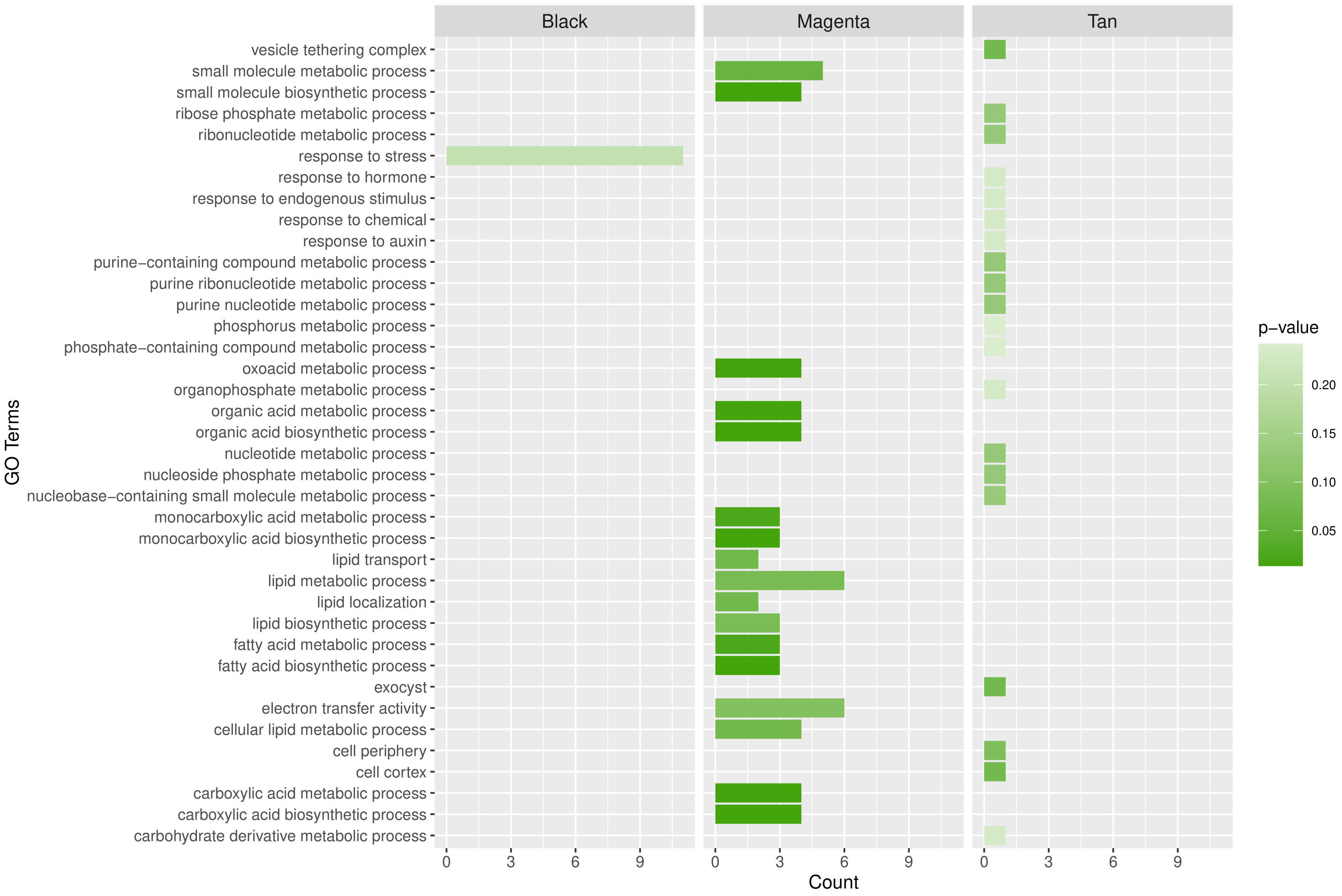


**Supplemental Figure S6(c):** Gene ontology enrichment analysis of “Black”, “Magenta”, and “Tan” coexpression modules in BTX642 leaf tissues, which were associated with post-flowering drought tolerance.

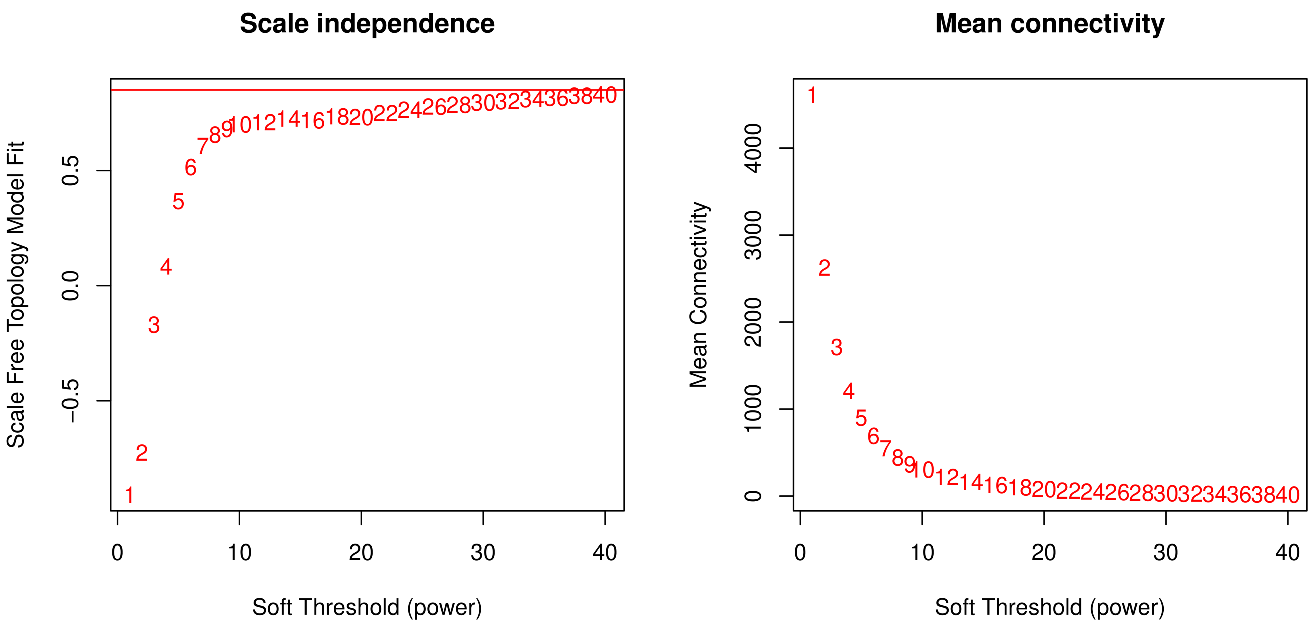


**Supplemental Figure S6(d):** WGCNA scale independence and mean connectivity analysis were performed to get the soft threshold (powers) ($R^2 > 0.85$) for BTX642 root tissues.

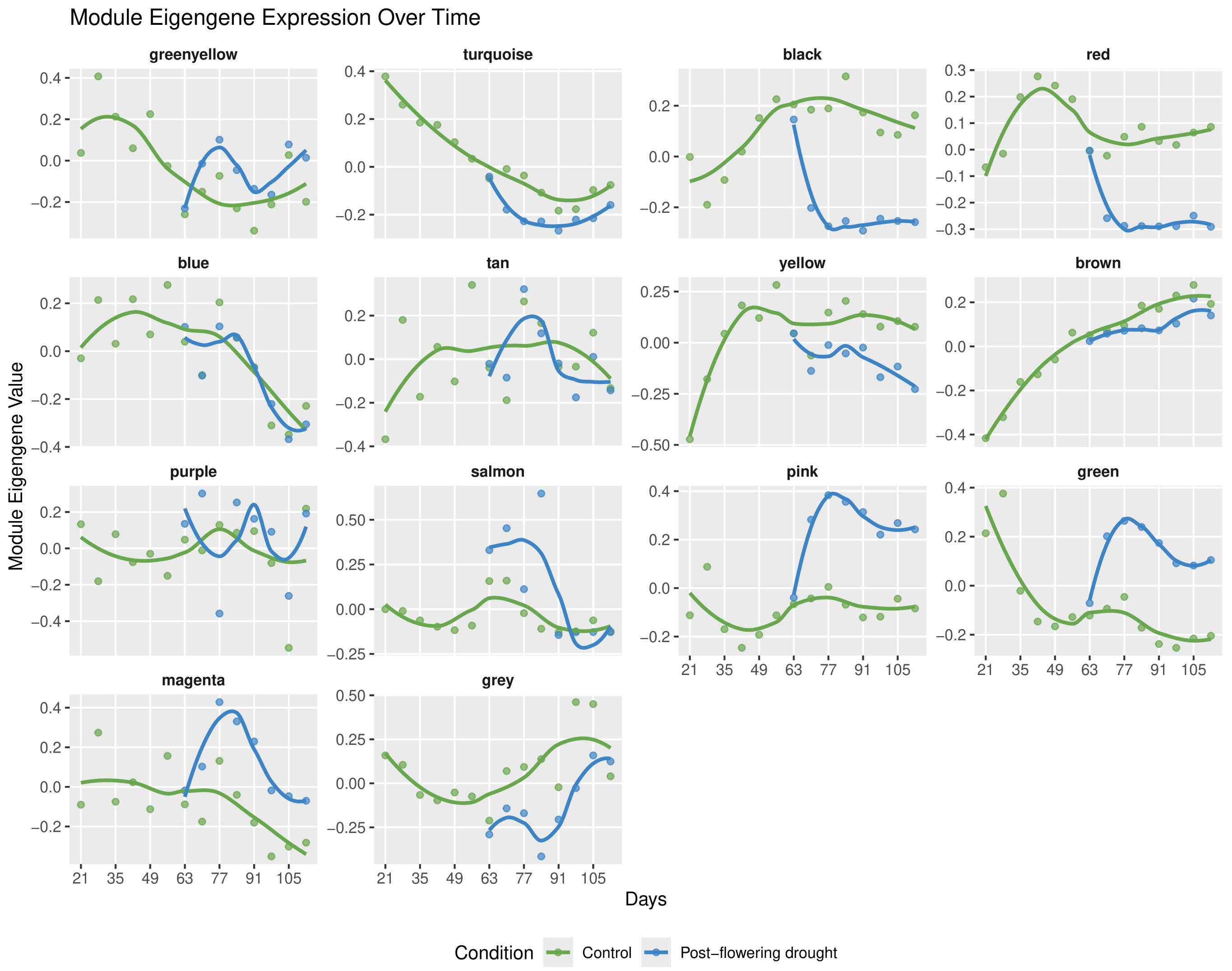


**Supplemental Figure S6(e):** WGCNA module eigengene analysis was performed to capture the overall expression trends of the different coexpression modules in BTX642 root tissues.

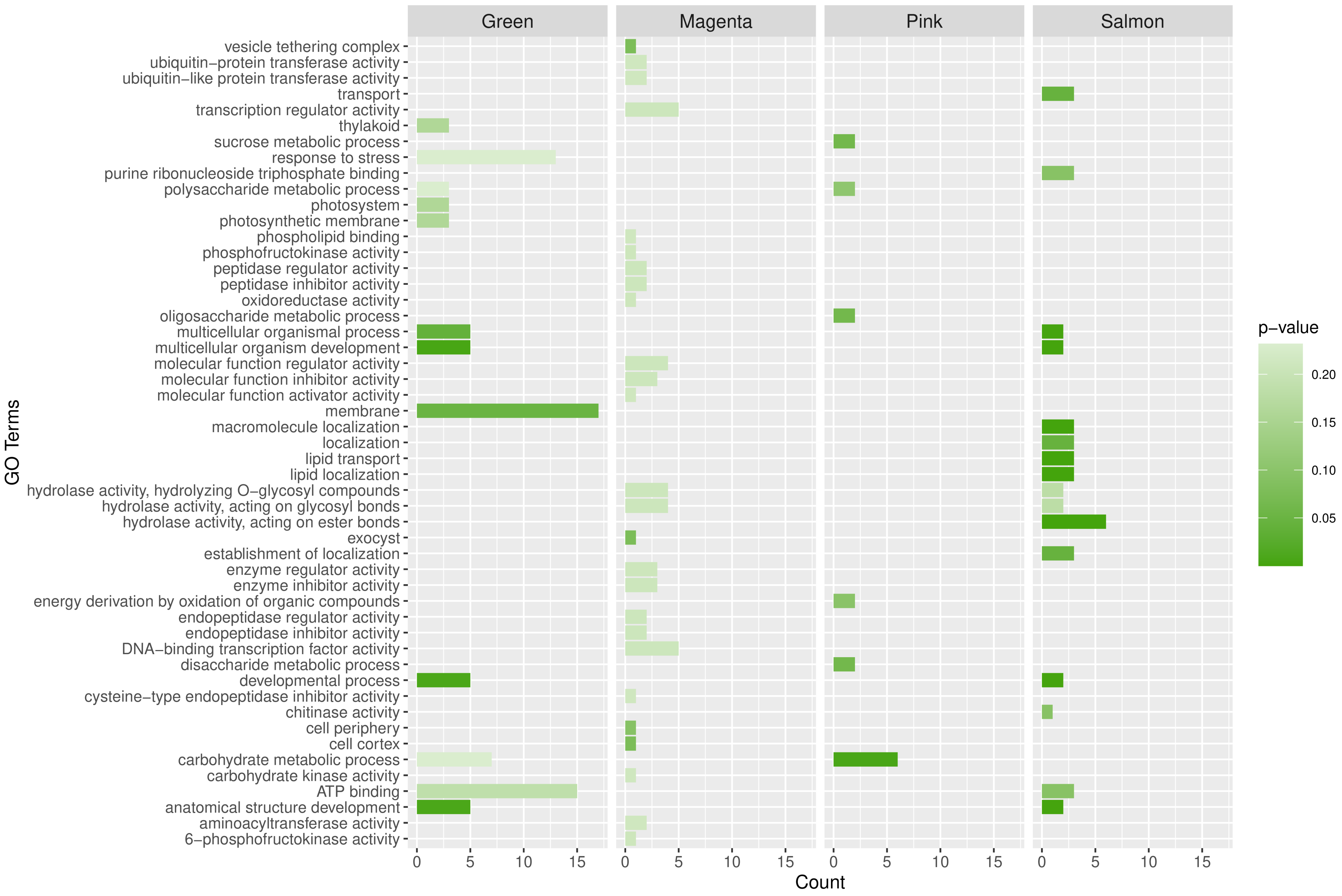


**Supplemental Figure S6(f):** Gene ontology enrichment analysis of “Green”, “Magenta”, “Pink”, and “Tan” coexpression modules in BTX642 root tissues, which were associated with post-flowering drought tolerance.

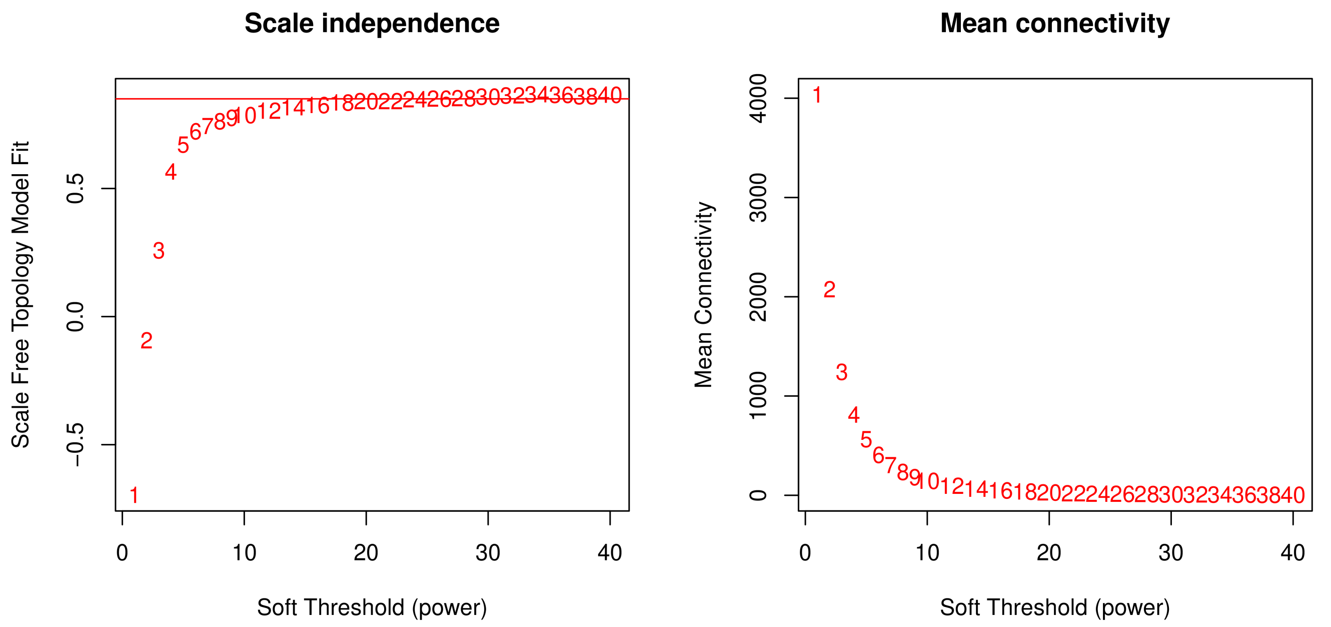


**Supplemental Figure S6(g):** WGCNA scale independence and mean connectivity analysis were performed to get the soft threshold (powers) ($R^2 > 0.85$) for RTX430 leaf tissues.

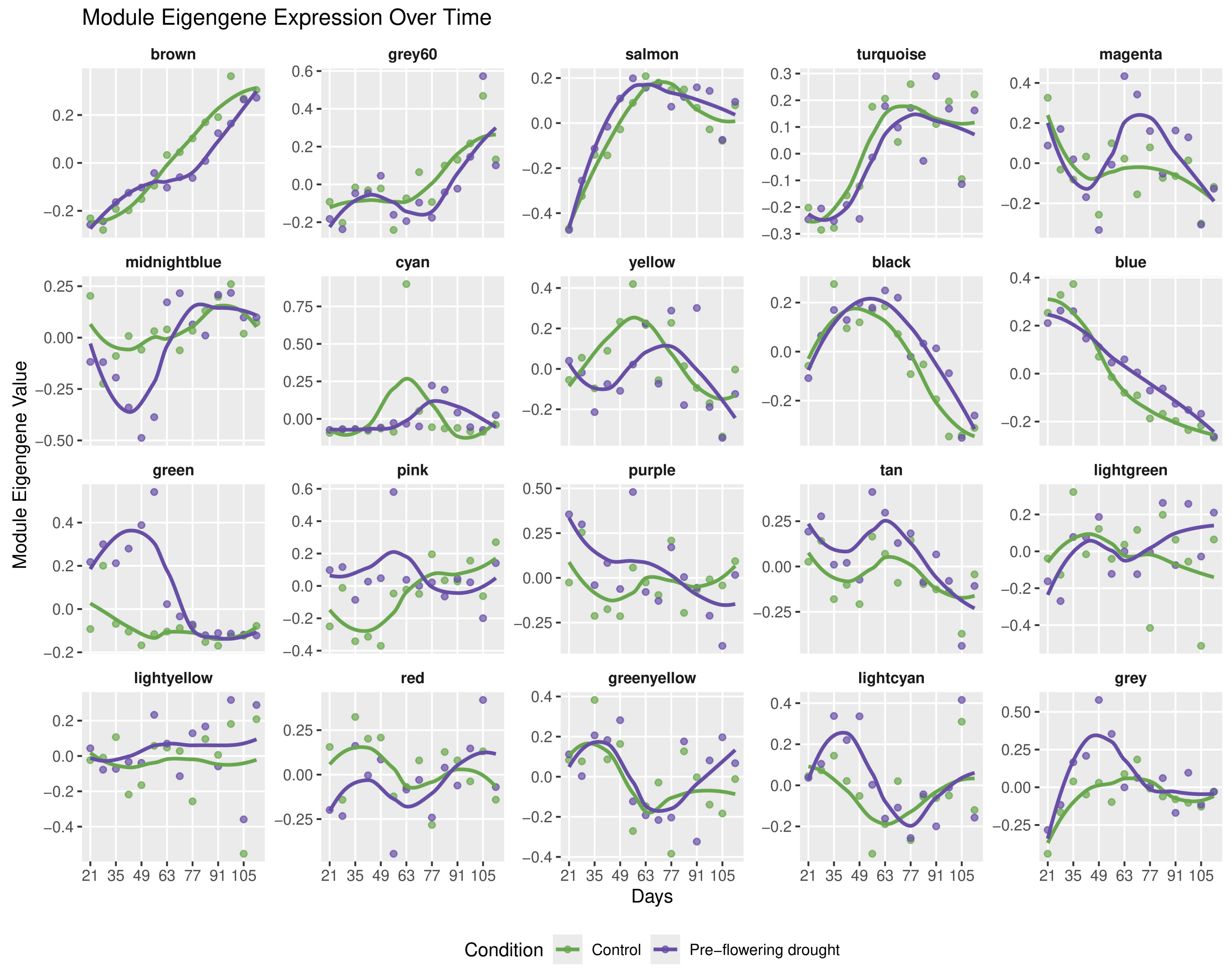


**Supplemental Figure S6(h):** WGCNA module eigengene analysis was performed to capture the overall expression trends of the different coexpression modules in RTX430 leaf tissues.

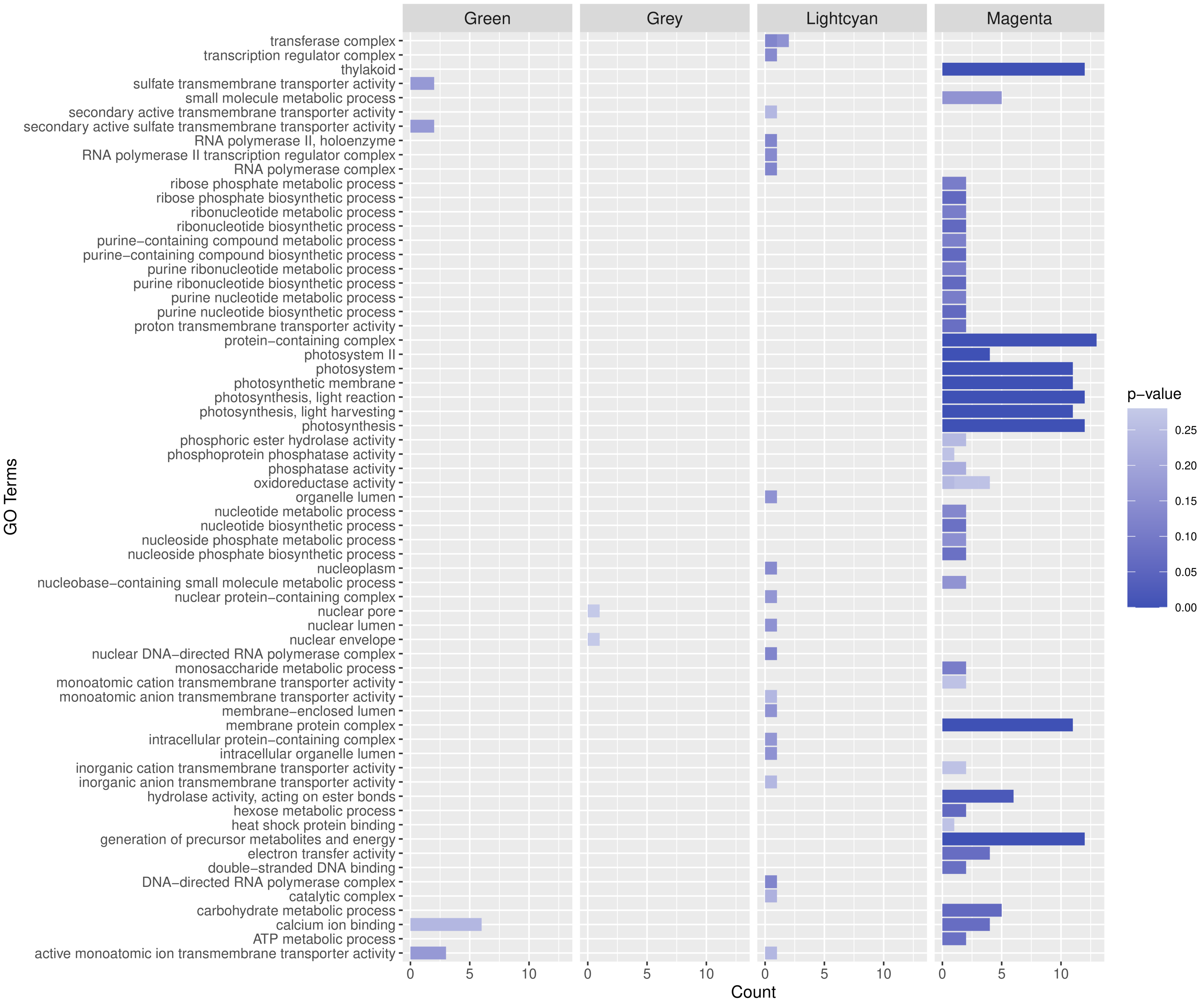


**Supplemental Figure S6(i):** Gene ontology enrichment analysis of “Green”, “Grey”, “Lightcyan”, and “Magenta” coexpression modules in RTX430 leaf tissues, which were associated with pre-flowering drought tolerance.

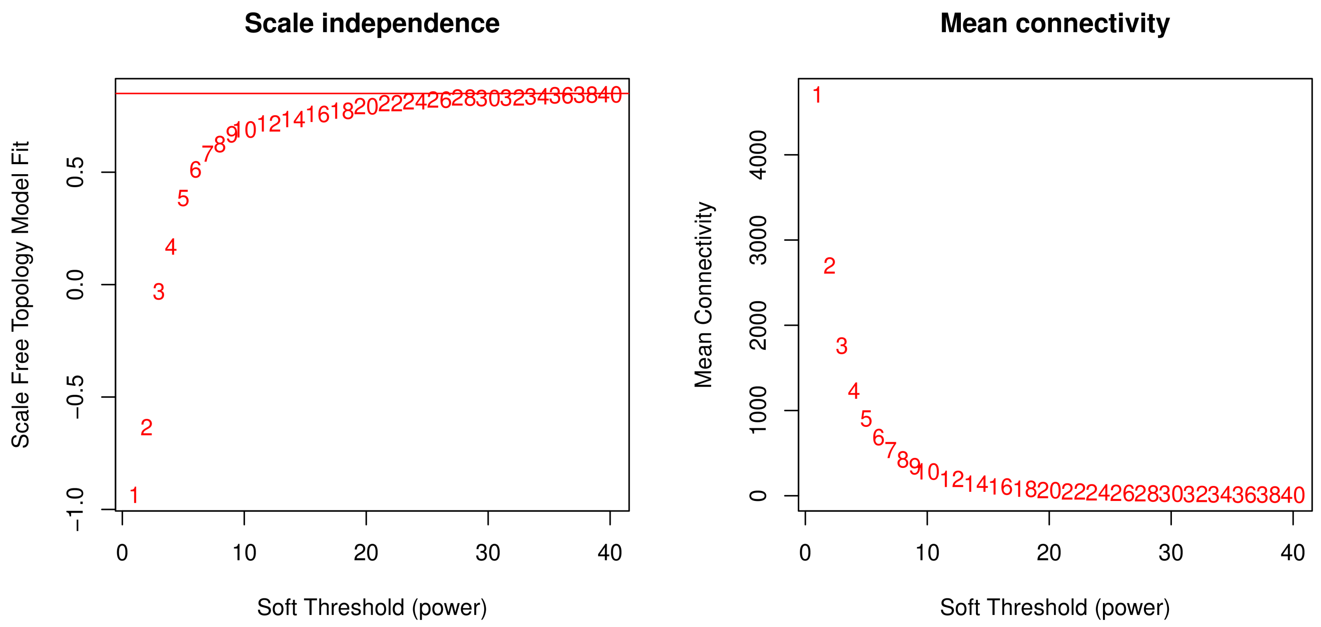


**Supplemental Figure S6(j):** WGCNA scale independence and mean connectivity analysis were performed to get the soft threshold (powers) ($R^2 > 0.85$) for BTX642 leaf tissues.

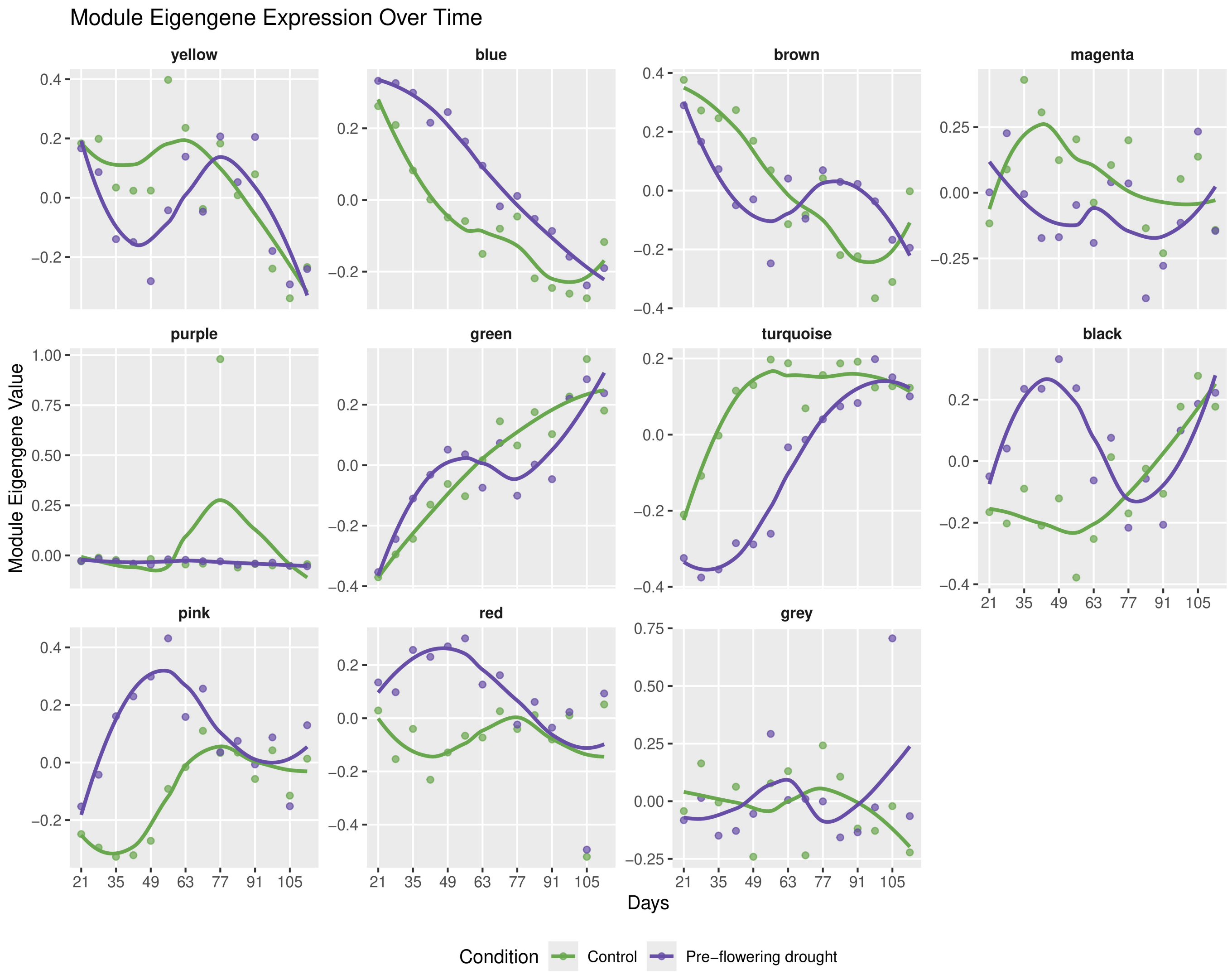


**Supplemental Figure S6(k):** WGCNA module eigengene analysis was performed to capture the overall expression trends of the different coexpression modules in RTX430 root tissues.

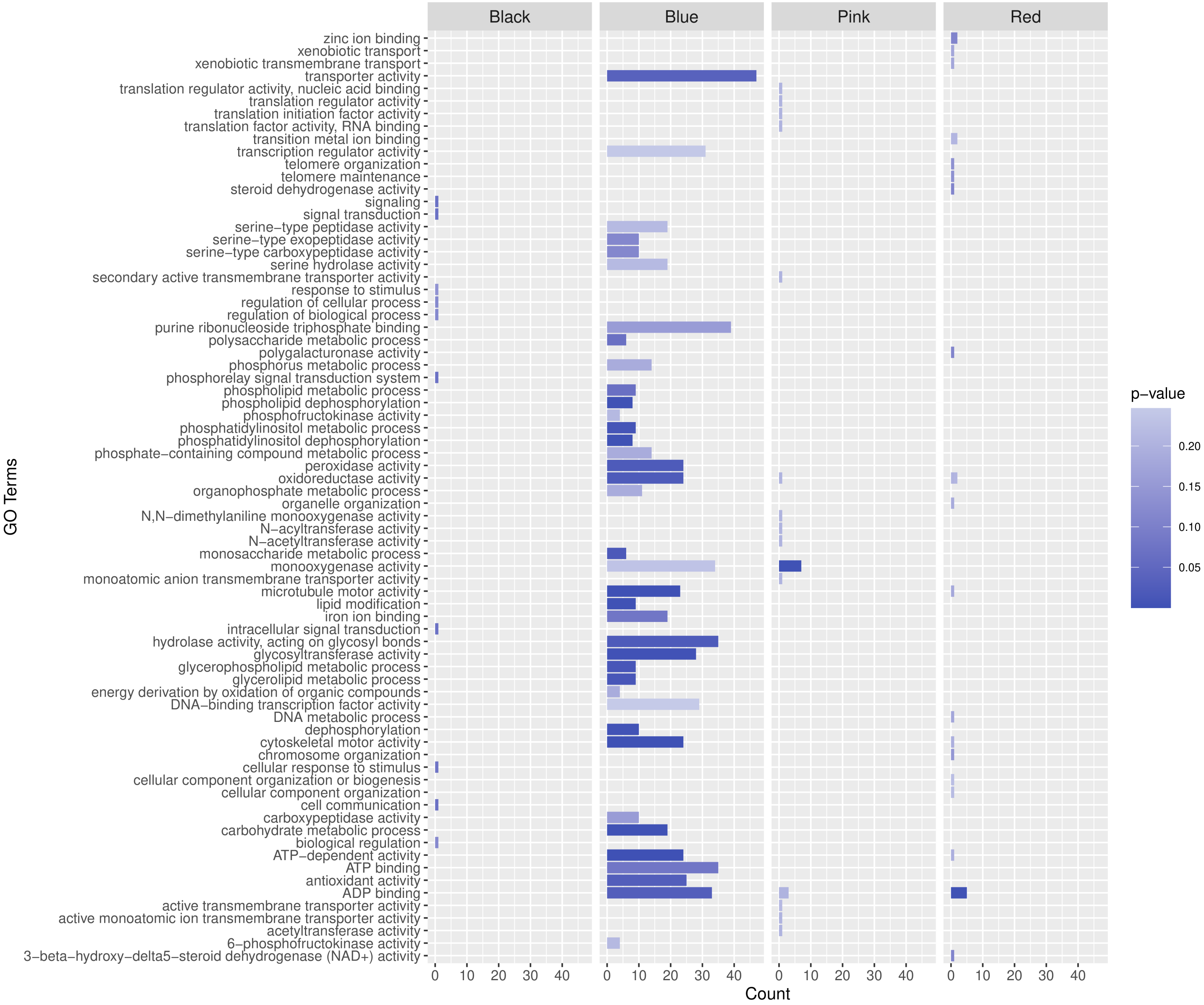


**Supplemental Figure S6(l):** Gene ontology enrichment analysis of “Black”, “Blue”, “Pink”, and “Red” coexpression modules in RTX430 root tissues, which were associated with pre-flowering drought tolerance.
